# E2F4 gene therapy reverses established synaptic and cognitive deficits and remodels learning-associated signaling in Alzheimer’s disease mice

**DOI:** 10.64898/2026.11.13.766444

**Authors:** Cristina Sánchez-Puelles, Alicia Hernández-Vivanco, Alice Grieco, Irene Camacho-Olmos, Ana Montero-Calle, Noelia López-Sánchez, Alberto Fraj-Cebrián, Alberto Garrido-García, Vanesa Cano-Daganzo, Anna Lozano-Ureña, Lorena Valdés-Lora, Rodrigo Barderas, Gertrudis Perea, Álvaro Pascual-Leone, José-María Frade

**Affiliations:** Department of Molecular and Cellular Neuroscience, Cajal Neurosciences Center, Consejo Superior de Investigaciones Científicas; Alcalá de Henares, Spain; Functional Proteomics Unit, Chronic Disease Programme, UFIEC, Instituto de Salud Carlos III, Carretera Pozuelo-Majadahonda Km 2.2, Majadahonda, 28220, Spain; Department of Computational and Systems Neuroscience, Cajal Neurosciences Center, Consejo Superior de Investigaciones Científicas; Alcalá de Henares, Spain; Hinda and Arthur Marcus Institute for Aging Research and Deanna, Sidney Wolk Center for Memory Health, Hebrew SeniorLife; Boston, MA, USA; Department of Neurology, Harvard Medical School; Boston, MA, USA; Tetraneuron; Madrid, Spain; CIBER Frailty and Healthy Aging, Madrid, E-28029 Madrid, Spain

**Keywords:** Alzheimer’s disease, dementia, synaptic dysfunction, cognitive decline, neuronal death, disease modification, gene therapy

## Abstract

Alzheimer’s disease (AD) causes progressive synaptic dysfunction, neurodegeneration, and cognitive decline, yet strategies capable of reversing established deficits remain limited. Here, we show that neuronal expression of a p38MAPK phosphorylation-resistant human E2F4 variant (hE2F4DN), delivered by adeno-associated virus, restores hippocampal long-term potentiation and memory in homozygous 5xFAD mice after functional impairment is established. Treatment also reduces amyloid-β and phospho-tau pathology and prevents hippocampal neuronal loss. Integrated proteomic and phosphoproteomic analyses reveal remodeling of metabolic, redox, neuroimmune, synaptic-vesicle, and GSK3β-associated signaling pathways. Following contextual fear conditioning, hE2F4DN-treated mice exhibit broader learning-associated synaptic proteomic signatures and distinct phosphoregulatory responses involving synaptic machinery. These findings demonstrate that E2F4 gene therapy can reverse established synaptic and cognitive deficits while preserving neuronal integrity and reshaping molecular responses to learning, identifying E2F4 phosphorylation as a potential therapeutic target for advanced AD states.

## INTRODUCTION

Alzheimer’s disease (AD) is a progressive neurodegenerative disorder and the leading cause of dementia, accounting for approximately 60–70% of cases worldwide (World Health Organization, 2026). The increasing prevalence of AD, together with the limited efficacy of current therapies, represents a major unmet medical need. This limitation is particularly evident at advanced stages, when neuronal damage and functional loss are already established. Currently, 73% of therapeutic strategies in development are disease targeted therapies, aimed to slow disease progression (Cummings et al., 2026). Therefore, existing treatments may transiently improve symptoms or modestly slow disease progression, but they do not halt or reverse the disease, underscoring the need for therapies targeting the underlying mechanisms driving neurodegeneration.

From a neuropathological perspective, AD has traditionally been associated with the accumulation of beta-amyloid (Aβ) plaques and neurofibrillary tangles composed of hyperphosphorylated tau protein (Selkoe, 2002). However, scientific evidence accumulated over recent decades has shown that these deposits alone do not account for the complexity of the disease, its clinical progression, or the variability in treatment response. Rather, AD involves a broad network of interconnected pathological processes, including synaptic loss, neuroinflammation, oxidative stress, mitochondrial dysfunction, alterations in neuronal metabolism, DNA damage, and aberrant cell-cycle reactivation (López-Sánchez et al, 2021; Orr et al, 2019; Nelson et al, 2013).

AD is increasingly recognized as a disorder of synaptic dysfunction (Selkoe, 2002), with synapse loss correlating more closely with cognitive impairment than neuronal loss and often preceding overt neurodegeneration (Tönnies & Trushina, 2017; Barthet & Mulle, 2020; Meftah & Gan, 2023; Yadav et al., 2026). Multiple homeostatic systems, including calcium regulation, proteostasis, mitochondrial function, endolysosomal integrity, and redox balance, become disrupted early in disease progression and converge on synaptic dysfunction (McDaid et al., 2020; Cozachenco et al., 2023; Sharma et al., 2026; Wal et al., 2026). Calcium dyshomeostasis, driven in part by aberrant ER calcium release and Aβ-induced dysregulation of NMDA and AMPA receptor signaling, impairs synaptic plasticity and can activate pathways associated with excitotoxicity, oxidative stress, neuroinflammation, and mitochondrial dysfunction (Mustaly-Kalimi et al., 2022; Zhang et al., 2022; Chaudhary et al., 2025). In parallel, impaired proteostasis and endolysosomal function disrupt protein degradation and synaptic autophagy, promoting the accumulation of pathological and synaptic proteins and contributing to synaptic dysfunction (Cozachenco et al., 2023; Mustaly-Kalimi et al., 2022; Jasutkar et al., 2024). Mitochondrial dysfunction further compromises ATP availability, calcium handling, redox balance, and mitochondrial trafficking, creating an energy deficit that limits the ability of synapses to sustain neurotransmission and plasticity (Tönnies & Trushina, 2017; Sharma et al., 2026; Wal et al., 2026).

These disturbances are further amplified by chronic neuroinflammation and aberrant complement activation. Aβ-driven activation of microglia and astrocytes can reactivate developmental complement pathways involving C1q, C3, and CR3, promoting excessive pruning of glutamatergic synapses before substantial neuronal loss occurs (Aranda-Abreu et al., 2026; Zhang et al., 2024). Oxidative stress additionally damages synaptic proteins and cellular components, further impairing synaptic plasticity and neurotransmission (Tönnies & Trushina, 2017; Yadav et al., 2026). Collectively, calcium dysregulation, proteostasis failure, mitochondrial and metabolic dysfunction, oxidative stress, and neuroimmune activation are homeostatic alterations mechanistically interconnected that can reinforce one another, producing progressive structural and functional synaptic deterioration (McDaid et al., 2020; Behroozi et al., 2026). This self-reinforcing disruption of neuronal homeostasis provides a mechanistic link between molecular pathology, synaptic failure, hippocampal circuit dysfunction, and cognitive decline, with synaptic dysfunction emerging years before extensive neuronal loss and the full clinical manifestation of AD (Song et al., 2019; Ebrahimi et al., 2025; Sharma et al., 2026).

Synaptic long-term potentiation (LTP), a fundamental form of synaptic plasticity and a key mechanism underlying learning and memory (Bliss and Collingridge, 1993; Malenka, 2003; Wu et al, 2019), is profoundly impaired in AD (Di Lorenzo et al, 2020), contributing to cognitive deficits, and preceding neuronal death (Di Lorenzo et al, 2020; Selkoe, 2002). In humans, transcranial magnetic stimulation studies demonstrate early alterations in LTP-like plasticity (Di Lorenzo et al, 2016; Jannati et al, 2023) that correlate with amyloid pathology (Buss et al, 2020) and predict both cognitive decline (Motta et al, 2018) and conversion from mild cognitive impairment to dementia (Di Lorenzo et al, 2020). In cell culture and animal studies, oligomeric amyloid has been shown to damage synapses and block LTP induction by altering several mechanisms (Knafo et al, 2016), including the function of NMDA and mGluR receptors (Skaper et al, 2017; Benaroch et al, 2018). Here we explore the possibility that LTP can be restored once it has been lost in AD.

The transcription factor E2F4 regulates neuronal homeostasis (Ramón-Landreau et al, 2022), and can be phosphorylated by the stress-activated p38^MAPK^ at a Thr motif orthologous to Thr248/Thr250 in the human sequence (Morillo et al, 2012), disrupting this protective function (Llabrés-Mas et al, 2026), while a phosphorylation-resistant variant of human E2F4 (hE2F4DN) maintains neuronal stability and prevents AD pathology in a mouse model of AD (López-Sánchez et al, 2021). We hypothesized that hE2F4DN might restore mechanisms of synaptic plasticity even after LTP has already been lost in an AD mouse model, by re-establishing physiological E2F4-dependent homeostatic regulation, thereby recovering cognitive function and preventing further neuronal death. To test this hypothesis across different stages of disease progression, we investigated hE2F4DN at an early stage, when its effects could reflect prevention of pathological progression, and at a later stage, when synaptic dysfunction and LTP impairment were already fully established. We further used hippocampal proteomic and phosphoproteomic profiling at the early stage to define the molecular programs underlying hE2F4DN-mediated neuroprotection, and at the late stage to identify synaptic changes potentiated by hE2F4DN. Together, these complementary approaches allowed us to examine both the molecular basis of E2F4-dependent protection during early disease and the capacity of this intervention to restore synaptic function once pathological dysfunction was established.

## MATERIALS AND METHODS

### Mice

Male and female mice aged 1.5–7 months were used in this study. All animal experiments were approved by the bioethics committee of the Consejo Superior de Investigaciones Científicas (Ref. 1235/2022) and the Autonomous Government of Madrid (PROEX 182.4/22), in compliance with the Spanish and European Union guidelines. Animals were housed under standard conditions of temperature (22±1 ◦C) and humidity (40–60%) and on a 12 h light/dark cycle. Control C57BL/6J mice were purchased from ENVIGO. Double transgenic mice in C57BL/6J genetic background expressing under the control of the Thy1 promoter both mutant human amyloid precursor protein 695 (APP695) with the Swedish (K670N, M671L), Florida (I716V), and London (V717I) FAD mutations and human PS1 harboring the M146L and L286V FAD mutations (Tg6799 or 5xFAD mice) (Oakley et al, 2006) were purchased from The Jackson Laboratory (strain #008730). These mice were PCR genotyped as indicated by The Jackson Laboratory. Homozygous 5xFAD (h5xFAD) mice, known to show an aggravated phenotype (Richard et al 2015), were obtained after repeated inbreeding of originally hemizygous mice (López-Sánchez et al. 2021). h5xFAD mice always yielded genetically homogeneous litter when crossed with wild-type (WT) mice. Homozygosity was confirmed by qPCR genotyping using specific primers of the human APP transgene [upstream primer: 5’-AGGATGGTGATGAGGTAGAG-3’ (from NM_201414 sequence) and downstream primer complementary to: 5’ CTGCTGTTGTAGGAACTCGA-3’ (896-915 bp and 1011-1030 bp from the NM_201414 sequence, respectively)], with the *Kcn3a* genomic sequence as a reference^18^.

### Viral vectors

Neuronal expression of human E2F4DNmyc and E2F4DN was achieved using the AAV-AS viral vector (Choudhury et al., 2016) containing the neuron-specific hSyn1 promoter (Kügler et al., 2001), referred to as dsAAV-AS.hSyn1.hE2F4DNmyc.WPRE3SL (AAV.hE2F4DNmyc) (Figure S1C) and dsAAV-AS.hSyn1.hE2F4DN.WPRE3SL (AAV.hE2F4DN) (Figure 1A), respectively. To generate these latter vectors, the human E2F4 coding sequence (or the human E2F4 coding sequence followed by the myc tag) with Thr248Ala/Thr250Ala mutations (hE2F4DN) and containing the Kozak sequence (ACCATGG) and a final stop codon were synthesized (GeneScript) and cloned into the pcDNA 3.1(+) vector (Invitrogen). After confirming their sequences by Sanger sequencing, hE2F4DN or hE2F4DNA-myc were PCR amplified and seamless cloned (Gibson et al., 2009) into the KD10 plasmid (Packgene), flanked by the hSyn1 promoter and the mutant WPRE sequence (WPRE3SL), which lacks tumorigenic capacity (Schambach et al., 2006). The expected sequence was confirmed by Sanger sequencing. The vectors were then manufactured as previously described (Choudhury et al., 2016) by co-transfecting each of the KD10 transgene plasmid together with the adenoviral helper plasmid pAdDeltaF6 (Plasmid #112867, Addgene), and a 5:1 ratio of the pGG-9.47_VP1,3 #2515 (Plasmid #74237; Addgene) and the pGG-9.47_VP2_AS #2562 (Plasmid #74238; Addgene) packaging plasmids. The final purified products were dialyzed in PBS with additional 0.001% Pluronic F-68. Viral titers were determined by qPCR and confirmed by silver stain.

**FIGURE 1.**
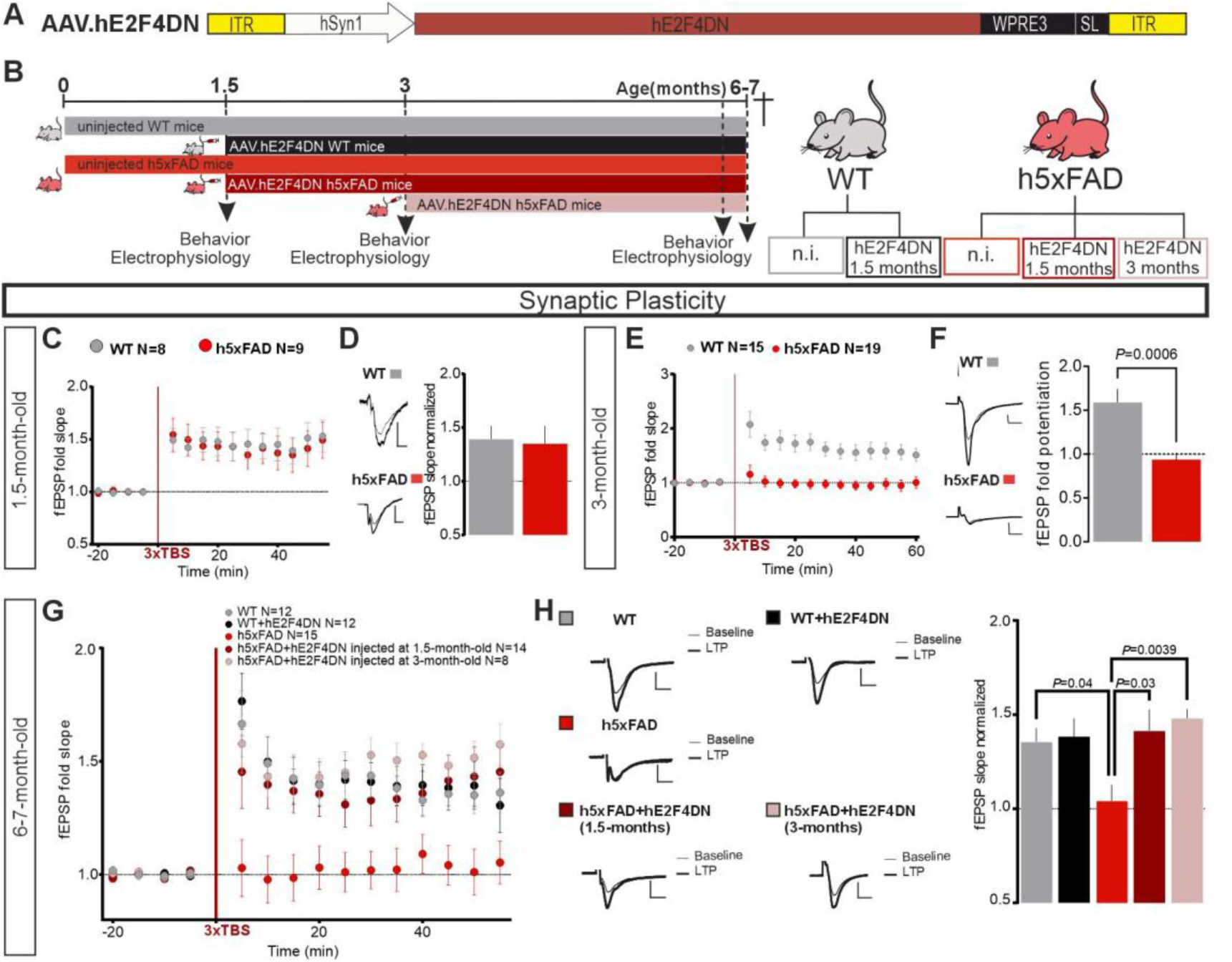
Intravenous administration of AAV.hE2F4DN rescues LTP in h5xFAD mice. **(A)** Scheme showing the genomic sequence of the AAV.hE2F4DN vector. **(B)** Scheme of the design of the experimental groups and the timing for treatments. **(C-H)** LTP recordings, induced in the CA1 from WT or h5xFAD mice, stimulating axons arising from the CA3 in untreated mice of 1.5 months **(C)** and 3 months **(E)**, or 6 months **(G)**, treated as indicated. *N* represents the number of slices from 5 WT and 6 h5xFAD untreated 1.5-month-old mice **(C)**; 13 WT and 13 h5xFAD untreated 3-month-old mice **(E)**; and the following untreated and treated 6-7 months old mice: 9 WT, 6 WT+hE2F4DN, 11 h5xFAD mice, 8 h5FAD+hE2F4DN treated at 1.5 months and 6 h5FAD+hE2F4DN treated at 3 months old mice **(G)**. **(D, F, H)** Left: representative traces of fEPSPs of the basal response (thin line) and after LTP induction (thick line). Right: quantification of mean fEPSPs slopes 45 min after induction at 1.5 months **(D)**, 3 months (*P* value was determined using a Mann–Whitney test) **(F)**, or 6-7 months [*H*(4) = 13.91, *P* = 0.007, Kruskal-Wallis test; *P* values were derived using Dunn’s multiple *post hoc* test] (H). Scale bars represent 0.5 mV, 10 ms.

For the proteomic and phosphoproteomic analyses of h5xFAD mice treated at 1.5 months of age, hE2F4DN was expressed using the previously described AAV.PHP.B-hSyn1.hE2F4DN vector (López-Sánchez et al., 2021), hereafter also referred to as AAV.hE2F4DN.

Neuronal expression of mouse E2F4DN (mE2F4DN) was carried out using the AAV-PHP.eB vector, which also crosses the blood brain barrier (Zhang & Jiao, 2015). The mouse E2F4 (mE2F4) coding sequence (NCBI accession number NM_148952) with Thr249Ala/Thr251Ala mutations (mE2f4dn) or the GFP coding sequence, both containing the Kozak sequence, flanked by the human hSyn1 promoter and the WPRE3SL sequence were synthesized (GenScript) and cloned into the pcDNA 3.1 (+) plasmid (Invitrogen). The expected sequences of the hSyn1-mE2f4dn-WPRE3SL or hSyn-GFP-WPRE3SL cassettes were confirmed by Sanger sequencing. These cassettes, flanked by ITR2 sequences, were subcloned into a transfer plasmid to generate recombinant single stranded AAV vectors referred to as dsAAV.PHP.eB.hSyn1.mE2F4DN.WPRE3SL (AAV.E2F4DN) or dsAAV.PHP.eB.hSyn1.GFP.WPRE3SL (AAV.GFP) [Viral Vector Production Unit (UPV), Barcelona, Spain]. These AAV vectors, employed for learning-induced proteomics analysis, were intravenously administered through the tail vein at 1.25 x 10^10^ vg/g body weight (Lozano-Ureña et al., 2024).

For *in vitro* experiments, the serotype 5 adenoviral vector (Ad5) (Davidson et al., 1993) was used to generate Ad5.hSyn1.hE2F4DNmyc.WPRE3SL (Ad5.hE2F4DNmyc) vector (Viral Vector Production Unit, Autonomous University of Barcelona). This vector contains the hSyn1 promoter followed by the specific transgene and the WPRE3SL post-transcriptional regulatory element described above. Hippocampal primary cultures were treated with these viruses (50 infective units/cell) at 14-17 DIV and used for immunocytochemistry experiments after 3 DIV to allow transgene expression (Figure S2A).

### AAV-AS dose determination

To determine the optimal AAV-AS dose, we evaluated two doses: a weight-adjusted dose based on a previous study in which intravenous administration of 1 × 10¹² vg/mouse of the AAV.PHP.B-hSyn1.hE2F4DN vector ameliorated AD-related phenotypes in h5xFAD mice (López-Sánchez et al., 2021), and a second dose corresponding to half that amount. Therefore, WT female mice (1.5 months old, ∼16 g) received intravenous injections of the AAV.hE2F4DN vector (Figure 1A), for qRT-PCR, or the AAV.hE2F4DNmyc vector (Figure S1C), for imaging experiments, at 3.125 × 10¹⁰ or 6.25 × 10¹⁰ vg/g, corresponding to 5 × 10¹¹ and 1 × 10¹² vg/mouse, respectively (Figure S1A). One month post-injection, qRT-PCR analysis showed a dose-dependent increase in hE2F4DN expression (Figure S1B), with ∼60% of cortical neurons positive for the myc tag at both doses (Figure S1C, D). Therefore, the lower dose was selected for all subsequent experiments, with the AAV.hE2F4DN vector administered intravenously via the tail vein at 3.125 × 10¹⁰ vg/g body weight. Transgene expression remained stable up to 7 months, as confirmed by qPCR, in h5xFAD mice administered at 1.5 months of age (Figure S1E, F).

### Antibodies

Immunocytochemistry: The rabbit anti-Myc-tag antibody (ab9106, Abcam) was used at 1/250 dilution, the mouse anti-NeuN (clone A60 MAB377, Merck) was used at 1/800 dilution, the chicken anti-MAP2 antibody (ab5392, Abcam) was used at 1/1500 dilution, the rabbit anti-Synapsin I antibody (ab64581, Abcam) was used at 1/100 dilution and the mouse anti-E2F4 (clone LLF4-2) antibody (MABE160, Merk) was used at 1/200. The following secondary antibodies were used at 1/1,000 dilution: Donkey anti-Rabbit IgG (H+L) Highly Cross-Adsorbed Secondary Antibody Alexa Fluor 488 (A21206, Life Technologies), Goat anti-Mouse IgG (H+L) Highly Cross-Adsorbed Secondary Antibody Alexa Fluor 488 (A-11029, Life Technologies), Goat anti-Chicken IgY (H+L) Secondary Antibody Alexa Fluor 647 (A-21449, Life Technologies), Goat anti-Rabbit IgG (H+L) Highly Cross-Adsorbed Secondary Antibody Alexa Fluor 594 (A-11037, Life Technologies), Goat anti-mouse IgG (H+L) Highly Cross-Adsorbed Secondary Antibody Alexa Fluor 568 (A-11004, Invitrogen).Western Blot: The rabbit anti-β-Amyloid pAb #2454 (Cell Signaling Technology, Danvers, MA) was diluted 1/1000, the mouse anti-Actin antibody, clone C4 (MAB1501R, Merk) was diluted 1/20,000, the, anti-E2F4 (E3G2G) Rabbit mAb (40291, Cell Signaling Technology) was diluted 1/6000, the rabbit anti-phospho-tau (Ser396) polyclonal antibody (44-752G, Invitrogen) at 1/2000 and the mouse monoclonal anti-tau antibody at 1/2000 (MABG3420, Merk). The following secondary antibodies were used at 1/15,000 dilution: IRDye® 680RD Goat anti-Mouse IgG Secondary Antibody (926-68070) (LI-COR) and IRDye® 800CW Goat anti-Rabbit IgG Secondary Antibody (926-32211) (LI-COR).

### Mouse hippocampal cultures

Primary hippocampal neurons were prepared as previously described (Walton et al., 2019) from CD1 mouse embryos at E17. Imaging studies were performed at 17-20 DIV. Briefly, pups were decapitated and brains were placed in cold HBSS, without calcium or magnesium (Gibco #14185-052) with Hepes (1:100, Gibco). Meninges were then removed, and hippocampi were carefully dissected. Hippocampi were incubated for 15 min at 37 °C in HBSS containing 2.5 mg/ml trypsin (Gibco) in a 15 ml conical centrifuge tube, 1 mg/ml DNAse (Roche) was added for the last 3 minutes. After chemical dissociation the reaction was stopped, and hippocampi were washed 3 times with 5 ml of HBSS at 37 °C each time. Then, hippocampi were mechanically dissociated by passing them through the enlarged bore of a Pasteur pipette 10–15 times. Non-dissociated tissue was allowed to be collected at the bottom by gravity and clean supernatant with dissociated cells transferred to a new 15 ml centrifuge tube. 1 ml of HBSS at 37 °C was added to the remaining tissue and dissociated again 10–15 times with a flame polished Pasteur pipette to reduce the diameter of the tip. Non-dissociated tissue was allowed to be collected at the bottom by gravity. Clean supernatant with dissociated cells was transferred to the same centrifuge tube in which cells from the first dissociation were collected. Then, HBSS containing hippocampi were centrifuged at 300 x g for 5 min and suspended in plating medium, DMEM (Gibco) containing 10% inactivated Horse Serum (Gibco) and penicillin-streptomycin (25 U/ml) (Gibco). Neurons were plated at a density of 15,000 cells (imaging studies) or 30,000 cells (electrophysiology) on 10-mm diameter coverslips (Menzel Glässer) previously coated with poly-L-lysine (Sigma-Aldrich), prepared at 1 mg/ml in 150 mM borate buffer, pH 8.4. Neurons were allowed to attach to coverslips for 1 to 2 h in a humidified atmosphere containing 5 % CO2 at 37 °C. Once neurons were settled, neuronal plating medium was washed with maintenance medium consistent of Neurobasal without L-Glutamine (Gibco) medium supplemented with B27 (Gibco), penicillin-streptomycin (25 U/ml) (Gibco) and GlutaMAX (Gibco). Half of the culture medium was exchanged by fresh maintenance medium every 3–4 days.

### Thioflavin S Staining

Thioflavin staining was performed as previously described [10]. Vibratome sections were washed 3 times with PBS containing 0.4% Triton X-100 (Sigma-Aldrich) and then incubated for 30 min in the dark with 0.05% thioflavin S (Sigma-Aldrich) in 50% ethanol (Merck). Finally, sections were washed twice with 50% ethanol and once with MilliQ water and mounted in InmunoSelect Antifading Mounting Medium. Samples were analyzed in duplicate; 2 slices per animal were used to obtain a mean value for each animal. Images were acquired with the Fluorescence Thunder microscope and Fiji ImageJ software was used for the analysis.

### Immunocytochemistry and image acquisition

Primary mouse hippocampal cultures were fixed for 15 min with 4% PFA at room temperature (RT) and then incubated for 30 min at RT with 0.1% Triton X-100 phosphate buffered saline (PBS-T) containing 10% fetal bovine serum (FBS) (Gibco), to block antibody unspecific binding. Coverslips containing neurons were subsequently incubated with the primary antibody diluted to the appropriate concentration in PBS-T containing 1% FBS (PBS-T/1% FBS) for 1 h in the case of mouse neurons, and overnight for human neurons. After 3 washes in PBS-T/1% FBS, incubation was carried out for 1 h at RT in PBS-T/1% FBS containing 100 ng/ml DAPI (Sigma-Aldrich) and secondary antibodies. Finally, 3 washes were performed with PBS-T and one with PBS. Coverslips were mounted in ImmunoSelect® Antifading Mounting Medium (Quimigen). Confocal images were acquired with a Leica SP5 confocal microscope.

### Immunohistochemistry and image acquisition

Mice were transcardially perfused with PBS, and then with 4% PFA, and brains were postfixed O/N at 4 °C with 4% PFA. Vibratome obtained 50-µm sections were permeabilized and maintained for 1 h at RT in Blocking Buffer (BB) containing phosphate buffered saline (PBS)/0.2% Triton X-100 (Sigma-Aldrich) (0.1% PBTx) and 10% FBS, and incubated O/N at 4 °C with the primary antibodies in BB. After washing with BB, the sections were incubated for 1 h at RT in BB containing 100 ng/ml DAPI with the secondary antibodies. The sections were washed in three times with 0.1% Triton X-100 PBS and twice with PBS before mounting with ImmunoSelect Antifading Mounting Medium. Images were obtained with a SP5 confocal microscope. Counting of Myc-positive -positive nuclei in brain sections was performed using z-projections as described previously (López-Sánchez et al., 2021).

### qPCR

qPCR genotyping used to confirm h5xFAD homozygosity was performed as previously described^18^. Reverse transcriptase (RT)-qPCR (qRT-PCR) was performed with the 7500 real-time PCR equipment (Applied Biosystems), using the SsoAdvanced Universal SYBR Green Supermix (BioRad) and specific primers for the house keeping gene Rps18 (qMmuCED0045430) and the human E2F4 gene (qHsaCED0048496). Standard curve method was used to calculate the quantity of both genes and transgene expression levels were relativized to those of Rps18.

### Ex-vivo Field Recordings

Acute slice preparation was performed as previously described (Knafo et al., 2016). To prepare acute slices, mice were cervical dislocated and decapitated. The brain was rapidly removed to ice-cold, oxygenated, dissection solution (10 mM D-Glucose, 4 mM KCl, 26 mM NaHCO3, 233.7 mM sucrose, 5 mM MgCl2, 1:1,000 Phenol Red). Coronal hippocampal slices (300 μm) were made with a vibratome (LeicaVT1200S), and then they were transferred to a recovery chamber containing ACSF (119 mM NaCl, 2.5 mM KCl, 1 mM NaH2PO4, 11 mM glucose, 26 mM NaHCO3, 1.2 mM MgCl2, 2.5 mM CaCl2) at 24–26 °C for at least 1.5 h before recording.

For field recording, fEPSPs were recorded from the stratum radiatum of CA1 with glass microelectrodes (700 kΩ-1 MΩ resistance) filled with ACSF, used also as external solution. To assess the function of excitatory synapses, electrophysiological recordings were performed in the presence of picrotoxin (100 μM), to block GABAA receptor responses. fEPSPs were evoked by stimuli (100 µs) delivered through a concentric bipolar electrode (FHC), located in CA1 stratum radiatum, to record the fEPSPs while stimulating presynaptic Schaffer collateral fibers. Field EPSPs slope was used to assess synaptic strength. For each slice, an input–output response curve was recorded to compare basal synaptic transmission (BST) at different conditions (Sánchez-Puelles et al., 2020). The response curve was obtained by varying the intensity of single-pulse stimulation from 10 to 250 µA and averaging 5 responses per intensity. This curve was also used to set the baseline fEPSP at ≈30% of maximal slope. Following the baseline period (10-20 minutes of stable responses), LTP was induced through a three theta-burst stimulation (TBS) protocol (each burst consisting of 4 pulses at 100 Hz repeated with 200 ms inter-burst intervals) (Kimura et al., 2009), and the responses were recorded for 1 h. All electrophysiological data were collected with pCLAMP software (Molecular Devices) and analyzed blindly.

### Hippocampal extracts

Extracts from hippocampus were obtained in cold extraction buffer (20 mM Tris–HCl pH 6.8, 10 mM β-mercaptoethanol, 1 mM EDTA, 1% Triton X-100, 1% SDS) including Complete Mini, EDTA-free, protease inhibitor cocktail (Roche) (one hippocampus in 500 μl extraction buffer). Extracts were centrifuged for 10 min at 14,000 × g (at 4 °C), and supernatants were then boiled for 5 min in either tricine loading buffer (50 mM Tris-HCl pH 8, 12% Glicerol, 4% SDS, 0.001% Coomasie blue, 2% β-mercaptoethanol), for Aβ analysis, or Laemli buffer (62.5 mM Tris-HCl pH 6.8, 2% SDS, 10% Glicerol, 2% β-mercaptoethanol, 0.005% Bromphenol blue), for E2F4 and tau quantification.

### Western blot

For the analysis of Aβ levels, hippocampal extracts were fractionated on a 16.5% Tris-tricine acrylamide gels (Bio-Rad). For E2F4 and tau analyses hippocampal extracts were fractionated on 4-20% TGX-precast protein gels (Bio-Rad). Then, they were transferred to 0.2 µm Low-Fluorescence PVDF transfer membranes (Thermo Scientific), for Aβ analysis, or to 0.45 μm Immobilon-FL transfer membranes (Millipore), for other analyses. The membranes were incubated for 1 h at RT with OBB (LI-COR) and then incubated ON at 4 °C with the primary antibodies at the appropriate dilution in OBB containing 0.1% Tween 20 (Sigma-Aldrich) (OBB-T). After washing the membranes with OBB-T, they were incubated for 1 h at RT with the secondary antibodies in OBB-T. Finally, the membranes were washed again with OBB-T, and the protein bands were visualized using the Odyssey CLx Infrared Imaging System (Li-COR). Finally, the Image J software (Fiji) was used for the quantification of the bands.

### Unbiased stereology

Mice were anesthetized with 1µl/g Pentobarbital (Vetoquinol) injected intraperitoneally and perfused with PBS followed by 4% PFA prepared in PBS. The brains were post-fixed O/N in 4% PFA and cryoprotected in 3% glycerol (Panreac) and 3% ethylene glycol (Panreac) prepared in 0.04 M Phosphate Buffer, pH 7.4. Serial coronal sections (50 μm) obtained with a Leica VT1000 S Vibrating blade microtome (Leica) were Nissl stained with 0.1% cresyl violet (Sigma-Aldrich) and then mounted on histological gelatinized glass slides (Sigma-Aldrich). For each brain, CA1 hippocampal neurons from sections at −1.6 to 3.3-3.8 mm from bregma were quantified stereologically according to the Allen Mouse Brain Atlas (https://mouse.brain-map.org) as a reference, following as previously described (Sánchez-Puelles et al., 2020). The Stereo Investigator software (Version 5.65, MBF Bioscience) was used both to drive a motorized stage (mac 6000 system, MBF Bioscience) on a dual optical head microscope (Nikon Eclipse 80i) and to mark and measure CA1 hippocampal region at 2x (Nikon Microscope Objective Plan UW 2x/0.06 7.5mm WD) under brightfield optics. Any incomplete stained or torn section was considered a random event, designated missing, and then accounted for in the stereological estimate using the fractionator principle (Stereo Investigator software v5.65, MBF Biosciences). Thereafter, the software sequentially chose counting frames (100×100 μm) randomly in the xyz axes, moving the motorized stage automatically within the previously delimited zones in the CA1 region at 100x (Nikon Microscope Objective Plan Apo VC 100x/1.4 Oil ∞/0.17 DIC N2). Neuronal nuclei were then randomly sampled from the defined CA1 region using optical dissector probes, and a neuronal population was determined using a 2 μm top and bottom guard zone CA1 (counting frame, 15×15; sampling grid, 100x100; dissector height, 30 μm; interval, 6). The average number of sampling sites per animal was 290.5± 14.65. Cells were marked only if their edges were within the dissector area and they did not intersect forbidden lines. The cells were also marked if they came into focus as the optical plane moved through the height of the dissector. The precision of our stereological counts was determined by the Gunderson coefficient of error equation^26^. This method of sampling and the section intervals were tested in a pilot experiment to ensure that the number of cells estimated was representative of their total number.

### Behavioral Assays

Animals were handled for 5 min in the experimental room on each of the 3 days before beginning the experiments. Experimental mice were transferred to the behavioral testing room 30 min prior to beginning the first trial to habituate them to the room. To avoid the influence of circadian fluctuations, the behavioral tests were always performed in the morning. Light levels were controlled in all the behavioral experiments. All apparatuses were cleaned with 0.1% Acetic Acid between mice to remove odors.

#### Novel Object Location Task (NOL)

This assay evaluates hippocampal-dependent memory and was selected for its low-stress nature, minimizing stress-related effects on cognitive performance (Knafo et al., 2016). Mice were first individually habituated for 30 min to an empty open field white box [40×30×15(height) cm] during three consecutive days prior to the test. The box presented different patterns in each wall (triangles, lines, circles and stars) as spatial cues. The NOL task consisted of 2 phases: sample phase, and test phase. The sample phase (object exposure) consisted of placing the mouse into the box that contained two identical objects during 10 min. Animals were free to explore the box and objects during this time. Once the sample phase was finished the mouse was removed from the test box and, after a 60 min delay (retention period), the mouse was placed back into the test box to perform the memory evaluation test. The test (choice session) consisted of switching the location of one of the objects (Novel Object Location trial) and letting the animal freely explore for 5 minutes. Spatial memory performance was assessed by comparing exploration time between the displaced object (novel location, NL) and the non-displaced object (familiar location, FL), with preferential exploration of the NL object indicating memory retention. The relative exploration of the novel object was expressed by a discrimination index [DI = (t NL -t FL)/(t NL + t FL)]. In addition, total exploration of both objects was quantified as a control measure of overall exploratory activity. No significant differences were observed between groups at 1.5 months (Figure S3A), three months (Figure S3B), or five–six months of age (Figure S3C). Behavior was recorded the day of the experiment and analyzed later using EthoVision XT 11.5 software or Any-Maze, with manual scoring conducted blind to experimental conditions. Exploration occurs when the animal is faced to the object and the nose is closed to it. The investigator recorded the times the mouse was actively investigating each object, which consists of its nose pointed at an object at a maximum distance of 2 cm from that object.

#### Contextual Fear Conditioning Task (CFC)

This assay is a form of associative learning depending on hippocampus in which a neutral stimulus (a given spatial context, i.e. contextual stimulus) is associated with an aversive event (an electric shock, i.e. unconditioned stimulus). This test consists in two day sessions: conditioning and memory recall sessions (Stefanelli et al., 2016). During the conditioning session, each mouse was individually placed in a methacrylate cage (inner dimensions 25cm x 25cm x 35cm (height) containing a metallic grid floor and walls covered with different patterns (black and white lines, squares and smooth gray) to produce contextual cues, which was inserted into a sound-attenuating chamber. Mice were allowed to explore for 3 min and then, three mild electric shocks (0.5 mA, 2 s) were delivered through the stainless-steel floor grid with 30 s interval. Mice stayed 1 min additionally before being returned to the home cage (total conditioning time: 5 min). The memory recall session was performed 24 hours later. Mice were placed back in the cage described above (conditioning context) for 5 minutes, during which no footshock was delivered. Mice exhibited a range of conditioned fear responses, including freezing, defined as behavioral immobility except for movement needed for respiration (Fanselow et al., 2003). Freezing was automatically scored and monitored by video recordings, which were digitized using Any-Maze Video Tracking System version 6.05 (Stoelting Co). Raw data, including freezing total time, was exported from Any-Maze. Freezing total time was divided by total time session to obtain the percentage of time freezing, which indicates the level of memory recall of mice. The percentage of freezing during the first 3 minutes of the training session, prior to shock delivery, was evaluated to ensure that no animals exhibited generalized fear behavior and to confirm that there were no differences between groups in baseline fear levels before exposure to the unconditioned stimulus (Figure S4D). To evaluate if there were any differences in pain sensitivity between groups, that could affect the results of this experiment, pain threshold evaluation was done after the CFC test. In the conditioning chamber, after 120 s of resting time, the electric current was gradually increased at a rate of 0.05 mA every 20 s until the animal showed the first signs of pain (jumping and vocalization). At this moment, the current was immediately switched off and the animal was back to its home cage after 30s of resting. The corresponding current was scored as a measure of the animal’s pain threshold. Thus, we confirmed that there were no significant differences between groups in pain sensitivity (Figure S4E). Both assays, CFC and pain sensitivity threshold, were recorded using a camera fixed above the apparatus and connected to a computer.

### Omics data generation

Hippocampal proteins were extracted with RIPA (Sigma) supplemented with 1:100 diluted protease and phosphatase inhibitors (MedChemExpress), and reduced, alkylated, and digested with trypsin and LysC (Thermo Fisher Scientific) in 50 mM ammonium bicarbonate buffer (ABC) according to established protocols (Montero-Calle et al., 2025). For bulk proteomics analysis, 20% of the resulting digested peptides were dried under vacuum on a Speed-Vac, resuspended in 20 µL of 0.1% formic acid (FA), and 1 µL were injected into the mass spectrometer. For phosphoproteomics analysis, 80% of the resulting digested peptides were dried under vacuum and enriched in phosphopeptides using ZrO_2_ and TiO_2_ magnetic beads (Resyn Bioscience). Briefly, peptides were resuspended in 200 µl of 80 % acetonitrile (ACN), 5 % trifluoroacetic acid (TFA), 0.1 M glycolic acid and incubated with 5 µl of previously equilibrated magnetic beads for 20 min at RT and 1000 rpm. Supernatants were recovered and labeled as flow-through (FT). Beads were then washed with 500 µl of 80 % ACN, 1 % TFA (2 min, RT, 1000 rpm) and 500 µl of 10 % ACN, 0.2 % TFA (2 min, RT, 1000 rpm). Then, phosphopeptides were eluted by a 10 min incubation at RT and 1000 rpm with 200 µl of 1 % NH4OH. A second and third rounds of enrichment using the obtained FTs were performed as described but with 5 µl (second round) or 10 µl (third round) of equilibrated beads. The eluates were vacuum-dried, resuspended in 10 µl of 0.1 % FA, and 8.5 µl injected into the mass spectrometer.

Bulk proteomics and phosphoproteomics analysis were performed using label free quantification (LFQ) data independent acquisition (DIA) in the Orbitrap Astral mass spectrometer (Thermo Fisher Scientific) coupled to a Vanquish Neo or Evosep One HPLC system.

For the analysis on the Orbitrap Astral coupled to the Vanquish Neo UHPLC System (Thermo Fisher Scientific), peptides were loaded into the precolumn PepMap Trap Catridge 5 µm, 300 µm x 5 mm (Thermo Fisher Scientific) and eluted in an Easy-Spray PepMap RSLC C18 2 µm, 75 µm x 15 cm (Thermo Fisher Scientific) heated at 50°C, as previously described (Montero-Calle et al., 2026). For the analysis on the Orbitrap Astral coupled to the Evosep One (Evosep), peptides were loaded onto Evotip Pure tips (Evosep) and separated using the standardized 60-sample-per-day (60SPD) method. Peptides were passed through an 8 cm column (150 μm inner diameter) packed with 1.7-μm C18 beads (IonOpticks), heated to 50 °C, into the precolumn PepMap Trap Catridge 5 µm, 300 µm x 5 mm (Thermo Fisher Scientific).

For mass spectrometry analysis, ionization was set at 1900 V of liquid junction voltage and 280°C capillary temperature was used. The full scan method employed a m/z 380-980 mass selection, an Orbitrap resolution of 240000 (at m/z 200), an automatic gain control (AGC) value of 500%, and maximum injection time (IT) 5 ms. The MS/MS was performed with the Astral mass analyzer, using an AGC of 500%, an IT of 3 ms, and a normalized collision energy (NCE) of 25 for fragmentation of precursors. The scan range was set from 380 to 980 m/z, with an isolation window of 2 m/z, and window placement optimization was enabled. Thus, a total of 299 windows were analyzed in each cycle.

For the analysis of the global proteome, raw data were analyzed with Spectronaut (version 21.0.260609.94842) using standardized workflows. Spectral libraries were generated using the directDIA workflow against the Uniprot UP000000589.fasta Mus musculus (December, 2025) reference proteome (17,234 protein entries), supplemented with the human E2F4 protein sequence obtained from UniProt. Trypsin/P and Lys/P were selected as the digestion enzymes and a maximum of 2 missed cleavages were allowed. Carbamidomethylation of cysteines was set as a fixed modification, and methionine oxidation and N-terminal acetylation were set as variable modifications. A standard workflow was used for DIA analysis. The maximum FDR for peptide spectral match (PSM), peptide, and protein identifications was set at 0.01, automatic cross-run normalization was enabled, and all identified peptides were used for protein quantification. Protein inference was performed using the IDPicker algorithm. Non-imputation was performed during peptide and protein identification and quantification with Spectronaut software.

Raw data phosphoenriched samples were analyzed as above, using standard PTM workflows. Briefly, STY phosphorylation was included as variable modification for Pulsar search, cross-run normalization was performed based on STY phosphopeptides, and a minimum PTM localization probability of 0.75 was required.

### Omics preprocessing and quality control

Two independent hippocampal cohorts were analyzed. In the early-treatment cohort, h5xFAD mice received AAV.hE2F4DN at 1.5 months of age and tissue was collected at 6 months under basal conditions. Proteomics included WT (*n* = 5), untreated h5xFAD (*n* = 6) and hE2F4DN-treated h5xFAD (*n* = 6) mice after facility quality control; phosphoproteomics included *n* = 6 animals per group. In the late-treatment learning cohort, h5xFAD mice received control AAV.GFP or AAV.E2F4DN at 3 months of age. At six months of age, mice were either maintained under basal conditions or subjected to CFC conditioning session 1 h before sacrifice and tissue collection. The experimental groups were GFP-basal (*n* = 4), GFP-CFC (*n* = 6), E2F4DN-basal (*n* = 5) and E2F4DN-CFC (*n* = 6).

Facility-provided abundances were analyzed without additional normalization. Proteins were retained if quantified in at least five replicates in any early-treatment group or at least 75% of samples in any late-treatment group. Phosphoproteomic features were retained if quantified in at least 50% of samples in any group. Missing values were not imputed for differential or enrichment analyses; for phosphoproteomic PCA only, missing values were temporarily replaced by feature medians before analyzing the 1,000 most variable features.

Protein abundance data were analyzed using facility-provided protein accession identifiers. Corresponding mouse gene symbols were used for functional annotation and selected figure labels, whereas protein symbols or commonly accepted names were used elsewhere. Phosphoproteomic features were labeled by protein symbol, phosphorylation site, and, where applicable, phosphorylation multiplicity. Gene-symbol labels in proteomic figures represent quantified proteins, not transcript abundance.

Primary condition-only models and sensitivity models additionally adjusted for sex yielded highly concordant effect estimates; therefore, condition-only models were used for primary analyses. PCA and sex-adjusted sensitivity results are presented for early-treatment proteomics (Figures S5–S6), early-treatment phosphoproteomics (Figures S8– S9), late-treatment proteomics (Figures S11–S13), and late-treatment phosphoproteomics (Figures S16–S18).

### Differential omics analyses

Protein abundance and phosphorylation were analyzed using limma with empirical Bayes moderation (Ritchie et al., 2015). Early-treatment contrasts compared untreated h5xFAD versus WT, hE2F4DN-treated versus untreated h5xFAD, and hE2F4DN-treated h5xFAD versus WT. Late-treatment contrasts assessed CFC responses within each treatment, treatment effects under basal and CFC conditions, and the treatment-by-learning interaction. Benjamini–Hochberg-adjusted P < 0.05 defined FDR significance; nominal P < 0.05 was used for predefined exploratory analyses. No minimum absolute log₂ fold-change threshold was applied.

### Functional enrichment analyses

Functional enrichment analyses were performed according to the experimental design and biological questions addressed by each cohort. Broad Gene Ontology Biological Process (GO-BP) annotations were used to characterize molecular pathways associated with cellular homeostasis and neuronal regulation in the early-treatment cohort, whereas synapse-specific SynGO v1.3 Biological Process (BP) and Cellular Component (CC) annotations were used to investigate learning-associated synaptic responses in the late-treatment cohort (Koopmans et al., 2019).

For early-treatment proteomics, over-representation analysis (ORA) was performed using clusterProfiler (Wu et al., 2021) on proteins with nominal P < 0.05, separated by the direction of abundance change, with the corresponding tested protein set as background. Preranked gene set enrichment analysis (GSEA) was additionally performed using fgsea (Korotkevich et al., 2016), with genes ranked according to limma moderated t-statistics. GSEA was performed independently for untreated h5xFAD versus WT and hE2F4DN-treated versus untreated h5xFAD mice. GO-BP pathways meeting nominal P < 0.05 and FDR < 0.25 in both comparisons were evaluated according to their normalized enrichment scores (NES) to identify concordant or opposite-direction enrichment patterns. Opposite-direction pathways were further examined using gene set variation analysis (GSVA). GSVA scores were centered to the WT mean, retaining pathways with an absolute untreated h5xFAD-to-WT mean difference ≥ 0.15 GSVA units. Treatment-associated movement toward WT was defined as a smaller absolute mean deviation from WT in treated than untreated h5xFAD mice. Representative pathways were selected for visualization across major functional categories.

For early-treatment phosphoproteomics, GO-BP ORA was performed on unique genes corresponding to nominally responsive phosphorylation features (P < 0.05), separated by direction, using genes represented by statistically tested phosphoproteomic features as background.

For late-treatment proteomics, learning-associated synaptic responses were evaluated using preranked GSEA with SynGO v1.3 BP and CC annotations. Proteins were ranked according to limma moderated t-statistics for the CFC-versus-basal comparison within each treatment group. Mouse genes were mapped to human orthologs for SynGO annotation, and multiple protein assignments to the same gene were resolved by retaining the representative feature with the largest absolute moderated t-statistic. Terms meeting the exploratory criterion of nominal P < 0.05 and FDR < 0.25 in at least one learning contrast were considered for comparative visualization, with FDR < 0.05 indicated separately. Enriched terms were reduced according to their hierarchical parent– child relationships before leading-edge recurrence analysis to identify proteins contributing to multiple synaptic annotations.

For late-treatment phosphoproteomics, SynGO BP and CC ORA was performed on unique genes corresponding to nominally responsive phosphorylation features (P < 0.05), separated by direction. Mouse genes were mapped to human orthologs, and genes represented by statistically tested phosphoproteomic features in the corresponding contrast were used as background. Enriched terms reaching FDR < 0.05 were considered for the comparative analysis of phosphorylation features decreased following CFC in E2F4DN-treated mice and those exhibiting negative treatment-by-learning interactions.

### Kinase–substrate enrichment and network analysis

Early-treatment kinase–substrate enrichment analysis (KSEA) used mouse kinase– substrate annotations from PhosR (Kim et al., 2021a,b). Features mapping to the same gene–phosphosite pair were collapsed by retaining the largest absolute treatment-associated log₂ fold change. Kinases with at least three quantified substrates were tested, with Benjamini–Hochberg correction. KSEA was interpreted as coordinated substrate phosphorylation rather than direct kinase activity. GSK3B-associated phosphosite changes were additionally compared with corresponding total-protein abundance changes.

For the late-treatment cohort, proteins associated with negative treatment-by-learning interaction phosphofeatures and SynGO categories enriched at FDR < 0.05 were analyzed using STRING v12.5 (Szklarczyk et al., 2025). Protein associations were evaluated using human identifiers, a minimum combined score of 0.40, and no additional interactors. Representative phosphosites were selected using a predefined ranking integrating network degree, the number of negative-interaction phosphofeatures per protein, and site-level interaction evidence.

### Statistical analyses

#### Non-Omics statistics

Quantitative data are represented as the mean ± standard error of the mean. Statistical tests were carried out using GraphPad Prism (GraphPad Software; www.graphpad.com). Before performing any statistics, normality was evaluated. The data that passed normality

Shapiro-Wilk test was analyzed using parametric tests. When two groups were analyzed, statistical differences were obtained between using two-tailed Student’s t test, meanwhile when 3 or more groups are analyzed, one-way ANOVA analysis was performed followed by Tukey’s or Dunnett’s post hoc test. Two-way ANOVA analysis followed by Bonferroni post hoc test was also performed for grouped analysis. The data that did not pass normality test was analyzed through nonparametric test, being the Mann-Whitney test used to compare two groups or Krustal-Wallis with Dunńs multiple comparisons post-test for 3 or more groups. Outliers, as evidenced by the Grubbs’ test (α=0.05), also called the ESD method (extreme studentized deviate), were eliminated from the analysis. The exact number of animals used per experiment (N) are indicated in the figures. The details of the statistical test employed for each experiment appear in the figure legends.

#### Omics statistical analyses

Analyses were performed in R as described above. Benjamini–Hochberg FDR < 0.05 was considered statistically significant unless otherwise indicated. Nominal P < 0.05 was restricted to predefined exploratory analyses.

## RESULTS

### hE2F4DN prevents LTP loss in homozygous 5xFAD (h5xFAD) mice

Given that E2F4 lacks a conventional nuclear localization signal (Puri et al., 1998) and contains functional nuclear export signals (Gaubatz et al., 2001), we sought to determine whether hE2F4DN localizes to specific cytoplasmic domains in neurons. To this end, mouse primary hippocampal neurons were transduced with an adenoviral vector (Ad5.hE2F4DNmyc) encoding Myc-tagged hE2F4DN (hE2F4DNmyc) (Figure S2A). Immunofluorescence analysis revealed that hE2F4DNmyc was present in the neuronal cytoplasm (Figure S2B, D–G), including axons (Figure S2D, E), dendrites, and dendritic spines (Figure S2B, C). Notably, hE2F4DNmyc was also detected in close proximity to synapsin-positive structures (Figure S2F, G), suggesting that *hE2F4DN may be associated with synaptic compartments and potentially contribute to synaptic function*.

To assess the effects of hE2F4DN on synaptic function, we intravenously administered an adeno-associated viral vector (AAV.hE2F4DN) encoding a dominant-negative form of human E2F4 carrying the Thr248Ala/Thr250Ala mutations to 1.5-month-old h5xFAD mice (Figure 1A, B), a well-characterized model of Alzheimer’s disease (AD) (Richard et al., 2015; López-Sánchez et al., 2021) in which hippocampal long-term potentiation (LTP) is markedly impaired by 6 months of age in heterozygous animals (Kimura et al., 2008).

At 1.5 months of age, h5xFAD mice exhibited normal LTP, comparable to WT mice (Figure 1C, D), as assessed by ex vivo recordings of field excitatory postsynaptic potentials (fEPSPs) at hippocampal CA3–CA1 synapses, a circuit critical for memory formation (Bartsch et al., 2011). Mice treated with AAV.hE2F4DN at 1.5 months of age were analyzed at 6–7 months, when h5xFAD mice exhibit advanced AD-like pathology (Richard et al., 2015). Notably, while h5xFAD mice had lost the ability to induce LTP, AAV.hE2F4DN-treated h5xFAD mice retained synaptic potentiation at levels comparable to those observed in WT mice (Figure 1G, H).

As expected, BST was comparable between 1.5-month-old h5xFAD and WT mice (Figure S3A). Consistent with the beneficial effect of hE2F4DN on LTP, BST was significantly enhanced in hE2F4DN-treated h5xFAD mice at 6–7 months of age (Figure S3C). In contrast, AAV.hE2F4DN administration had no detectable effect on BST in WT mice (Figure S3C), suggesting that the beneficial effects of hE2F4DN are preferentially manifested under pathological conditions.

Taken together, *these data indicate that neuronal expression of hE2F4DN at an early, non-pathological stage preserves LTP in h5xFAD mice during advanced stages of pathology*.

### *In vivo* administration of AAV.hE2F4DN reduces Aβ pathology in h5xFAD mice

Given the involvement of Aβ in synaptic dysfunction and neuronal death in both AD patients and mouse models (Buss et al., 2020; Knafo et al., 2016), we investigated the impact of hE2F4DN on Aβ production and deposition in h5xFAD mice. We first examined mice treated at 1.5 months of age, extending our previous findings (López-Sánchez et al., 2021). To this end, we assessed soluble Aβ levels (Figure 2A–E) and amyloid plaque burden (Figure 2F–H) in WT and h5xFAD mice, including control and hE2F4DN-treated animals.

**FIGURE 2.**
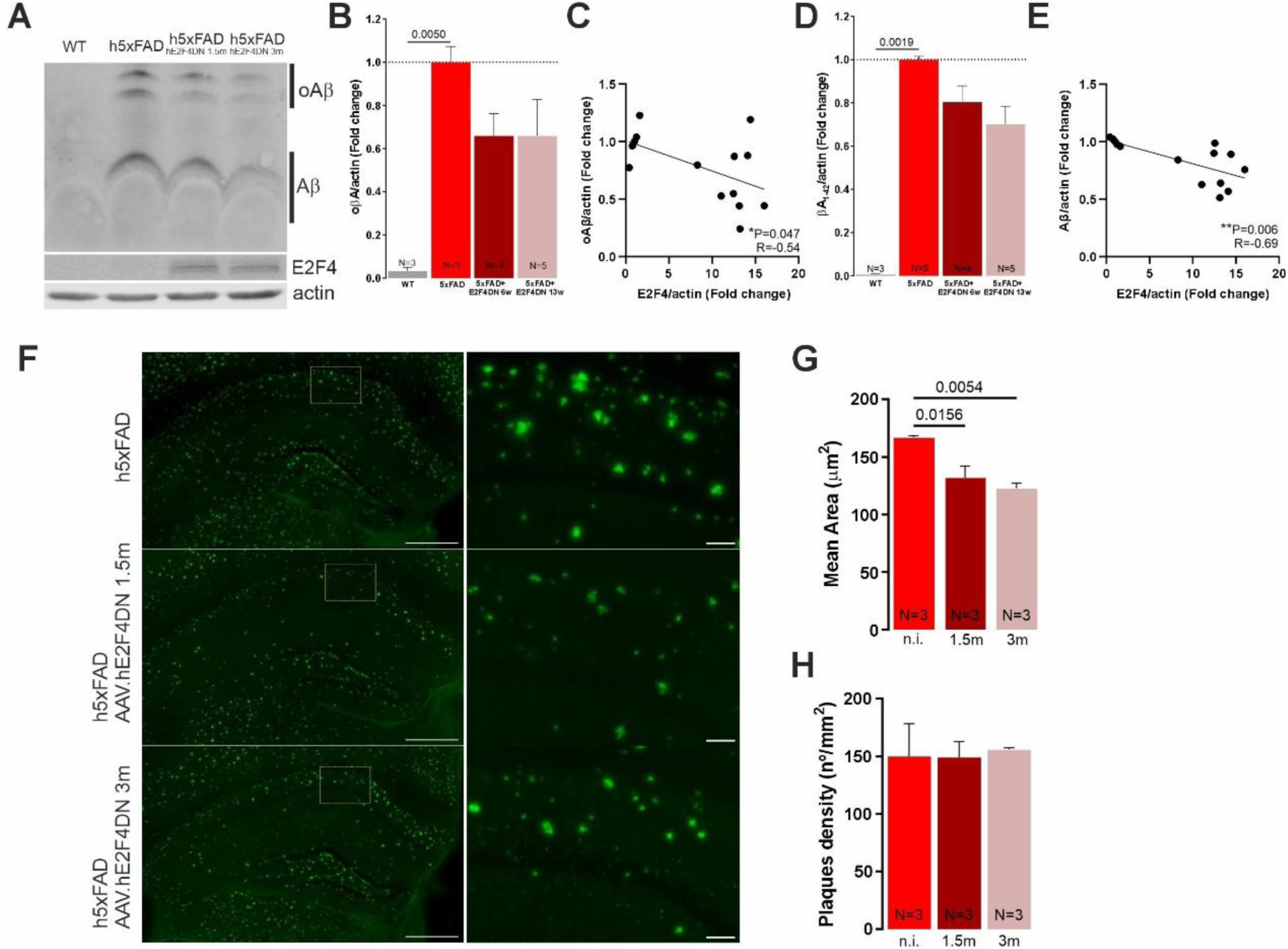
Intravenous administration of AAV.hE2F4DN reduces Aβ pathology in h5xFAD mice. Quantification of soluble Aβ levels **(A-E)** and Aβ plaques **(F-H)** in hippocampus of WT and h5xFAD mice, treated or untreated with AAV.hE2F4DN at 1.5 o 3 months. **(A)** Representative immunoblot of oligomeric (oAβ) or monomeric (Aβ) Aβ species, E2F4 and actin of the indicated experimental groups. **(B)** Bar graph representing oAβ levels vs. actin, normalized to non-treated h5xFAD. **(C)** Correlation between oAβ and E2F4 levels. **(D)** Bar graph representing Aβ levels vs. actin, normalized to non-treated h5xFAD. **(E)** Correlation between Aβ and E2F4 levels. **(F)** Left: representative images of Aβ plaques in hippocampal slices. No plaques were observed in WT. Right: high magnification of the yellow squares in left panels showing a section of the CA1 region at higher magnification. **(G)** Bar graph showing the area occupied by the plaques in the different groups. **(H)** Bar graph showing plaque density in the different groups. **p<0.01, *p<0.05 in **C**, **E** (Pearson’s *R* correlation test). *P* values in **B**, **D**, **G** were obtained by One-way ANOVA test [F (2, 6) = 12.82; *P*=0.0068] followed by Dunnett’s multiple comparisons test. Scale bars: 500 μm (**F** left), 50 μm (**F** right).

We found a significant increase of soluble Aβ monomers and oligomers (oAβ) when hippocampal fractions from 6-month-old h5xFAD mice were compared to those from WT mice, an effect that was reduced by AAV.hE2F4DN (Figure 2A, B, D). This finding was further substantiated by a significant negative correlation between the measured amount of E2F4 and that of both oAβ (Figure 2C) and Aβ (Figure 2E). Furthermore, the mean area of Aβ plaques in the hippocampus of 6-month-old h5xFAD mice treated with AAV.hE2F4DN at 1.5 months of age was significantly reduced compared with control-treated h5xFAD mice (Figure 2F, G). In contrast, plaque density remained unchanged (Figure 2H). Taken together, *these data suggest a reduction in Aβ-pathology due to hE2F4DN-neuronal expression at early stages of the pathology*.

### hE2F4DN blocks neuronal death in h5xFAD mice

Previous analyses in heterozygous 5xFAD mice demonstrated significant neuron loss at 9 months of age (Oakley et al., 2006), an effect likely contributing to the progression of the cognitive decline observed in these mice and paralleling the course of AD in humans (Arendt et al., 2015).

To assess the consequences of hE2F4DN expression on neuronal survival we used unbiased stereology to determine the number of neurons in the hippocampal CA1 pyramidal layer of WT and h5xFAD mice. Stereological quantification revealed a 44% reduction in CA1 pyramidal neuron number (Figure 3A-C) and a 38% decrease in CA1 volume (Figure 3A,B,D) in untreated 6-month-old h5xFAD mice relative to WT controls. AAV.hE2F4DN administration at 1.5 months fully preserved both neuronal number and regional volume (Figure 3A-D). WT mice receiving AAV.hE2F4DN showed values comparable to untreated WT animals (Figure 3A-D). *These findings demonstrate that hE2F4DN expression prevents CA1 neurodegeneration in h5xFAD mice*.

**FIGURE 3.**
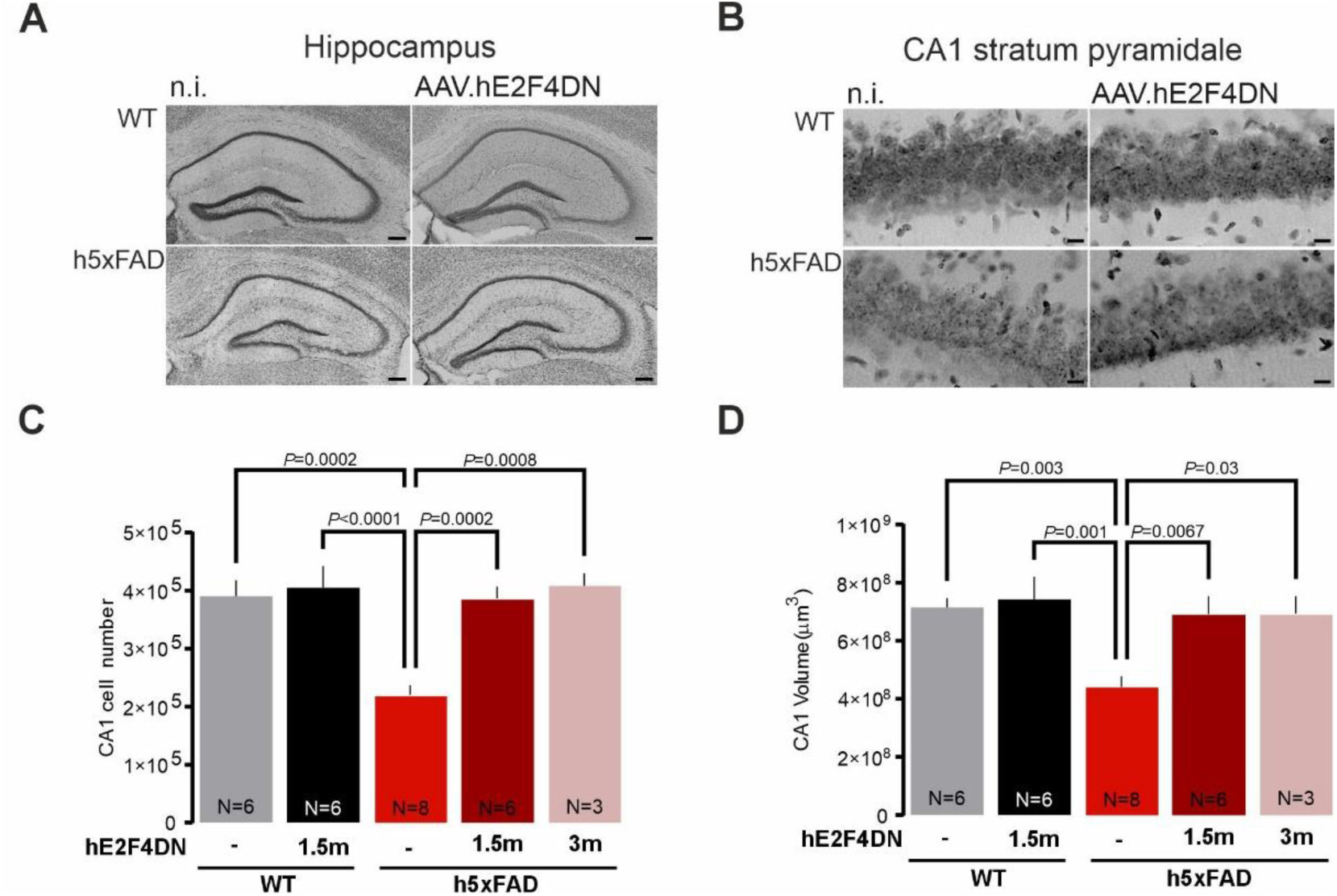
Intravenous administration of AAV.hE2F4DN improves neuronal survival in h5xFAD mice. **(A)** Nissl-stained hippocampal sections from WT (top) and h5xFAD (bottom) mice either non-injected (left) or injected (right) with AAV.hE2F4DN at 1.5 months. **(B)** Representative images of Nissl-stained CA1 stratum pyramidale from a WT mouse (top) and h5xFAD mice (bottom) either non-injected (left) or injected (right) with AAV.hE2F4DN at 1.5 months. **(C)** Number of CA1 neurons in 6-month-old mice of the indicated experimental groups [*F*(4, 24) = 11.3, *P* <0.0001, One-way ANOVA]. *P* values shown in the graphs were derived using Dunnett’s multiple comparisons test **(D)** CA1 volumes in 6-month-old mice of the indicated experimental groups [*F*(4, 24) = 6.227, *P*=0.001, One-way ANOVA]. *P* values shown in the graphs were derived using Dunnett’s multiple comparisons test. 1.5m: AAV.hE2F4DN administration at 1.5 months, 3m: AAV.hE2F4DN administration at 3 months. Scale bars: 200 µm **(A)**, 10 µm **(B)**. *N* represents number of animals.

### Proteomics profiling highlights metabolic remodeling alongside neuroimmune and amyloid-associated responses after hE2F4DN treatment

To investigate the long-term molecular effects of early hE2F4DN administration, we analyzed the hippocampal proteomes of 6-month-old WT mice, untreated h5xFAD mice, and h5xFAD mice treated with hE2F4DN at 1.5 months of age. Six mice were analyzed per condition. However, one 6-month-old WT mice, identified as an outlier, was excluded from the final analysis because it exhibited a markedly different global protein intensity distribution compared with the other biological replicates. Proteomic profiling revealed a treatment-associated response (Figure 4A, B), with increased abundance of proteins involved in mitochondrial and redox homeostasis, including methylenetetrahydrofolate dehydrogenase 1-like (*Mthfd1l*), glutathione reductase (*Gsr*), biliverdin reductase A (*Blvra*) and Lon peptidase 1 (*Lonp1*), as well as neuronal signaling and structural proteins such as DARPP-32 (*Ppp1r1b*), MARCKS-like protein 1 (*Marcksl1*), and UNC79 (*Unc79*). Conversely, regulator of G-protein signaling 14 (*Rgs14*), involved in hippocampal synaptic plasticity, and CRMP4 (*Dpysl3*), associated with cytoskeletal organization, were decreased.

**FIGURE 4.**
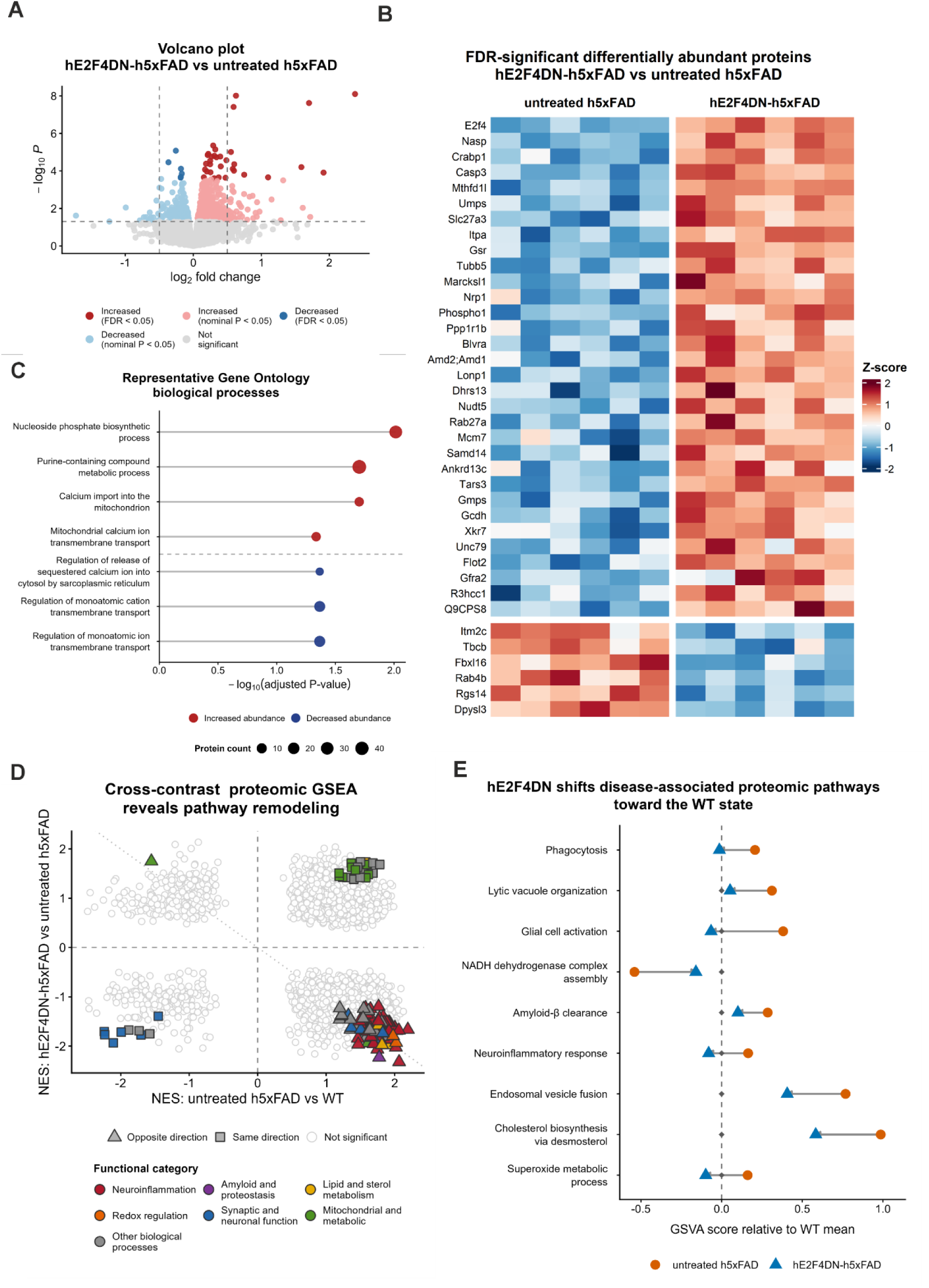
Proteomic profiling highlights metabolic, neuroimmune and amyloid-associated responses to hE2F4DN. Proteomic analysis included WT (*n* = 5), untreated h5xFAD (*n* = 6) and hE2F4DN-treated h5xFAD mice (*n* = 6). **(A)** Volcano plot of differential protein abundance in hE2F4DN-treated versus untreated h5xFAD hippocampi. Of 690 proteins with nominal *P* < 0.05, 38 reached FDR < 0.05, including 32 increased and six decreased proteins (Table S2). Dark shading indicates FDR-significant proteins, light shading nominal-only proteins (*P* < 0.05, FDR ≥ 0.05), and grey non-significant proteins. Vertical lines at |log₂ fold change| = 0.5 are shown for reference only and were not used as significance thresholds. **(B)** Heatmap of the 38 FDR-significant proteins showing abundance as row-wise Z-scores across individual untreated and hE2F4DN-treated h5xFAD samples. Proteins are ordered by decreasing limma moderated *t*-statistic, and corresponding gene symbols are displayed. For visualization only, five missing values across four proteins were replaced by the mean abundance of the corresponding protein; all statistical analyses used the original non-imputed data. **(C)** Representative GO-BP terms significantly enriched among nominally responsive proteins, analyzed separately according to the direction of abundance change in the hE2F4DN-treated versus untreated h5xFAD comparison. Increased proteins were enriched in nucleotide/purine metabolism and mitochondrial Ca²⁺ transport, whereas decreased proteins were associated with Ca²⁺- and ion-homeostasis processes. The x-axis represents −log₁₀(FDR), point size indicates protein count, and color denotes direction of change. Complete enrichment results are provided in Table S3. **(D)** Comparative proteomic Gene Set Enrichment Analysis (GSEA) of disease- and treatment-associated GO-BP pathways. Normalized enrichment scores (NES) for hE2F4DN-treated versus untreated h5xFAD mice are plotted against those for untreated h5xFAD versus WT mice. A total of 3,506 pathways with available NES values in both contrasts are displayed. Pathways meeting the predefined exploratory enrichment criterion (nominal *P* < 0.05 and FDR < 0.25 in both contrasts) are highlighted according to functional category, whereas pathways not meeting this joint criterion are represented by pale open circles. Among the 121 highlighted pathways, 90 exhibited opposite NES directions between disease- and treatment-associated comparisons, whereas 31 retained the same direction. Triangles distinguish pathways with opposite-direction enrichment, while squares indicate pathways with same-direction enrichment. Point color denotes functional category. Complete GSEA results are provided in Table S4. **(E)** GSVA-based assessment of representative pathways exhibiting opposite-direction GSEA enrichment and treatment-associated shifts toward the WT state. Of the 90 pathways showing opposite-direction enrichment in GSEA, 86 met the minimum disease-associated GSVA deviation criterion (|mean GSVA score in untreated h5xFAD − mean GSVA score in WT| ≥ 0.15). Among these, 57 showed mean scores closer to WT following treatment. Nine representative pathways spanning neuroimmune/inflammatory, amyloid/lipid/redox, and cellular/mitochondrial processes are shown. Individual-sample profiles are provided in Figure S7.

GO-BP enrichment analysis of differentially abundant proteins in hE2F4DN-treated versus untreated h5xFAD mice revealed an enrichment of upregulated proteins in processes related to nucleotide and purine metabolism and mitochondrial Ca²⁺ transport. Conversely, downregulated proteins were enriched in processes associated with ion transport and intracellular Ca²⁺ release (Figure 4C). These findings suggest that hE2F4DN treatment may modulate Ca²⁺ signalling pathways, potentially contributing to the restoration of Ca²⁺ homeostasis and neuronal excitability in this AD mouse model.

To investigate broader disease-associated molecular programs, we compared GSEA profiles from untreated h5xFAD versus WT and hE2F4DN-treated versus untreated h5xFAD mice. Most pathways meeting the predefined exploratory criteria in both contrasts showed opposite normalized enrichment score (NES) directions, including neuroimmune and inflammatory regulation, amyloid-β fibril formation and clearance, lipid/sterol metabolism, redox homeostasis, intracellular trafficking and mitochondrial function (Figure 4D). Consistently, sample-level GSVA showed movement toward WT values for pathways associated with phagocytosis, glial activation, amyloid-β clearance, cholesterol and superoxide metabolism, intracellular trafficking and NADH dehydrogenase complex assembly (Figure 4E; Figure S7).

*Together, these findings identify prominent metabolic and redox remodeling after hE2F4DN treatment, accompanied by broader neuroimmune, amyloid-associated and mitochondrial responses that may explain the reduced deposition of Aβ and the prevention LTP impairment and neuronal death triggered by hE2F4DN*.

### Phosphoproteomics profiling highlights synaptic-vesicle regulation and GSK3B-associated phosphosignaling

Having characterized treatment-associated changes in the hippocampal proteome, we next investigated whether hE2F4DN also modified protein phosphorylation, a regulatory mechanism involved in neuronal signalling and synaptic function.

Phosphoproteomic profiling revealed a focused treatment-associated response, with 461 phosphorylation features showing nominal changes and four remaining significant after multiple-testing correction (Figure 5A, B). AP-3 complex subunit β2 (*Ap3b2*)-S733 showed increased phosphorylation following hE2F4DN treatment, whereas Transformer-2 protein homolog α (*Tra2a*)-S259 and Serine/arginine repetitive matrix protein 1 (*Srrm1*)-S655/S657 were decreased. Notably, AP-3 complex subunit β2 is a neuronal component of the AP-3 complex involved in synaptic-vesicle biogenesis and trafficking. Consistent with this observation, GO-BP enrichment analysis identified synaptic vesicle cycle and synaptic vesicle localization among the significantly affected biological processes (Figure 5C).

**FIGURE 5.**
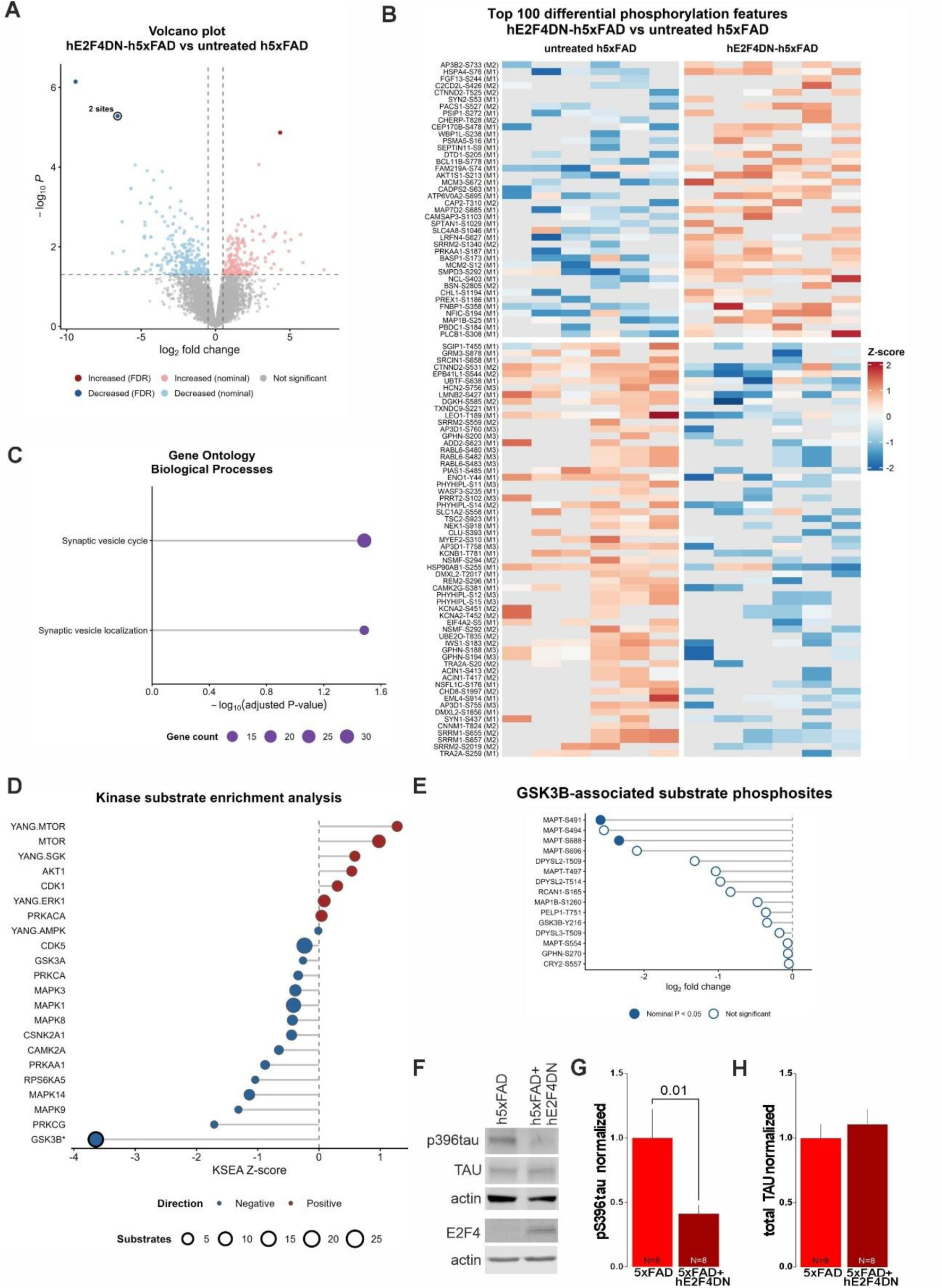
Phosphoproteomic profiling highlights synaptic-vesicle regulation and GSK3B-associated phosphosignaling. Phosphoproteomic analysis included WT, untreated h5xFAD and hE2F4DN-treated h5xFAD mice (*n* = 6 per group). **(A)** Volcano plot of differential phosphorylation in hE2F4DN-treated versus untreated h5xFAD hippocampi. A total of 461 phosphorylation features showed nominal *P* < 0.05, including 170 increased and 291 decreased features, mapping to 307 unique genes. Four features reached FDR < 0.05: AP3B2-S733 increased, whereas TRA2A-S259, SRRM1-S655 and SRRM1-S657 decreased (Table S5). Dark shading indicates FDR-significant features, light shading nominal-only features (*P* < 0.05, FDR ≥ 0.05), and grey indicates non-significant features. Vertical lines at |log₂ fold change| = 0.5 are shown for reference only and were not used as significance thresholds. SRRM1-S655 and SRRM1-S657 overlap because they have identical statistical estimates. **(B)** Heatmap of the 100 nominally responsive phosphorylation features with the largest absolute limma moderated *t*-statistics, displayed as row-wise Z-scores across individual untreated and hE2F4DN-treated h5xFAD samples (*n* = 6 per group). Features are grouped by direction of phosphorylation change and ordered by decreasing signed moderated *t*-statistic. Individual phosphosite identities are displayed as gene symbol, phosphorylation site and phosphorylation multiplicity [GENE-SITE (M1/M2/M3)]. Samples are grouped by treatment condition without individual sample labels. Missing values are shown in light grey and were not imputed. **(C)** Gene Ontology Biological Process (GO-BP) over-representation analysis of genes associated with nominally responsive phosphorylation features, using genes represented by statistically tested phosphoproteomic features as background. Only the two processes reaching FDR < 0.05 are displayed: synaptic vesicle cycle and synaptic vesicle localization. The x-axis represents −log₁₀(adjusted *P*-value), and point size indicates the number of genes contributing to each enriched term. Complete GO-BP enrichment results are provided in Table S6. **(D)** Primary kinase–substrate enrichment analysis (KSEA) using one representative phosphorylation feature per unique gene–phosphosite pair, selected according to the largest absolute log₂ fold change while retaining its direction. All 22 evaluated kinase-associated signatures are displayed and individually labeled. KSEA Z-scores indicate the direction of substrate-level phosphorylation changes, with point colour representing positive or negative enrichment and point size indicating the number of quantified substrates. GSK3B was the only kinase-associated signature reaching FDR < 0.05, showing negative enrichment (Z = −3.64, FDR = 0.0059; 15 substrates). The significant GSK3B signature is highlighted by a black outline and an asterisk (Table S7). **(E)** Treatment-associated log₂ fold changes of the 15 GSK3B-associated phosphosites contributing to the primary KSEA signature. All 15 exhibited negative treatment-associated changes. Filled circles indicate individually nominally significant phosphosites (*P* < 0.05), whereas open circles indicate non-significant phosphosites. None reached FDR < 0.05 individually. Comparison with the corresponding total-protein measurements is provided in Figure S10. (**F**) showing phospho-tau (pTau Ser396), total tau, and E2F4 expression in the indicated experimental groups, with their respective actin loading controls. (**G**) Bar graph representing pTau Ser396 levels/tau vs. actin, normalized to non-treated h5xFAD (*P* value was determined using a Mann–Whitney test). (**H**) Bar graph representing total tau vs. actin, normalized to non-treated h5xFAD.

Kinase–substrate enrichment analysis identified GSK-3β (*Gsk3b*) as the only kinase-associated signature remaining significant after multiple-testing correction (Figure 5D). The phosphosites contributing to this signature showed a coordinated negative treatment-associated response and included several MAPT sites together with neuronal substrates such as CRMP2 (*Dpysl2*), Microtubule-associated protein 1B (*Map1b*), Gephyrin (*Gphn*) and Regulator of calcineurin 1 (*Rcan1*) (Figure 5E). Comparison with the corresponding total-protein measurements showed that GSK3β-associated phosphorylation changes were largely not accompanied by comparable changes in protein abundance, particularly for tau (*Mapt*), supporting a predominantly phosphoregulatory response (Figure S10). Tau is a well-established substrate of GSK3β, which phosphorylates tau at multiple residues, including Ser396 (Li & Paudel, 2006), a phosphorylation site associated with pathological tau found in paired helical filaments (PHFs), the major constituents of neurofibrillary tangles in AD (Goedert et al., 1994). Increased tau phosphorylation at Ser396 has also been reported in 5xFAD mice (Kanno et al., 2014). Since phosphorylation at this residue was not detected in our phosphoproteomic analysis, due to the limited phosphopeptide coverage associated with trypsin-based digestion, we assessed tau phosphorylation at Ser396 by Western blot in hippocampal samples from 6-month-old h5xFAD mice that were either untreated or treated with AAV.hE2F4DN at 1.5 months of age (Figure 5F). Consistent with the reduced GSK3β activity inferred from kinase–substrate enrichment analysis, we observed a significant decrease in tau phosphorylation at Ser396 following hE2F4DN treatment (Figure 5F, G), while total tau levels remained comparable to those in untreated mice (Figure 5F, H).

*Collectively, these observations indicate synaptic-vesicle regulation and coordinated GSK3B-associated substrate phosphorylation as the predominant phosphoregulatory responses to hE2F4DN treatment*.

### hE2F4DN reverses LTP loss, and prevents Aβ pathology, and neuronal death in h5xFAD mice treated at 3 months

To further explore the therapeutic potential of hE2F4DN in AD, we investigated whether its effects extend beyond preventing synaptic dysfunction to restoring synaptic plasticity. By 3 months of age, LTP was abolished in h5xFAD mice (Figure 1E, F), accompanied by impaired BST performance (Figure S3B). Notably, intravenous administration of AAV.hE2F4DN at this stage restored LTP when assessed at 6–7 months to WT levels (Figure 1G, H). This was accompanied by a significant rescue of BSL (Figure S3C). In parallel, neuronal expression of hE2F4DN at this stage reduced Aβ pathology (Figure 2) and resulted in absence of neuronal loss in h5xFAD mice (Figure 3).

*Collectively, these results indicate that neuronal expression of hE2F4DN not only prevents the development of synaptic and neuropathological deficits but can also reduce neuropathological alterations and restore synaptic plasticity after LTP impairment has been established*.

### Late hE2F4DN treatment engages a coordinated synaptic proteomic response to learning

Building on the restoration of hippocampal LTP and the synaptic-related molecular changes identified following E2F4DN treatment, we next investigated whether these effects were accompanied by alterations in the hippocampal proteomic response to learning.

To address this question, we used contextual fear conditioning (CFC), a hippocampus-dependent learning paradigm that engages synaptic potentiation mechanisms involved in memory formation (Knafo et al., 2016, Penn et al., 2017; Subramaniyan et al., 2021).

A separate cohort of h5xFAD mice received either control AAV.GFP or AAV.E2F4DN at 3 months of age. Three months later, mice were either maintained under basal conditions or subjected to a contextual fear conditioning (CFC) session 1 h before sacrifice. Dorsal hippocampal tissue was subsequently analyzed by quantitative proteomics to determine whether E2F4DN modifies the molecular response to learning.

Proteomic profiling following CFC revealed relatively modest changes in individual protein abundance, with no proteins remaining significant after multiple-testing correction in either learning contrast or the formal treatment-by-learning interaction (Figure 6A, B; Figure S14). Nevertheless, SynGO-based preranked GSEA identified broader positive enrichment of pre- and postsynaptic signaling, synapse organization, synaptic-vesicle function and synaptic specialization pathways following CFC in E2F4DN-treated mice, compared with more limited enrichment in GFP controls (Figure 6C).

**FIGURE 6.**
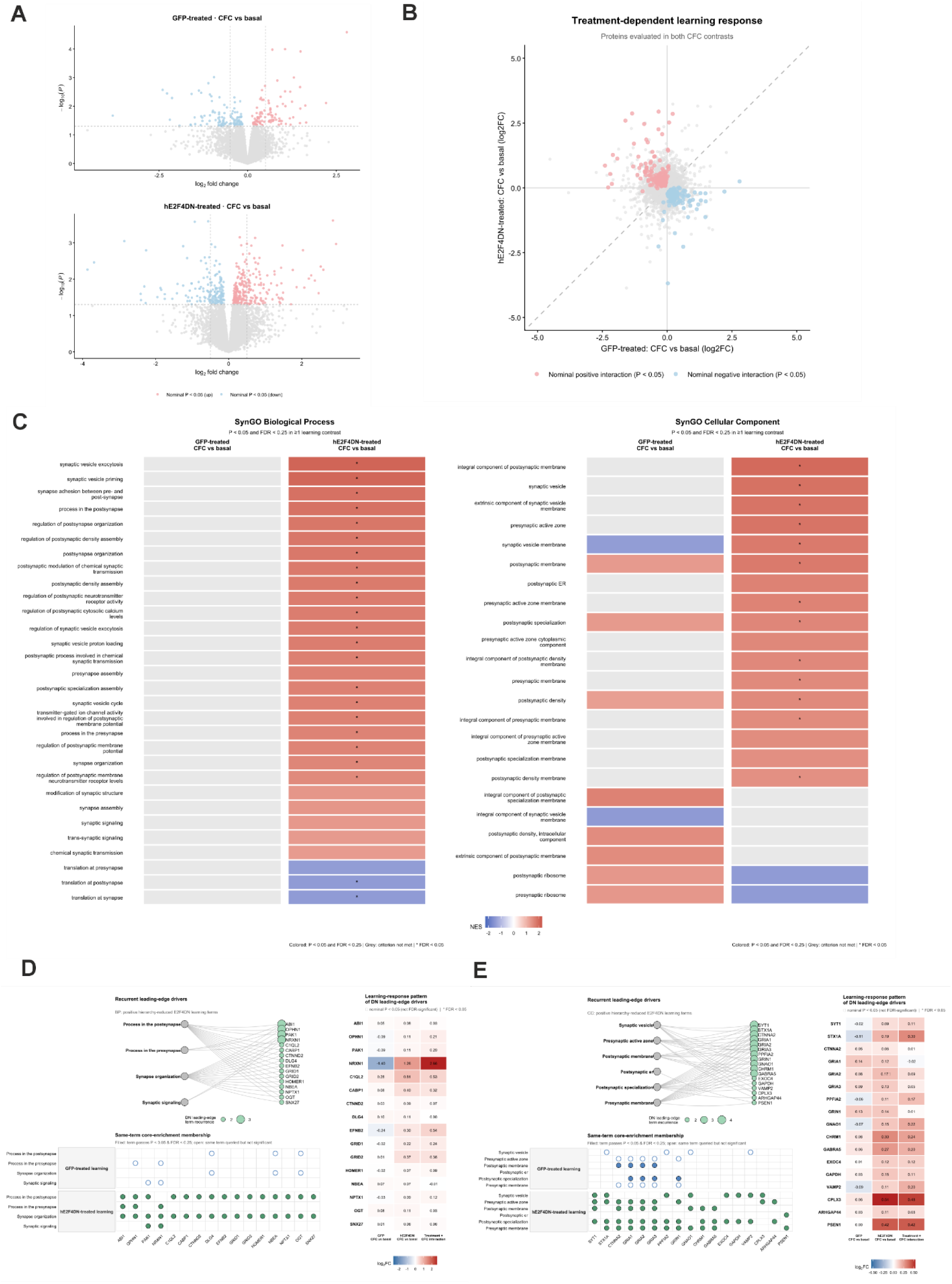
hE2F4DN reshapes the hippocampal proteomic response to learning. Proteomic analysis included GFP-treated (*n* = 10; basal *n* = 4, CFC *n* = 6) and hE2F4DN-treated h5xFAD mice (*n* = 11; basal *n* = 5, CFC *n* = 6). Hippocampal protein abundance was analyzed under basal conditions and following contextual fear conditioning (CFC). **(A)** Volcano plots of differential protein abundance following CFC in GFP- and hE2F4DN-treated h5xFAD mice. Among 8,194 proteins tested in each learning contrast, 218 showed nominal changes in GFP-treated mice (106 increased and 112 decreased), compared with 442 in hE2F4DN-treated mice (262 increased and 180 decreased; *P* < 0.05). No proteins reached FDR < 0.05 in either comparison. Light red and blue points indicate nominally increased and decreased proteins, respectively, whereas grey points indicate non-significant proteins. Vertical lines at |log₂ fold change| = 0.5 are shown for reference only and were not used as significance thresholds. Complete differential abundance results are provided in Table S8. **(B)** Comparison of learning-associated protein abundance changes between treatments. Log₂ fold changes following CFC in hE2F4DN-treated mice are plotted against those observed in GFP-treated mice for 8,182 proteins evaluated in both learning contrasts and the formal interaction analysis. Proteins showing nominal treatment-by-learning interactions (*P* < 0.05) are highlighted in light red (positive interaction) or light blue (negative interaction), whereas non-significant proteins are shown in grey. A total of 296 proteins showed nominal interaction evidence, comprising 161 positive and 135 negative interactions; none reached FDR < 0.05. Learning-associated effect sizes showed limited correlation between treatments (Pearson *r* = −0.050). The dashed diagonal indicates equivalent responses in both groups. The corresponding interaction volcano plot is provided in Figure S14. **(C)** Comparative SynGO preranked Gene Set Enrichment Analysis (GSEA) of learning-associated proteomic responses. Proteins were ranked according to limma moderated *t*-statistics and analyzed using SynGO Biological Process (BP) and Cellular Component (CC) annotations. Complete SynGO enrichment results are provided in Table S9. Terms meeting the predefined exploratory enrichment criterion (nominal *P* < 0.05 and FDR < 0.25 in at least one learning contrast) are shown. GFP-treated mice exhibited no enriched BP terms and 10 CC terms meeting this criterion, whereas hE2F4DN-treated mice showed enrichment of 30 BP and 19 CC terms, including 23 BP and 13 CC terms reaching FDR < 0.05. Positive enrichment in hE2F4DN-treated mice encompassed synaptic-vesicle function, pre- and postsynaptic signaling, synapse organization, presynaptic active zones and postsynaptic specializations. Cell colors represent normalized enrichment scores (NES; red, positive; blue, negative), grey indicates comparisons not meeting the exploratory criterion, and asterisks identify terms with FDR < 0.05. **(D)** Hierarchy-reduced leading-edge analysis of four positively enriched SynGO Biological Process terms following CFC in hE2F4DN-treated mice. Recurrent proteins included glutamatergic receptor components (GRIA1–3, GRIN1) and proteins involved in synaptic adhesion and structural organization (NRXN1, CTNNA2, OPHN1 and PAK1). The network illustrates term–protein associations, the membership matrix compares leading-edge contributions between treatments, and the heatmap shows protein-level log₂ fold changes across both learning contrasts and their formal interaction. **(E)** Corresponding analysis of six positively enriched SynGO Cellular Component terms, highlighting synaptic vesicles, presynaptic active zones and postsynaptic specializations. Recurrent proteins included synaptic vesicle and release machinery (SYT1, STX1A, VAMP2 and CPLX3), together with synaptic scaffolding and receptor-associated proteins (PPFIA2, GRIA1–3 and GRIN1). Graphical conventions follow panel D. In both heatmaps, dots indicate nominal *P* < 0.05 without surviving multiple-testing correction, whereas asterisks indicate FDR < 0.05. The complete SynGO hierarchy is provided in Figure S15.

Hierarchy-reduced leading-edge analysis further identified recurrent proteins contributing across multiple enriched synaptic pathways (Figure 6D, E; Figure S15). These included glutamatergic receptor components involved in excitatory neurotransmission [GluA1–3 (*Gria1-3*) and GluN1 (*Grin1*)], presynaptic proteins regulating vesicle docking, fusion and neurotransmitter release [Syntaxin-1A (*Stx1a*) and Synaptotagmin-1 (*Syt1*)], and proteins involved in synaptic adhesion, cytoskeletal organization and structural remodeling [Neurexin-1 (*Nrxn1*), α-Catenin 2 (*Ctnna2*), Liprin-α2 (*Ppfia2*), Oligophrenin-1 (*Ophn1*), and p21-activated kinase 1 (*Pak1*)].

The recurrence of these proteins across enriched synaptic annotations supported the convergence of the learning-associated proteomic response on both pre- and postsynaptic molecular machinery.

*Collectively, these findings indicate that E2F4DN treatment is associated with a coordinated activity-dependent learning remodeling of synaptic molecular machinery that was not observed to the same extent in control h5xFAD mice*.

### E2F4DN reshapes learning-associated synaptic phosphoregulation

Given the central role of phosphorylation-dependent signalling in synaptic plasticity, we next examined the hippocampal phosphoproteomic response to CFC. Two phosphosites remained significant after multiple-testing correction in the E2F4DN learning contrast: FBXO41 (*Fbxo41*)-S476, associated with a protein involved in synaptic transmission and vesicle release, and VPS13A (*Vps13a*)-S1929, belonging to a lipid-transfer protein. Notably, FBXO41-S476 was also the only phosphosite exhibiting an FDR-significant treatment-by-learning interaction (Figure 7A,B). Neither phosphosite showed corresponding changes of comparable magnitude in total-protein abundance (Figure S19). Beyond these individual sites, treatment-by-learning interactions were predominantly negative (Figure 7B; Figure S20). To characterize this pattern, we compared CFC-associated log₂ fold changes between GFP and E2F4DN mice for the 752 phosphofeatures nominally responsive to CFC (P < 0.05) in at least one group. Changes tended to be positive in GFP controls but negative in E2F4DN-treated mice (Figure 7C). A similar, though less pronounced, directional difference was observed across all 7,734 commonly tested phosphofeatures, independently of statistical significance (Figure S21). These findings suggest that E2F4DN treatment alters the directionality of learning-associated phosphorylation responses rather than simply increasing overall phosphosite abundance, consistent with previous studies (Heo et al., 2023).

**FIGURE 7.**
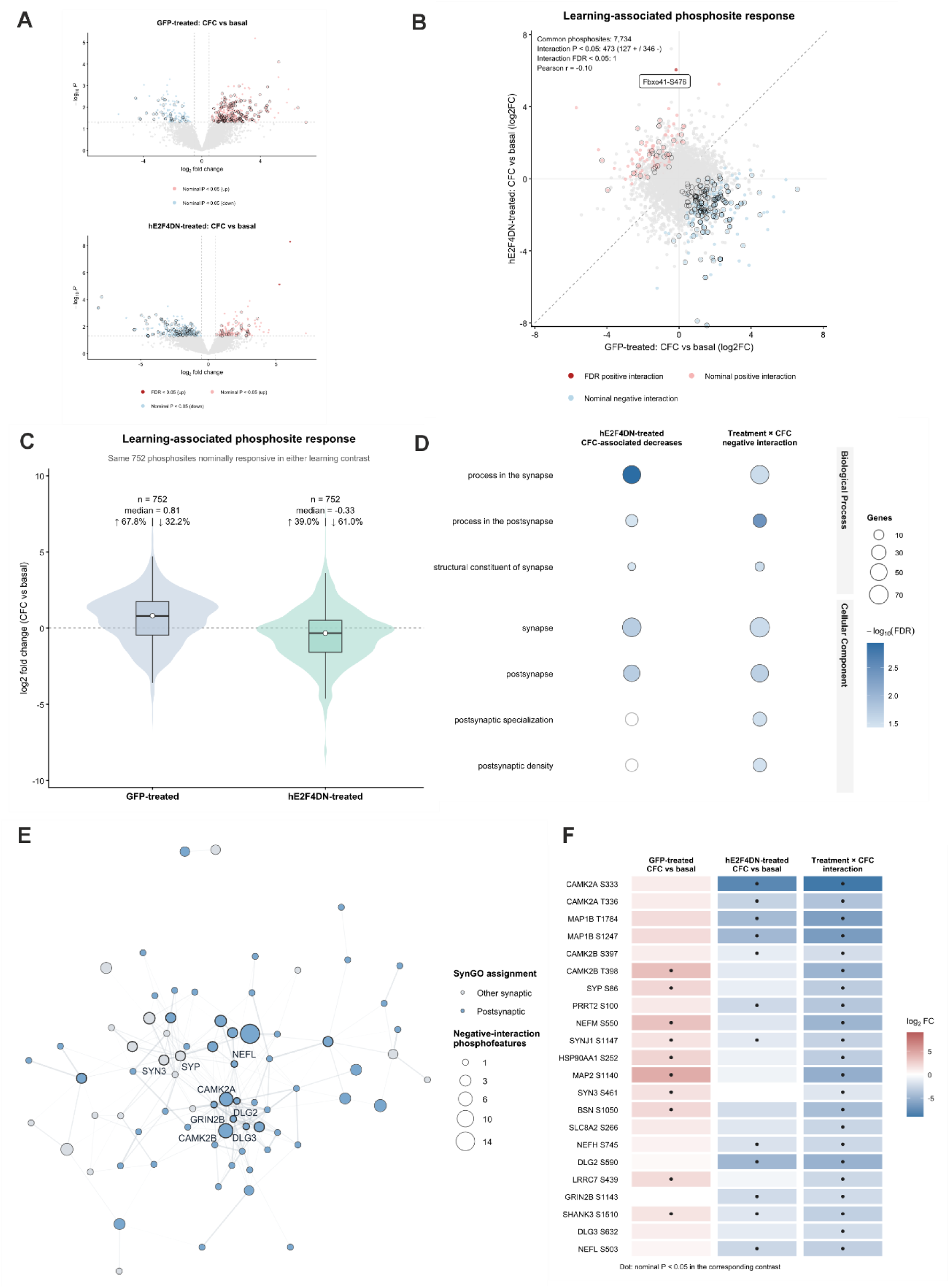
hE2F4DN reshapes learning-associated synaptic phosphoregulation. Phosphoproteomic analysis was performed on hippocampal samples from GFP- and hE2F4DN-treated h5xFAD mice under basal conditions or following contextual fear conditioning (CFC). **(A)** Volcano plots of differential phosphofeature abundance following CFC in GFP- and hE2F4DN-treated mice. In GFP-treated mice, 459 phosphofeatures showed nominal changes (340 increased and 119 decreased), compared with 394 in hE2F4DN-treated mice (138 increased and 256 decreased; *P* < 0.05). No phosphofeatures reached FDR < 0.05 in GFP-treated mice, whereas two were FDR-significant in hE2F4DN-treated mice: FBXO41-S476 (log₂FC = 6.05, FDR = 4.07 × 10⁻⁵) and VPS13A-S1929 (log₂FC = 5.25, FDR < 0.05). Dark red and blue points indicate FDR-significant features, lighter shading indicates nominal-only changes, and grey points indicate non-significant features. Black outlines highlight nominally altered phosphofeatures belonging to proteins with multiple responsive sites. Vertical lines at |log₂FC| = 0.5 are shown for reference only and were not used as significance thresholds. Complete differential phosphoproteomic results are provided in Table S10, and the corresponding total-protein measurements for the two FDR-significant sites are shown in Figure S19. **(B)** Comparison of learning-associated phosphoproteomic changes between treatments. Log₂ fold changes following CFC in hE2F4DN-treated mice are plotted against those observed in GFP-treated mice for 7,734 phosphofeatures evaluated in both learning contrasts and the formal interaction analysis. A total of 473 phosphofeatures showed nominal treatment-by-learning interactions (*P* < 0.05), comprising 127 positive and 346 negative effects. FBXO41-S476 was the only FDR-significant interaction phosphosite (log₂FC = 6.20, FDR = 0.0058). Point colors distinguish interaction direction and significance, whereas black outlines identify nominally interacting phosphofeatures from proteins exhibiting multisite responses. The dashed diagonal indicates equivalent learning-associated changes between treatments. The corresponding interaction volcano plot is provided in Figure S20. **(C)** Directionality of learning-associated phosphoproteomic changes. The same 752 phosphofeatures showing nominal CFC responsiveness in at least one treatment group were evaluated in both learning contrasts. GFP-treated mice exhibited predominantly positive changes (median log₂FC = 0.81; 67.8% positive), whereas hE2F4DN-treated mice showed predominantly negative changes (median log₂FC = −0.33; 61.0% negative). The corresponding analysis of all 7,734 commonly tested phosphofeatures is provided in Figure S21. **(D)** Comparative SynGO over-representation analysis of phosphofeatures decreased following CFC in hE2F4DN-treated mice and those exhibiting nominally negative treatment-by-learning interactions. Mouse genes were mapped to human orthologs, and contrast-specific tested genes were used as backgrounds. Terms reaching FDR < 0.05 in either comparison are displayed. Enriched categories included synaptic and postsynaptic processes and compartments, including postsynaptic specialization and postsynaptic density. Point size represents the number of foreground genes, color intensity indicates −log₁₀(FDR), and open grey circles indicate terms not reaching FDR < 0.05 in the corresponding comparison. Complete SynGO enrichment results are provided in Table S11. **(E)** STRING protein-association network of SynGO-associated proteins contributing to the negative treatment-by-learning interaction. Of 85 input proteins, 75 showed at least one association at a combined score ≥ 0.40, and 73 belonged to the largest connected component. No additional interactors were introduced. Node size represents the number of negative-interaction phosphofeatures per protein, while node fill distinguishes postsynaptic from other synaptic SynGO assignments. STRING-isolated proteins are not displayed. Prominent network members included CAMK2A, CAMK2B, GRIN2B, SYP, DLG2, DLG3, SHANK3 and MAP1B. **(F)** Representative phosphosite-level changes across the STRING-associated network, comprising 22 phosphosites from 19 proteins selected according to network connectivity, multisite recurrence and interaction evidence. Heatmap colors represent log₂ fold changes in the GFP-treated CFC versus basal, hE2F4DN-treated CFC versus basal, and formal treatment-by-learning interaction contrasts. Black dots indicate nominal *P* < 0.05 for the corresponding phosphosite and contrast. Complete network and phosphosite-selection data are provided in Table S12, and the corresponding total-protein abundance changes are shown in Figure S22.

SynGO over-representation analysis identified enrichment of predominantly postsynaptic processes and compartments among phosphofeatures showing negative learning-associated changes or negative treatment-by-learning interactions, particularly postsynaptic specializations and the postsynaptic density (Figure 7D). STRING protein-association analysis revealed an interconnected network including CaMKIIα/CaMKIIβ (*Camk2a*/*b*), GluN2B (*Grin2b*), Synaptophysin (*Syp*), PSD-93/SAP102 (*Dlg2*/*3*), and SHANK3 (*Shank3*) (Figure 7E). Representative phosphosites illustrated this coordinated response (Figure 7F), which was largely not accompanied by comparable changes in total-protein abundance (Figure S22).

*Overall, these results highlight distinct learning-associated phosphoregulation following E2F4DN treatment, involving selective phosphorylation changes in FBXO41 and VPS13A alongside broader remodeling enriched in postsynaptic processes*.

### hE2F4DN reverses hippocampal-dependent memory loss in h5xFAD mice

To determine whether LTP restoration, suppression of neuronal death and the associated molecular changes translate into functional recovery, we evaluated hippocampal-dependent spatial memory using the NOL (Figure 8A) and CFC (Figure 8F) paradigms.

**FIGURE 8.**
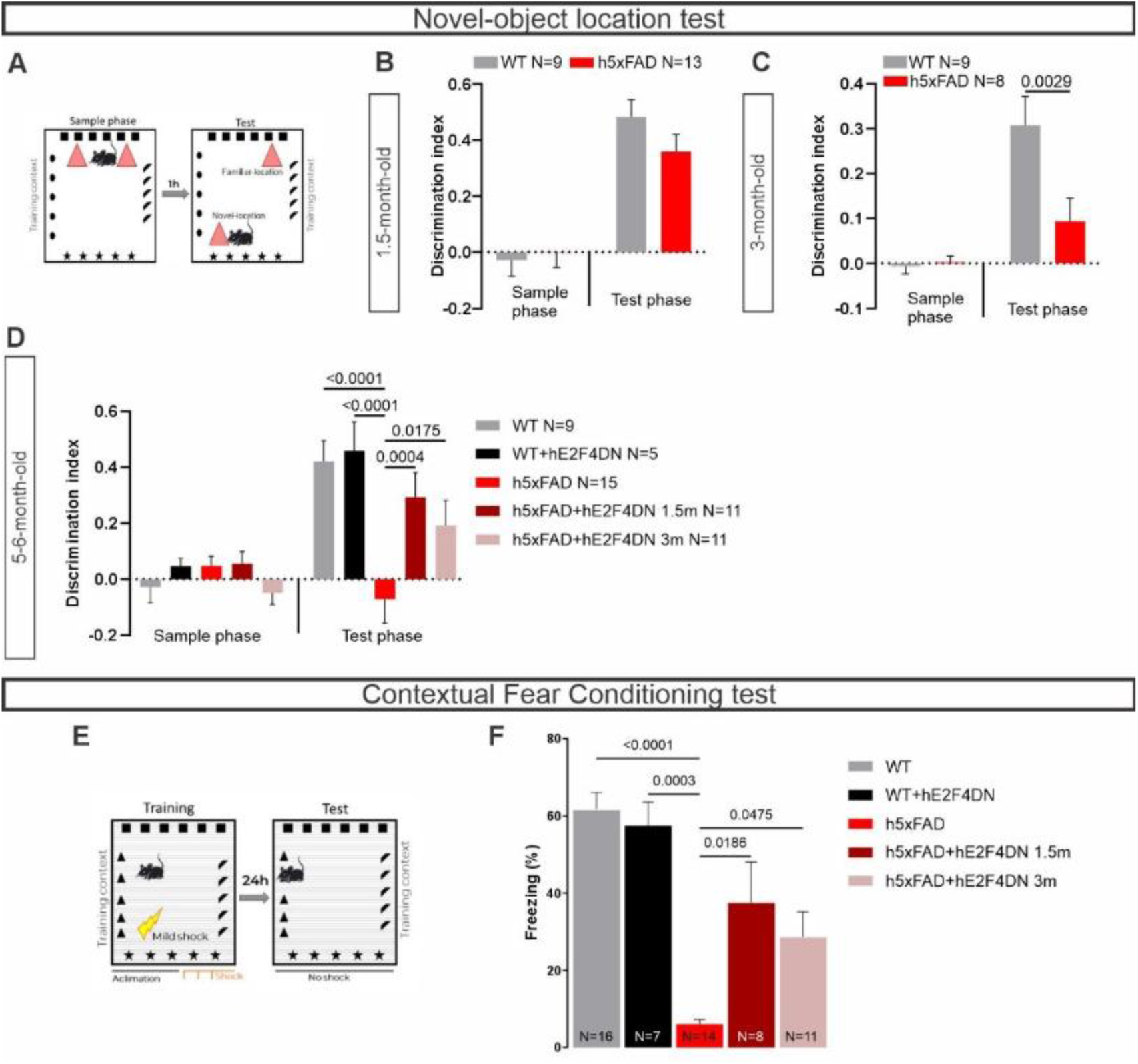
Intravenous administration of AAV.hE2F4DN recovers hippocampal-dependent memory in h5xFAD mice. Hippocampal-dependent memory was tested using the NOL **(A-E)** and CFC **(F, G)** tasks at 5-6 months of age in WT or h5xFAD mice, untreated or injected at 1.5 months (1.5m) or 3 months (3m) with the AAV.hE2F4DN vector (hE2F4DN). **(A)** Scheme of the NOL task. **(B-D)** Bar graphs illustrate the discrimination index in the sample and test phases by WT or h5xFAD mice of 1.5 months **(B)**, 3 months **(C)**, or 5-6 months **(D)**, treated as indicated. *P* values shown in the graphs were derived using Two way ANNOVA test [1.5 months: phase x group F (1, 20) = 1.508, P=0.2336; phase F (1, 20) = 50.71, P<0.0001; group F (1, 20) = 0.6776, P=0.4201]; [3 months: phase x group F (1, 15) = 6.533, P=0.0219; phase F (1, 15) = 21.50, P=0.0003; group F (1, 15) = 5.762; P=0.0298]; [6 months: phase x Group F (4, 46) = 5.583, P=0.0009; phase F (1, 46) = 25.46, P<0.0001; Group F (4, 46) = 4.455, P=0.0040] followed by Bonferroni’s multiple comparisons test for group comparison. **(E)** Scheme of CFC task. **(F)** Percentage of freezing to the context of the indicated experimental groups during the duration (5 min) of the CFC test, *P* values shown in the graphs were derived using Kruskal-Wallis test [(H(4) =33.50, P < 0.0001)], followed by Dunn’s multiple comparisons test. N represents number of animals.

In the NOL test, both untreated and hE2F4DN-treated WT mice exhibited intact hippocampus-dependent memory, with comparable discrimination index (DI) (Figure 8D), thus demonstrating that *hE2F4DN expression does not affect the formation of hippocampal-dependent memory.* In contrast, non-treated h5xFAD mice showed a significant impairment in the NOL task, with a DI close to 0, reflecting disrupted memory (Figure 8D). However, h5xFAD mice who had received the AAV.hE2F4DN vector at 1.5 months of age, a stage in which memory formation can still be observed (Figure 8B), displayed significantly higher DI (Figure 8D), indicating that the *neuronal expression of hE2F4DN prevents memory loss*, as memory was intact at the time of injection (Figure 8B). As an additional control, mice from all experimental groups spent similar time exploring both objects during the sample phase (Figure S4A-C).

At 3 months, consistent with the electrophysiological data, h5xFAD mice showed marked deficits in spatial memory (Figure 8C) despite normal exploratory behavior in the NOL test (Figure S4B). AAV.hE2F4DN treatment at 3 months restored memory performance when tested at 5–6 months (Figure 8D), paralleling the restoration of LTP.

Consistent with the results obtained in the NOL test, we did not observe changes at 5-6 months when WT mice treated with the AAV.hE2F4DN vector were compared to untreated WT mice in the CFC paradigm (Figure 8G). When h5xFAD mice were exposed to the CFC test, a significant increase in freezing behaviour could be observed in 1.5 and 3 months-injected h5xFAD mice, when compared to non-treated h5xFAD mice (Figure 8G).

*Thus, we found evidence of memory recovery induced by AAV.hE2F4DN administered after pathological damage and memory loss were established*.

## DISCUSSION

AD is characterized by progressive synaptic dysfunction, impaired neuronal plasticity, and ultimately neuronal loss, leading to the deterioration of learning and memory. Although considerable efforts have focused on preventing or slowing disease progression, restoring neuronal function once cognitive deficits are established remains a major therapeutic challenge. Here, we demonstrate that gene therapy based on hE2F4DN exerts both preventive and restorative effects in homozygous 5xFAD (h5xFAD) mice, an aggressive model of AD-like pathology. Administration of hE2F4DN at 1.5 months of age, before the onset of severe functional impairment, prevented the loss of hippocampal long-term potentiation (LTP), reduced amyloid-β (Aβ) pathology, and protected against neuronal death. This extends previous studies of our laboratory demonstrating that this early intervention is able to attenuate neuroinflammation and to prevent cell cycle reentry leading to neuronal tetraploidization (López-Sánchez et al., 2021), a process likely equivalent to neuronal senescence (Wong & Chow, 2023; Wu et al., 2024), and to recover adult neurogenesis (Lozano-Ureña et al., 2024). Moreover, our results confirm previous findings of our laboratory showing that hE2F4DN, when administered at 1.5 months, reduces Aβ pathology (López-Sánchez et al., 2021). Remarkably, administration at 3 months of age, when hippocampal LTP and learning and memory performance are already profoundly impaired, restored the capacity to generate LTP and was associated with reduced Aβ pathology and preservation of neuronal integrity. Importantly, both treatment paradigms resulted in preserved or recovered memory performance at 6 months of age, as demonstrated by NOL and CFC tests. These findings indicate that E2F4DN can not only interfere with the development of AD-like pathology but also promote functional recovery after substantial synaptic impairment has occurred.

Proteomic and phosphoproteomic analyses further revealed distinct molecular signatures associated with hE2F4DN treatment. Under basal conditions, hE2F4DN administration to h5xFAD mice was associated with metabolic remodeling, attenuation of neuroimmune and amyloid-related responses, and phosphoproteomic changes consistent with reduced GSK3β signaling and altered regulation of synaptic vesicle-associated processes that may underly the prevention by hE2F4DN of LTP deficits in h5xFAD mice. Following learning, h5xFAD mice treated with hE2F4DN at a stage when pathological deficits were already evident exhibited coordinated remodeling of presynaptic and postsynaptic molecular machinery, suggesting the re-engagement of pathways involved in activity-dependent synaptic plasticity. These changes may involve local translation of pre-existing mRNAs, potentially facilitating the rapid replenishment or reorganization of synaptic components involved in memory encoding. Moreover, increased phosphorylation of FBXO41 and VPS13A, proteins implicated in neuronal protein regulation and lipid homeostasis, respectively, points to additional pathways potentially contributing to synaptic recovery. Collectively, these findings suggest that hE2F4DN engages distinct, context-dependent molecular programs that may promote both neuroprotection and functional synaptic reorganization, even when administered after the onset of synaptic dysfunction.

Our findings demonstrate for the first time that the ability to induce LTP, the molecular substrate of memory, can be restored even after it has been lost in advanced AD stages in mice. Therefore, synaptic plasticity failure is not irreversible. Prior studies have shown that pharmacological interventions can enhance the amplitude of residual LTP in models with preserved, but abnormal, LTP-induction capacity (Gong et al., 2004; Hwang et al., 2017; Kong et al., 2023; Martinez-Hernandez et al., 2018; Rex et al., 2006; Vitolo et al., 2002; Cortese et al., 2018; Kauwe et al., 2024; Chaubey et al., 2026). However, hE2F4DN reinstated the ability to generate LTP at a disease stage when it had been lost, directly linking molecular rescue of synaptic plasticity with cognitive improvements. The ability of hE2F4DN to prevent neuronal loss enables sustained LTP recovery, which necessarily depends on the presence of viable neurons and synapses.

Mechanistically, the synaptic localization of hE2F4DN within dendritic spines (Figure S2F, G) supports a direct role of E2F4 in synaptic regulation and contrasts with the deleterious activity of its E2F-family member E2F1, which increases with age in dendritic spines (Ting et al., 2014) and promotes cognitive decline (López-Sánchez et al., 2017). This would be another example of the opposing actions of these transcription factors on different functions (Crosby & Almasan, 2004).

In the early-treatment cohort analyzed under basal conditions, proteomic remodeling involved proteins associated with neuronal signaling, structural regulation, and Ca²⁺ homeostasis. The latter is particularly relevant, as neuronal calcium dyshomeostasis has been implicated in AD pathogenesis. Indeed, impaired calcium homeostasis has been reported in hippocampal neurons of 5xFAD mice even before the onset of amyloid plaque accumulation (Skobeleva et al., 2022). Aβ has also been shown to disrupt neuronal calcium regulation, leading to elevated intracellular Ca²⁺ levels, metabolic dysfunction, and apoptosis, which may contribute to cognitive decline (Ge et al., 2022). At later stages, amyloid plaques further disrupt neuritic calcium homeostasis, as demonstrated in vivo in an AD mouse model, potentially compromising neuronal network function (Kuchibhotla et al., 2008). In this context, the proteomic changes induced by hE2F4DN may reflect a reorganization of molecular mechanisms involved in neuronal calcium signaling and ionic homeostasis, both essential for maintaining neuronal excitability and function. In the early-treatment cohort analyzed under basal conditions, proteomic remodeling involved proteins associated with neuronal signaling, structural regulation, and Ca²⁺ homeostasis. The latter is particularly relevant, as neuronal calcium dyshomeostasis has been implicated in AD pathogenesis. For instance, it has been described to be impaired in 5xFAD mice hippocampal neurons, preceding amyloid plaque accumulation (Skobeleva et al., 2022). Importantly, Aβ upregulates calcium in neurons, leading to an increase in intracellular calcium concentration, which results into abnormal neuronal metabolism and apoptosis, causing memory decline (Ge et al., 2022). Additionally, in later stages senile plaques impair neuritic calcium homeostasis, as shown in vivo in AD mouse model, disrupting neuronal networks (Kuchibhotla et al., 2008). Thus, given the alterations in calcium homeostasis and neuronal excitability reported in AD, hE2F4DN-modulation may reflect a reorganization of mechanisms regulating neuronal calcium signaling and ionic homeostasis, essential for neuron excitability and homeostasis.

Phosphoproteomic analysis highlighted synaptic-vesicle regulation and coordinated changes in neuronal phosphosignaling upon hE2F4DN treatment at 1.5 months. These steady-state effects were relatively modest at the level of synaptic protein abundance, suggesting that hE2F4DN may prime synaptic regulatory mechanisms that become more apparent in response to learning-related neuronal activity.

Neuroimmune remodeling represented another prominent component of the early treatment response confirming the attenuation of reactive astrocytosis and microgliosis in h5xFAD mice (López-Sánchez et al., 2021), and paralleled the reduction in amyloid pathology and neuronal damage. Proteomic analysis revealed treatment-associated changes in inflammatory and glial programs, together with pathways involved in phagocytosis, endolysosomal trafficking, and Aβ clearance. Rather than indicating a general suppression of neuroinflammation, these findings suggest a reorganization of glial and immune responses that may favor Aβ homeostasis by facilitating its uptake, intracellular trafficking, and degradation. This interpretation is consistent with the observed reduction in soluble Aβ oligomers and amyloid plaque area following hE2F4DN treatment. Given the well-established inhibitory effects of soluble Aβ oligomers on synaptic plasticity (Li et al., 2011; Knafo et al., 2016), their reduction may, in turn, contribute to the restoration of hippocampal LTP.

Metabolic and redox remodeling further supported enhanced cellular resilience. Proteomic analyses identified treatment-associated changes in lipid and sterol metabolism, redox homeostasis and mitochondrial processes. Altered brain cholesterol homeostasis is increasingly recognized as a feature of AD, with excess neuronal cholesterol and dysregulated cholesterol biosynthesis, transport and clearance contributing to amyloidogenic processing and neuronal dysfunction (Varma et al., 2021;

Ahmed et al., 2024). Increased astrocytic cholesterol synthesis has also been linked to Aβ-driven dysfunction in 5xFAD mice (Zhang et al., 2026). Thus, the opposite shift of cholesterol/sterol-associated programs following hE2F4DN treatment is consistent with partial correction of an AD-associated lipid-homeostasis disturbance. Metabolic remodeling was not restricted to lipid pathways but also extended to mitochondrial and redox processes. In particular, changes involving NADH dehydrogenase complex assembly, together with increased levels of MTHFD1L, GSR, BLVRA and LONP1 (Pike et al., 2010; Barone et al., 2011; Wang et al., 2023; Yu et al., 2024), further implicate mitochondrial metabolism, antioxidant defense and proteostasis in the treatment response. Together, these changes are consistent with a metabolic and redox environment more favourable to neuronal resilience and survival, in line with the preservation of neuronal populations observed following hE2F4DN treatment.

The early-treatment phosphoproteomic analysis additionally identified a coordinated shift in GSK3β-associated substrate phosphorylation. The reduction in GSK3β-associated phosphorylation following hE2F4DN treatment, including at several MAPT sites, is consistent with a potentially beneficial shift in phosphorylation patterns, given the established involvement of GSK3β in tau hyperphosphorylation, synaptic dysfunction, and neuronal pathology (Lucas et al., 2001; Hooper et al., 2008; Sayas and Ávila, 2021). Increased biliverdin reductase A (Blvra) expression may further support this interpretation, as this enzyme promotes Akt-mediated inhibition of GSK3β (Sharma et al., 2019). Although phosphopeptide coverage after tryptic digestion did not include several canonical pathological tau phospho-epitopes, the coordinated decrease across multiple GSK3β-associated phosphosites nevertheless supports broader remodeling of GSK3β-linked phosphosignaling. Importantly, KSEA does not directly measure kinase catalytic activity; these findings should therefore be interpreted as coordinated changes in GSK3β-associated substrate phosphorylation rather than direct evidence of GSK3β inhibition. Nevertheless, we provide evidence that hE2F4DN reduces the levels of a pathological form of phosphorylated tau in the hippocampus of h5xFAD mice. Increased tau phosphorylation at Ser396 has been reported in patients with AD (Regalado-Reyes et al., 2019) and in mouse models of AD (Kanno et al., 2014; Li et al., 2025). Moreover, phosphorylation at this residue has been detected in paired helical filaments (PHFs), the major constituents of neurofibrillary tangles in AD (Goedert et al., 1994). Tau phosphorylation at Ser396 has also been associated with synaptic weakening (Mitchell et al., 2025) and has been shown to precede the onset of spatial learning and memory deficits, remaining elevated at 6 months of age in 5xFAD mice (Kanno et al., 2014). Thus, the reduction in GSK3β activity and tau phosphorylation may contribute to the improvements in synaptic function and learning observed in hE2F4DN-expressing mice.

While these early-treatment analyses describe the molecular state associated with long-term protection, they do not address how neuronal hE2F4DN expression shapes the hippocampal molecular response to a learning task in h5xFAD mice with pre-existing cognitive impairment. The late-treatment learning cohort provided an opportunity to examine this question directly. Although individual protein changes 1 h after CFC were modest, preranked GSEA using SynGO gene sets revealed broad enrichment of pre- and postsynaptic programs in E2F4DN-treated mice, involving glutamatergic receptor components, synaptic-vesicle release machinery and proteins controlling synaptic organization. To further explore the relationship between the learning-associated proteomic response and dendritic mRNA localization, we compared our dataset with the repertoire of dendritic mRNAs reported by Ainsley et al. (2014) to rapidly associate with ribosomes following CFC. Of the 1,874 genes associated with dendritic ribosome-bound mRNAs in that study, 1,204 (64.2%) were represented in our hippocampal proteomic dataset. Among these, 57 proteins exhibited nominal changes in abundance following CFC in E2F4DN-treated mice, with 37 showing increased abundance (Supplementary Figure S23; Supplementary Table S12). Although this overlap does not provide direct evidence of local translation, it identifies learning-responsive proteins encoded by transcripts previously associated with dendritic ribosomes. This activity-dependent reorganization was even more apparent at the phosphoproteomic level. The post-CFC response showed contrasting directional patterns between treatments, with a predominantly positive phosphorylation-associated response in GFP-treated h5xFAD mice and a predominantly negative response following E2F4DN treatment. Interestingly, Heo et al. (2023) reported widespread decreases in postsynaptic phosphorylation following associative learning, indicating that reduced phosphorylation at selected sites can accompany experience-dependent synaptic remodeling Although the experimental paradigms and tissue preparations differ, our results similarly emphasize the importance of dynamic phosphoregulation rather than generalized phosphorylation activation or suppression. Interestingly, treatment-by-learning interactions were particularly associated with postsynaptic processes and an interconnected network involving CAMK2α/β (*Camk2a*/*b*), GluN2B (*Grin2B*), PSD-93/SAP102 (*Dlg2*/*3*), SHANK3 (*Shank3*), Synaptophysin (*Syp*), and Microtubule-associated protein 1B (*Map1b*), linking glutamatergic signaling, vesicle regulation, postsynaptic organization, and structural plasticity. Taken together, these findings suggest that E2F4DN reshapes the hippocampal phosphorylation response to learning in cognitively impaired h5xFAD mice, particularly within an interconnected synaptic network involved in glutamatergic signaling, synaptic-vesicle function, postsynaptic organization, and structural plasticity, rather than simply inducing a global increase or decrease in phosphorylation.

Two individual phosphosites provided additional insight into this activity-dependent remodeling. FBXO41-S476 and VPS13A-S1929 showed pronounced CFC-associated increases in hE2F4DN-treated mice without corresponding changes in total-protein abundance, with FBXO41-S476 also exhibiting the only FDR-significant treatment-by-learning interaction (Figure 7A,B; Figure S19). FBXO41 is a neuronal F-box protein involved in synaptic transmission, whose loss reduces synaptic vesicle release probability in hippocampal excitatory neurons (Quadros et al., 2022). VPS13A mediates interorganelle lipid transport (Kumar et al., 2018), and its depletion has been associated with impaired corticostriatal long-term depression (García-García et al., 2023) and altered lipid-dependent PKC signaling and dendritic spine integrity (Besa-Selva et al., 2026). These findings highlight two potentially distinct components of activity-dependent synaptic regulation, involving neurotransmitter release and lipid-dependent synaptic organization.

The hE2F4DN-associated response should not be interpreted simply as greater molecular activation: the proteomic and phosphoproteomic datasets instead point to a distinct organization of the response to learning. Because hippocampi were collected 1 h after CFC, these changes are most consistent with molecular events occurring during early memory consolidation and post-learning synaptic remodeling rather than with the initial induction of LTP itself.

Overall, our observations suggest that hE2F4DN may act at two complementary levels: preserving a cellular environment conductive to homeostasis, neuronal survival, and synaptic function, while also reshaping the hippocampal molecular response to learning. These findings identify hE2F4DN as a promising therapeutic strategy with the potential to restore molecular processes underlying learning and memory, even in the context of established AD-related pathology and cognitive impairment. If successfully translated to humans, hE2F4DN could offer a novel therapeutic approach aimed at limiting neurodegeneration and restoring synaptic function, with the potential to improve cognition in patients with moderate dementia due to AD, for whom curative treatments remain unavailable.

## AUTHOR CONTRIBUTIONS

Conceptualization: Cristina Sánchez-Puelles, José María Frade. Methodology: Cristina Sánchez-Puelles, Alice Grieco, Ana Montero-Calle, Rodrigo Barderas, José María Frade. Investigation: Cristina Sánchez-Puelles, Alicia Hernández-Vivanco, Alice Grieco, Ana Montero-Calle, Rodrigo Barderas, Noelia López-Sánchez, Alberto Fraj-Cebrián, Irene Camacho-Olmos, Alberto Garrido-García, Vanesa Cano-Daganzo, Lorena Valdés-Lora. Supervision: Cristina Sánchez-Puelles, José María Frade. Writing – original draft: Cristina Sánchez-Puelles, Alice Grieco, José María Frade. Writing – review & editing: Cristina Sánchez-Puelles, Alice Grieco, Ana Montero-Calle, Rodrigo Barderas, Gertrudis Perea, Álvaro Pascual-Leone, José María Frade.

## ACKNOWLEDGMENTS

We acknowledge J. DeFelipe and P. Merino-Serrais for their early discussions and for providing the Stereoinvestigator system. We thank P. Mendez for providing the CFC system. We are grateful to E. Boyden (MIT), L.-H. Tsai (MIT), C. Barnes (University of Arizona), and M. Albert (Johns Hopkins University) for their critical reading of the manuscript and their scientific advice. We thank Juan Vicente Torres for his review of the statistical analyses.

