## Supplementary Material for "E2F4 gene therapy reverses established synaptic and cognitive deficits and remodels learning-associated signaling in Alzheimer’s disease mice"

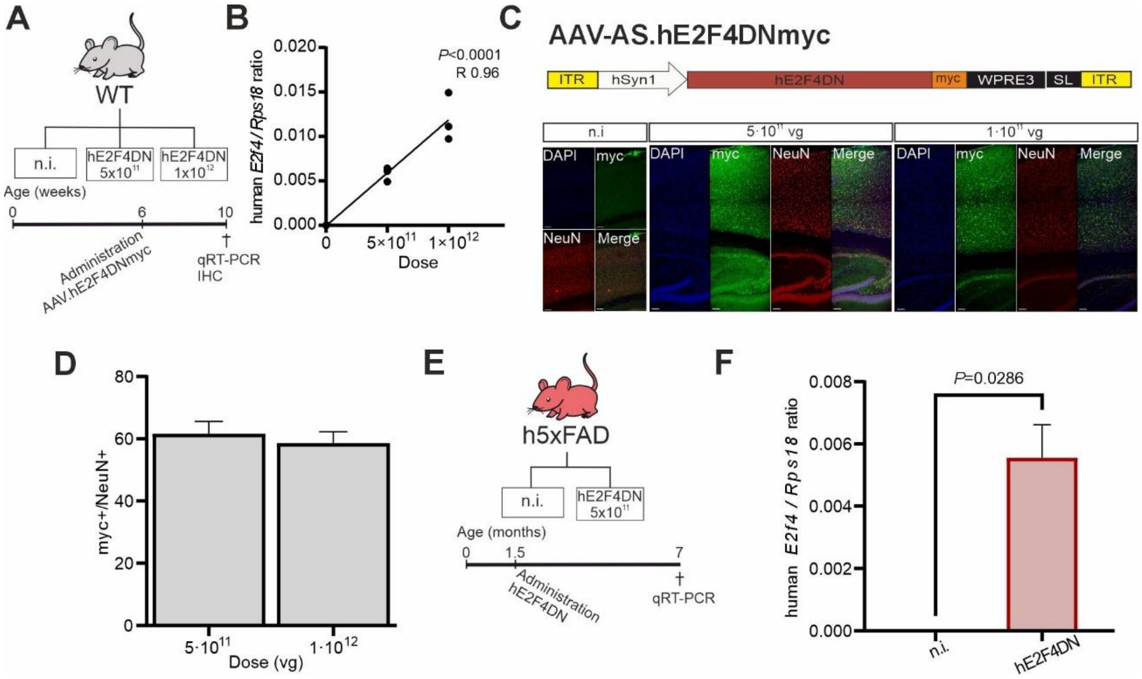


**FIGURE S1. AAV.hE2F4DN characterization. (A)** Scheme of the experimental design using WT mice (doses and the time course of the treatments). **(B)** Ratio between *hE2f4dn* (human *E2F4*) and *Rps18* gene expression, quantified by qRT-PCR from cDNA obtained from the cerebral cortex of WT mice injected at 1.5 months of age with the indicated doses of hE2F4DN vector and sacrificed one month later (Pearson's R correlation test). **(C)** Top: Scheme showing the genomic sequence of the AAV-AS.hSyn1-hE2F4myc-WPR3SL vector (AAV-AS.hE2F4DNmyc). Bottom: Representative images of the cerebral cortex and hippocampus from WT mice administered at 1.5 months of age with the AAV.hE2F4DNmyc vector at the indicated doses and sacrificed one month later. Blue: DAPI. Green: anti-Myc tag. Red: NeuN. Scale bar: 100 µm. **(D)** Percentage of NeuN$^{+}$ cortical neurons expressing hE2F4DN-myc one month after administration of the hE2F4DN vector at the indicated doses. **(E)** Scheme of the experimental design using h5xFAD mice (doses and the time course of the treatments). **(F)** Ratio between *hE2f4dn* (human *E2F4*) and *Rps18* gene expression, quantified by qRT-PCR from cDNA obtained from the cerebral cortex of h5xFAD mice injected at 1.5 months of age with the hE2F4DN vector ($5{\times}10^{11}$vg) and sacrificed at 7 months of age (*P* value was determined using a Mann–Whitney test).

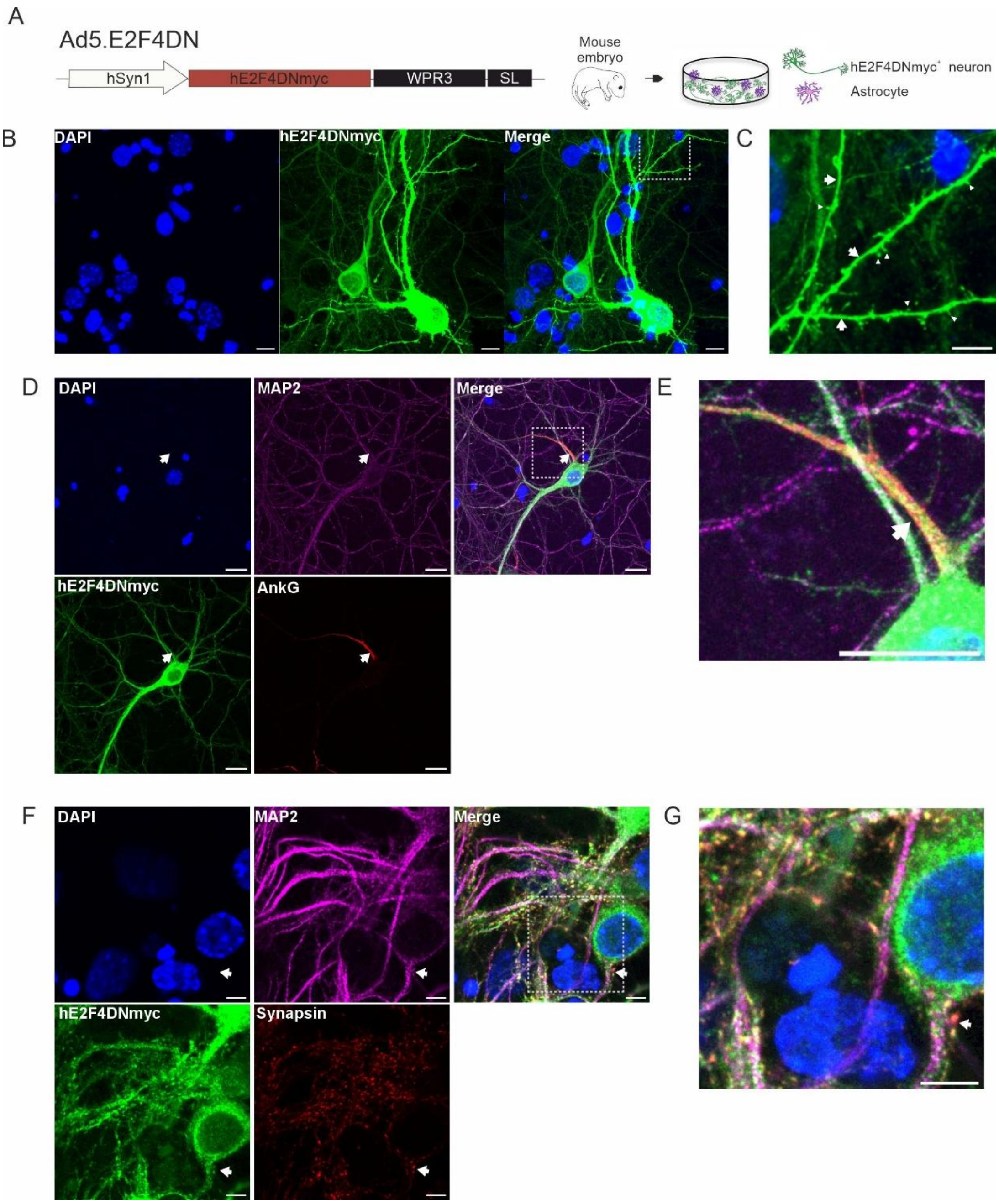


**FIGURE S2. hE2F4DNmyc expression in dendritic spines. (A)** Scheme describing the genomic sequence of the Ad5.hE2F4DNmyc vector (Ad5.E2F4DN) used to transduced mixed hippocampal primary cultures. White: hSyn1 promoter. Dark red: hE2f4DN sequence with a myc tag. Black: WPRE3SL sequence. **(B)** Confocal projection showing hE2F4DNmyc expression (Blue: DAPI; green: myc). **(C)** High magnification of the dotted square in panel B illustrating dendrites (arrows) and dendritic spines (arrowheads). **(D)** Confocal projection showing hE2F4DNmyc expression in dendrites and axons of hippocampal neurons. Dendrites are revealed as MAP2+ processes. The axon initial segment (arrow) expressing Ankyrin G (AnkG) identifies the proximal part of the axon. Blue: DAPI; green: myc; red: AnkG; magenta: MAP2. **(E)** High magnification of the dotted square (merge) in panel **D** illustrating axon initial segment (arrow). **(F)** Confocal projection showing hE2F4DNmyc expression in synapses. Notice proximal localization of hE2F4DNmyc and synapsin (arrows). Blue: DAPI; green: myc; red: synapsin; magenta: MAP2. **(G)** High magnification of the dotted square in panel **F** illustrating spines (arrow) with colocalization of synapsin and hE2F4DN. Scale bars: 10 µm **(G)**, 5 µm **(B, E, F)**, 20 µm **(C, D)**.

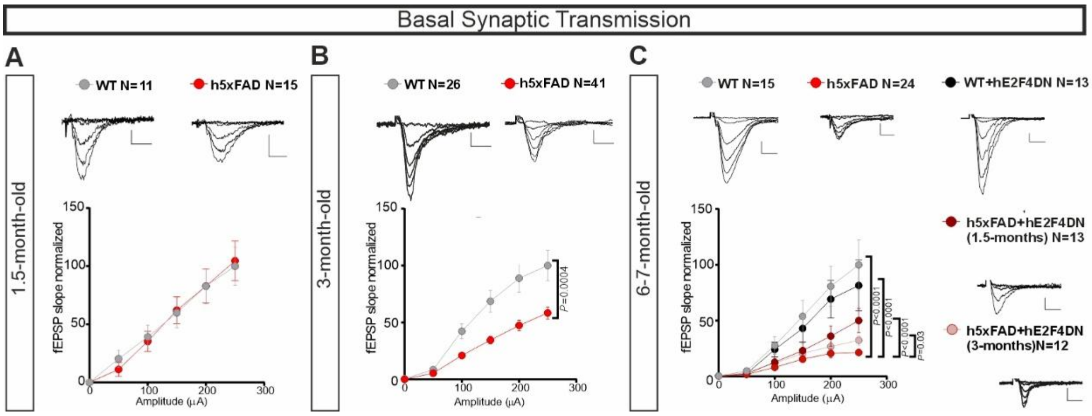


**FIGURE S3. Intravenous administration of AAV.hE2F4DN partially rescues basal synaptic transmission in h5xFAD mice.** Input–output curves of fEPSPs evoked in hippocampal coronal slices by stimulation of Schaffer collateral fibers and recorded in the CA1 region of untreated WT and h5xFAD mice of 1.5 months **(A)** or 3 months. *P* value was determined using two-way ANOVA (F (1, 65) = 13,77) , followed by Bonferroni post-hoc test **(B)**. **(C)** Input–output curves of fEPSPs evoked and recorded as in **A-B** of 6-7-month-old mice of the indicated genotypes (WT or h5xFAD), either non-treated or administered with AAV.hE2F4DN (hE2F4DN) at 1.5 or 3 months of age. Top: Representative traces of fEPSPs superimposed at 10, 50, 100, 150, 200 and 250 μA of stimulation intensity. Bottom: Input-output curve, *P* values were determined with a two-way ANOVA (F (2,305, 479,5) = 21,52), followed by Bonferroni post-hoc test. N represents number of slices from 5 WT and 8 h5xFAD untreated 1.5-month-old mice **(A)**, 11 WT and 17 h5xFAD untreated 3-month-old mice **(B)**; and the following untreated and treated 6-7 months old mice: 9 WT, 7 WT+hE2F4DN, 11 h5xFAD mice, 8 h5FAD+hE2F4DN treated at 1.5 months and 6 h5FAD+hE2F4DN treated at 3 months old mice **(C)**.

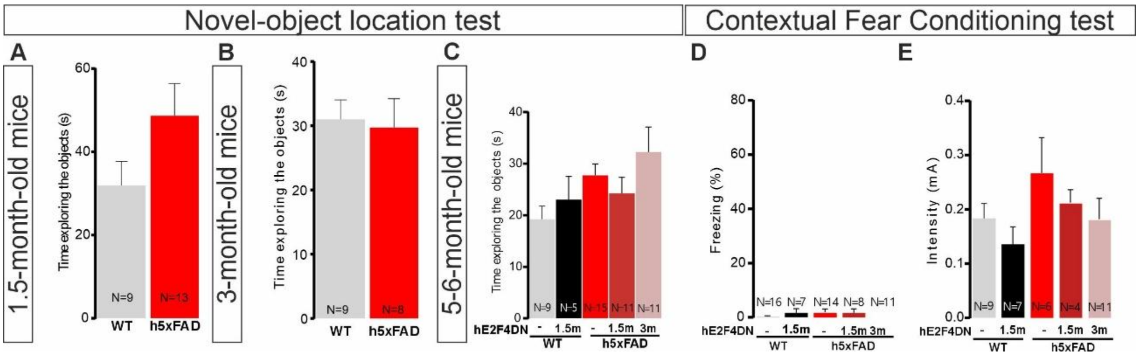


**FIGURE S4. Intravenous injection of AAV.hE2F4DN in h5xFAD mice recovers hippocampal-dependent memory (control results). (A-E)** Control results in the NOL test performed in WT and h5xFAD mice of 1.5 months and 3 months, and in 5-6-month-old WT and h5xFAD mice untreated or injected with the AAV.hE2F4DN viral vector at 1.5 months (1.5m) or 3 months (3m) (h5xFAD). Time exploring both objects in the test phase in 1.5 month-old **(A)**, 3-month-old **(B)** and 5-6 month-old **(C)** mice. **(D, E)** Control results in the CFC test performed in WT and h5xFAD mice of 5-6 months untreated or injected with the AAV.hE2F4DN viral vector at 1.5 month (1.5m) or 3 month (3m) (h5xFAD). **(D)** Percentage of freezing displayed by WT mice (control and injected at 1.5 month with AAV.hE2F4DN) and h5xFAD mice (control and injected with AAV.hE2F4DN at 1.5 and 3 month of age) during the first 3 minutes of CFC training, before shock exposure. **(E)** Pain threshold evaluated as the intensity in which the indicated animals vocalized or jumped. N: number of mice analyzed.

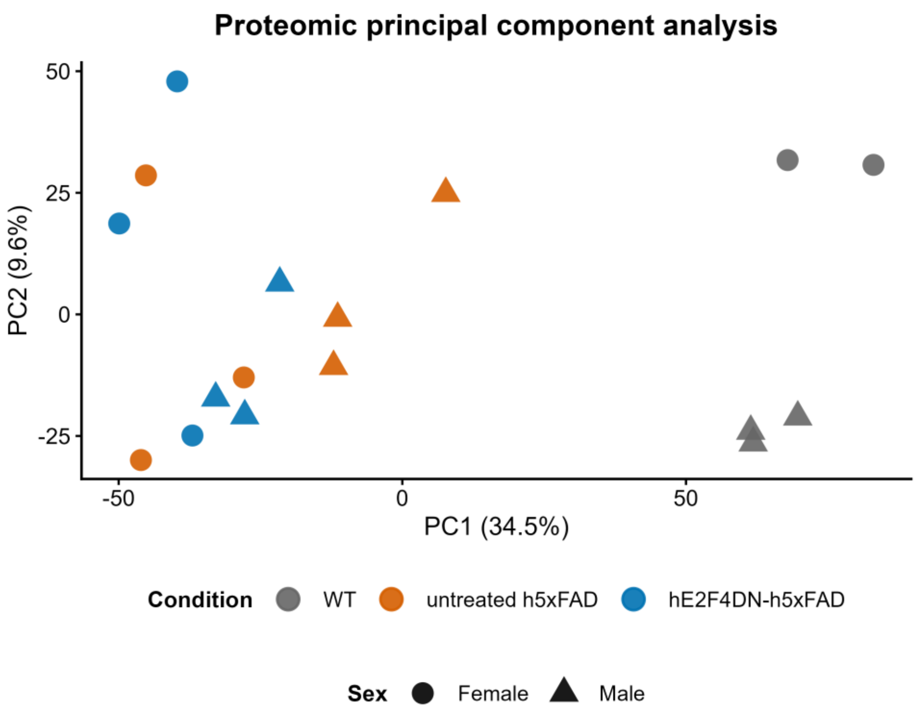


**FIGURE S5. Principal component analysis of the hippocampal proteomic dataset.** Principal component analysis (PCA) of quantitative hippocampal protein-abundance profiles from WT ($n$ = 5), untreated h5xFAD ($n$ = 6) and hE2F4DN-treated h5xFAD ($n$ = 6) mice. Each point represents one biological sample; colors indicate experimental group and point shapes indicate sex. The percentage of total variance explained by each principal component is indicated on the corresponding axis. Global proteomic variation was primarily associated with disease genotype, whereas untreated and hE2F4DN-treated h5xFAD samples showed substantial overlap in principal-component space.

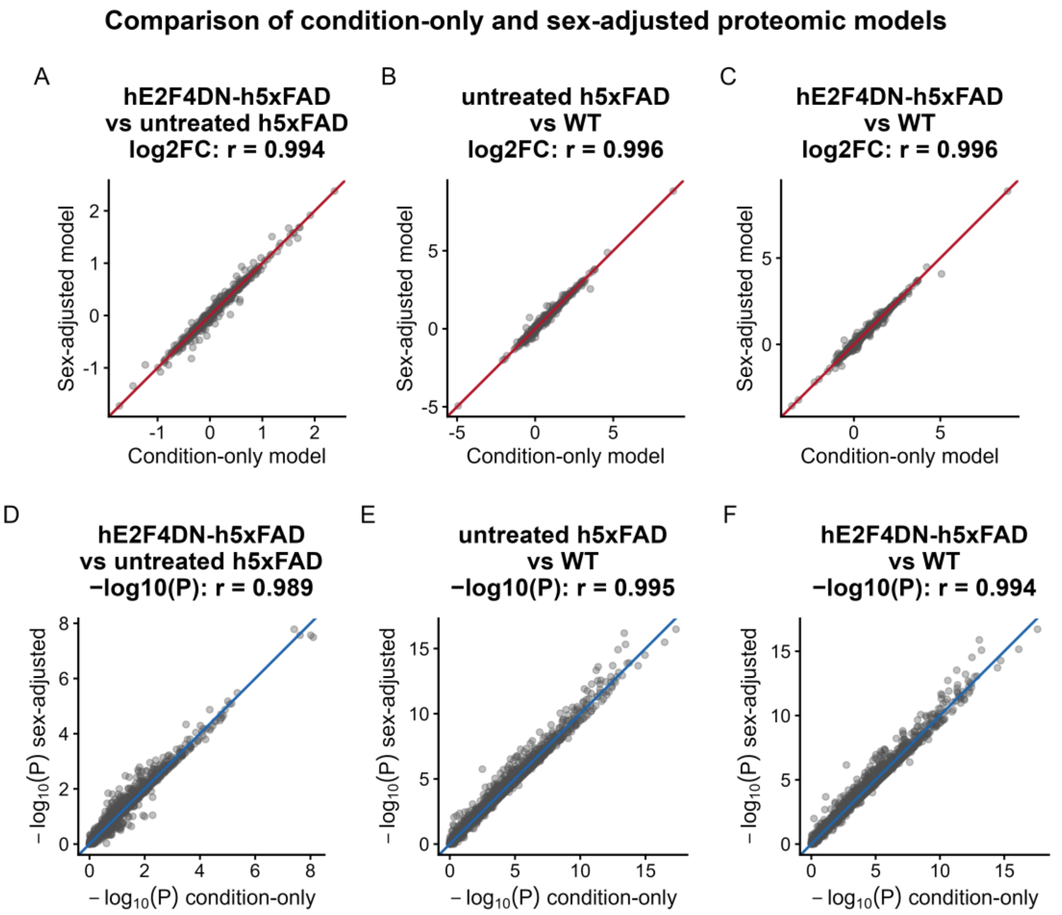


**FIGURE S6. Comparison of condition-only and sex-adjusted proteomic differential-abundance models.** Sensitivity analysis comparing protein-level differential-abundance estimates obtained using condition-only and sex-adjusted *limma* models. **(A–C)** Correlation of $\log_2$ fold-change estimates for hE2F4DN-treated h5xFAD versus untreated h5xFAD, untreated h5xFAD versus WT, and hE2F4DN-treated h5xFAD versus WT, respectively. **(D–F)** Corresponding comparison of $-\log_{10}(P)$ values for the same contrasts. Each point represents one quantified protein. Diagonal lines indicate identity between models, and Pearson correlation coefficients ($r$) are shown in each panel. The high concordance between models indicates that adjustment for sex had minimal influence on the overall differential-abundance estimates.

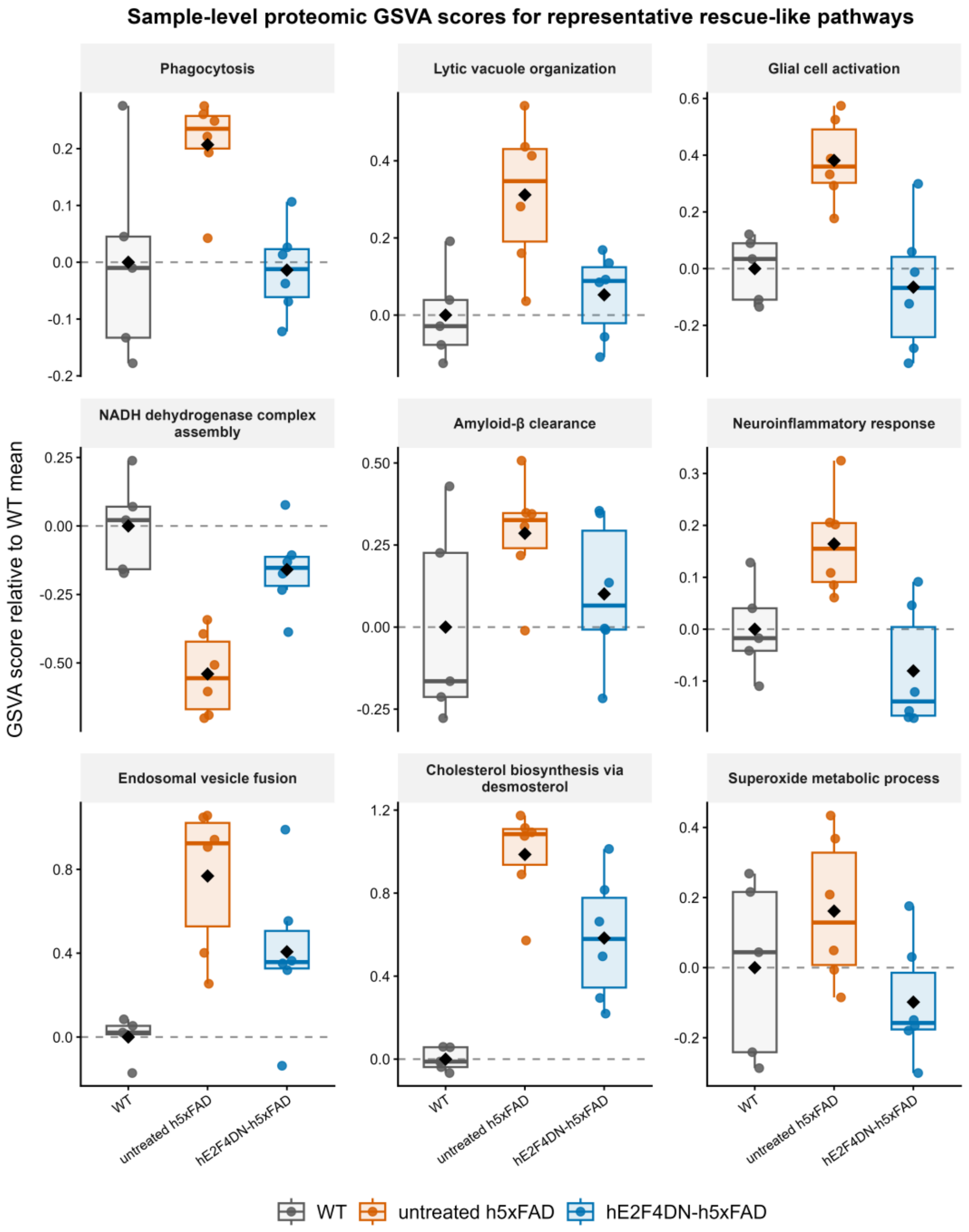


**FIGURE S7. Sample-level GSVA profiles of representative proteomic rescue-like pathways.** Sample-level GSVA scores for representative pathways displayed in Fig. 4E and selected among pathways showing rescue-like comparative GSEA behavior together with a treatment-associated shift toward WT by GSVA. Representative pathways encompass neuroimmune and inflammatory responses, amyloid-β handling and endolysosomal trafficking, lipid and sterol metabolism, redox regulation and mitochondrial function. For each pathway, GSVA scores were centered to the WT group mean. Individual points represent biological samples, boxplots summarize group distributions, and black diamonds indicate group means.

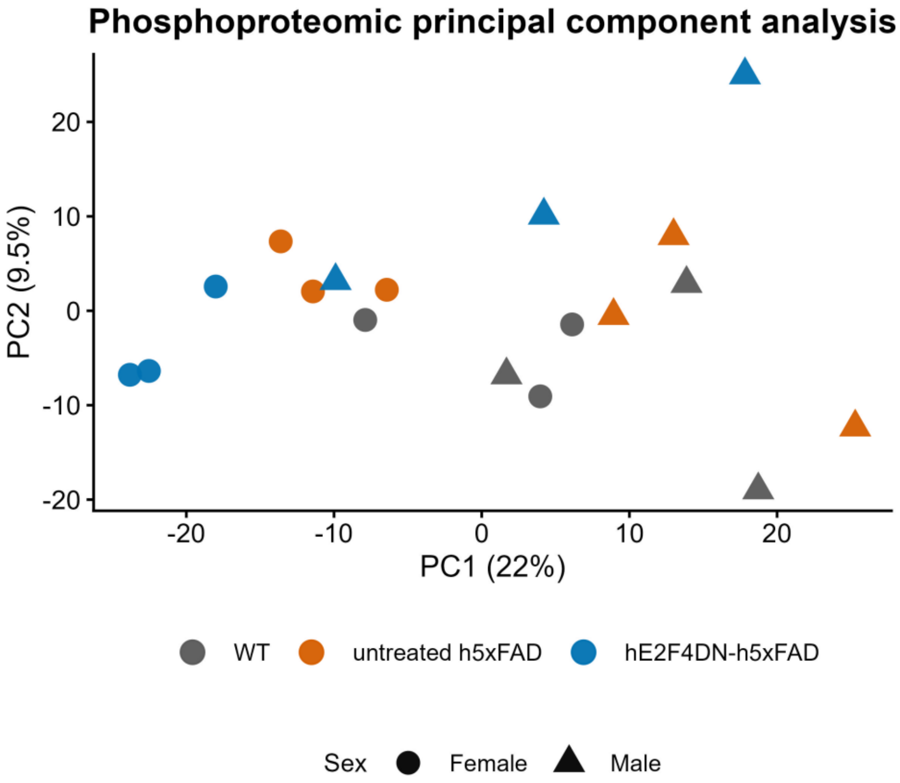


**FIGURE S8. Principal component analysis of the hippocampal phosphoproteomic dataset.** Principal component analysis (PCA) of hippocampal phosphoproteomic profiles from WT, untreated h5xFAD and hE2F4DN-treated h5xFAD mice (*n* = 6 per group). For PCA visualization only, missing values were temporarily replaced by the median abundance of the corresponding phosphosite-associated feature, after which the 1,000 most variable features were centered and scaled. Imputed values were not used for differential phosphorylation, Gene Ontology enrichment or kinase–substrate enrichment analyses. Each point represents one biological sample; colors indicate experimental group and point shapes indicate sex. The percentage of total variance explained by each principal component is indicated on the corresponding axis. No clear separation between experimental groups was observed in principal-component space.

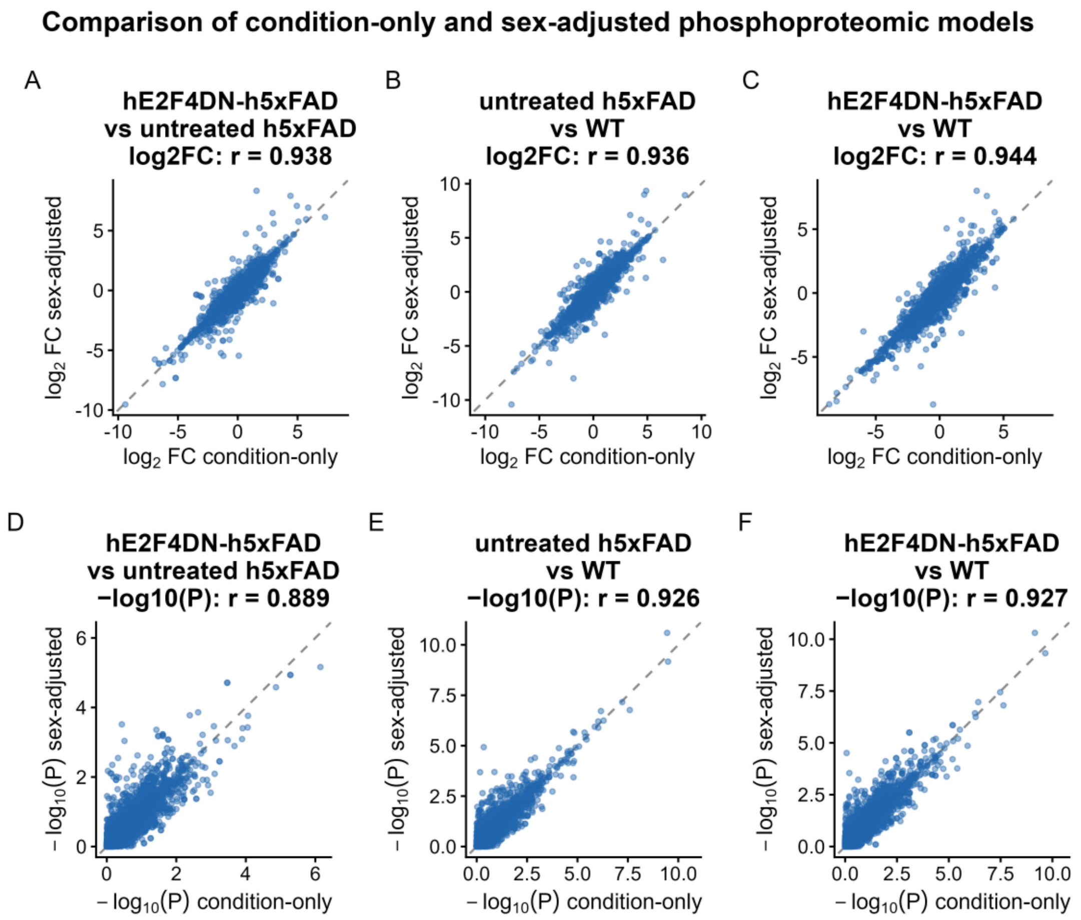


**FIGURE S9. Comparison of condition-only and sex-adjusted phosphoproteomic differential models.** Sensitivity analysis comparing phosphosite-level differential estimates obtained using a condition-only *limma* model and a model additionally adjusted for sex. **(A–C)** Correlation of $\log_2$ fold-change estimates for hE2F4DN-treated h5xFAD versus untreated h5xFAD, untreated h5xFAD versus WT, and hE2F4DN-treated h5xFAD versus WT, respectively. **(D–F)** Corresponding comparison of $-\log_{10}(P)$ values for the same contrasts. Each point represents one quantified phosphosite-associated feature. Diagonal lines indicate identity between models, and Pearson correlation coefficients ($r$) are shown in each panel. The high concordance between models indicates that adjustment for sex had minimal influence on the overall treatment-associated phosphoproteomic estimates.

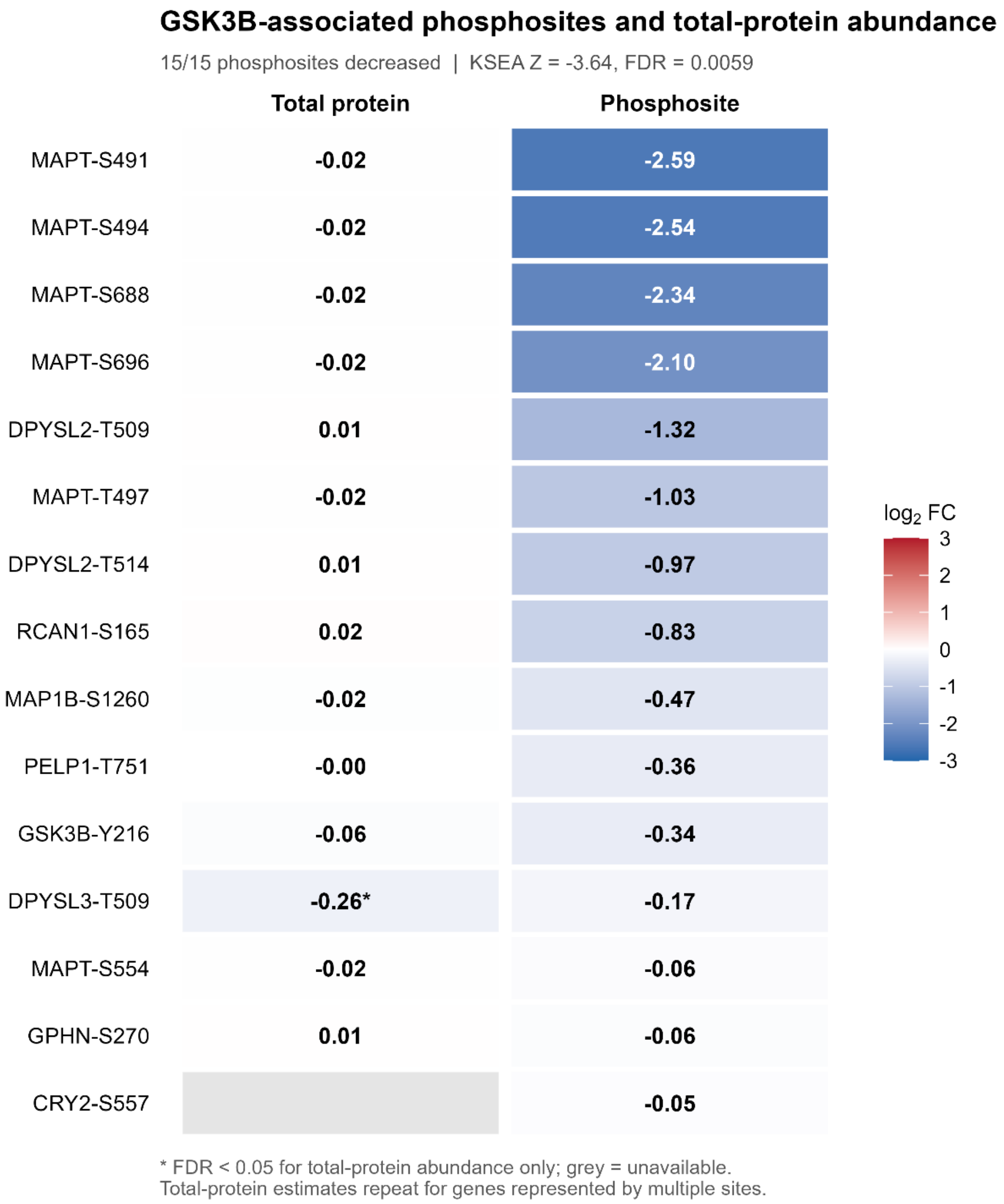


**FIGURE S10. GSK3B-associated phosphosite changes occur largely independently of total-protein abundance.** Treatment-associated $\log_2$ fold changes are shown for the 15 phosphosites contributing to the primary GSK3B kinase–substrate enrichment analysis (KSEA) signature and their corresponding total-protein measurements in hE2F4DN-treated versus untreated h5xFAD hippocampi. All 15 phosphosites exhibited negative treatment-associated changes (KSEA Z = −3.64, FDR = 0.0059). Corresponding total-protein measurements were available for 14 phosphosites, representing eight distinct genes, and generally showed substantially smaller changes. Notably, MAPT exhibited pronounced reductions across several phosphorylation sites ($\log_2$FC ranging from −2.59 to −0.06), despite minimal changes in total-protein abundance ($\log_2$FC = −0.02, FDR = 0.893). DPYSL3 was the only matched protein exhibiting an FDR-significant abundance change ($\log_2 FC = -0.26$). Protein abundance values are repeated for genes represented by multiple phosphosites. Cell colors represent $\log_2$ fold changes on a shared diverging scale (blue, decreased; red, increased), whereas grey indicates unavailable measurements. Asterisks indicate FDR $< 0.05$ for total-protein abundance only. These findings are consistent with treatment-associated phosphoregulation that is not primarily explained by changes in total-protein abundance.

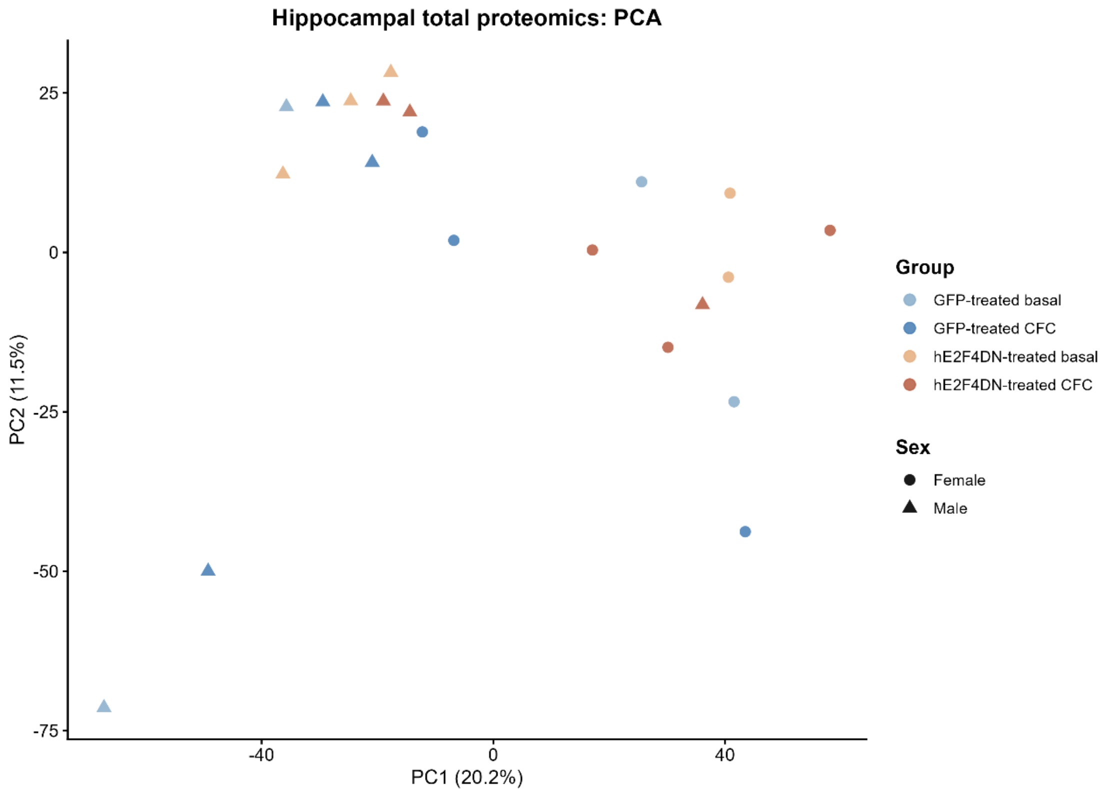


**FIGURE S11. Principal component analysis of hippocampal proteomic profiles following contextual fear conditioning.** Principal component analysis (PCA) of hippocampal total-protein abundance profiles from GFP-treated and hE2F4DN-treated h5xFAD mice under basal conditions or following contextual fear conditioning (CFC). The four groups comprised GFP-treated basal (n = 4), GFP-treated CFC (n = 6), hE2F4DN-treated basal (n = 5), and hE2F4DN-treated CFC (n = 6). Each point represents one biological sample; colors identify experimental groups, and point shapes distinguish females (circles) from males (triangles). Percentages on the axes indicate the variance explained by the first two principal components. PCA provides an overview of global proteomic variability across treatment, learning condition, and sex.

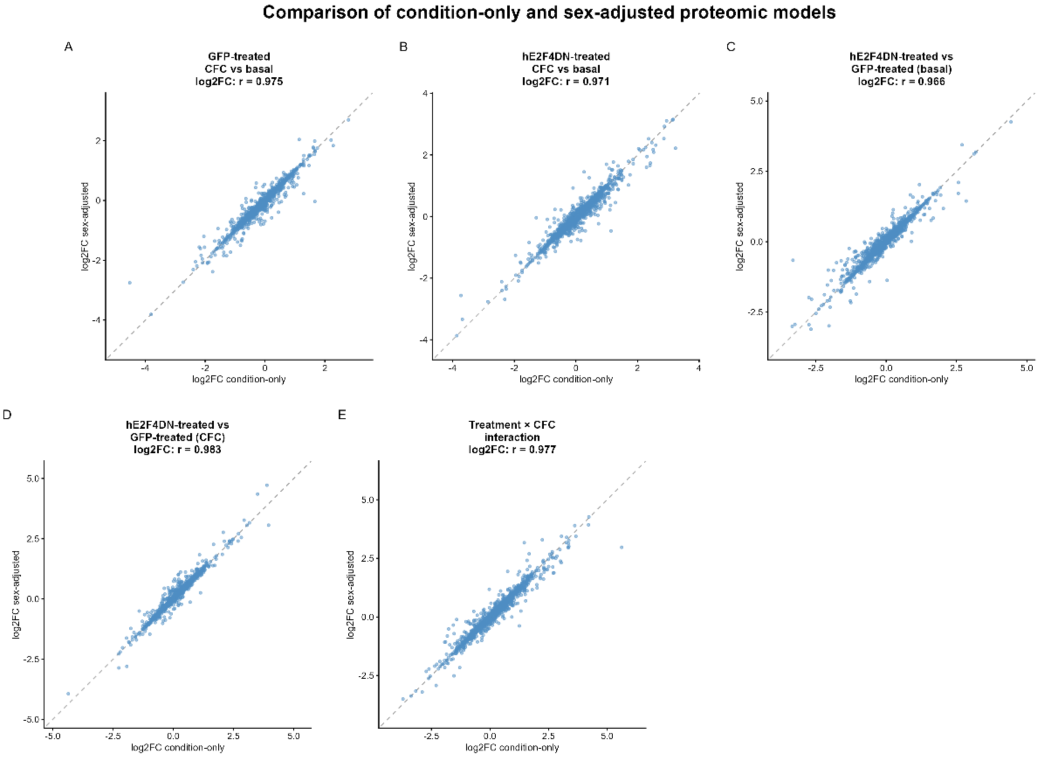


**FIGURE S12. Concordance of proteomic effect sizes between condition-only and sex-adjusted statistical models.** Comparison of protein-level $\log_2$ fold-change estimates from the primary condition-only limma model (x axis) and a sensitivity model additionally adjusted for sex (y axis). The five contrasts are (A) GFP-treated CFC versus basal; (B) hE2F4DN-treated CFC versus basal; (C) hE2F4DN-treated versus GFP-treated under basal conditions; (D) hE2F4DN-treated versus GFP-treated following CFC; and (E) the formal treatment-by-learning interaction. Each point represents one protein evaluated in both models. Dashed diagonal lines indicate identical effect-size estimates, and Pearson correlation coefficients (r) are shown in the corresponding panels. This comparison assesses the sensitivity of protein-level effect sizes to sex adjustment.

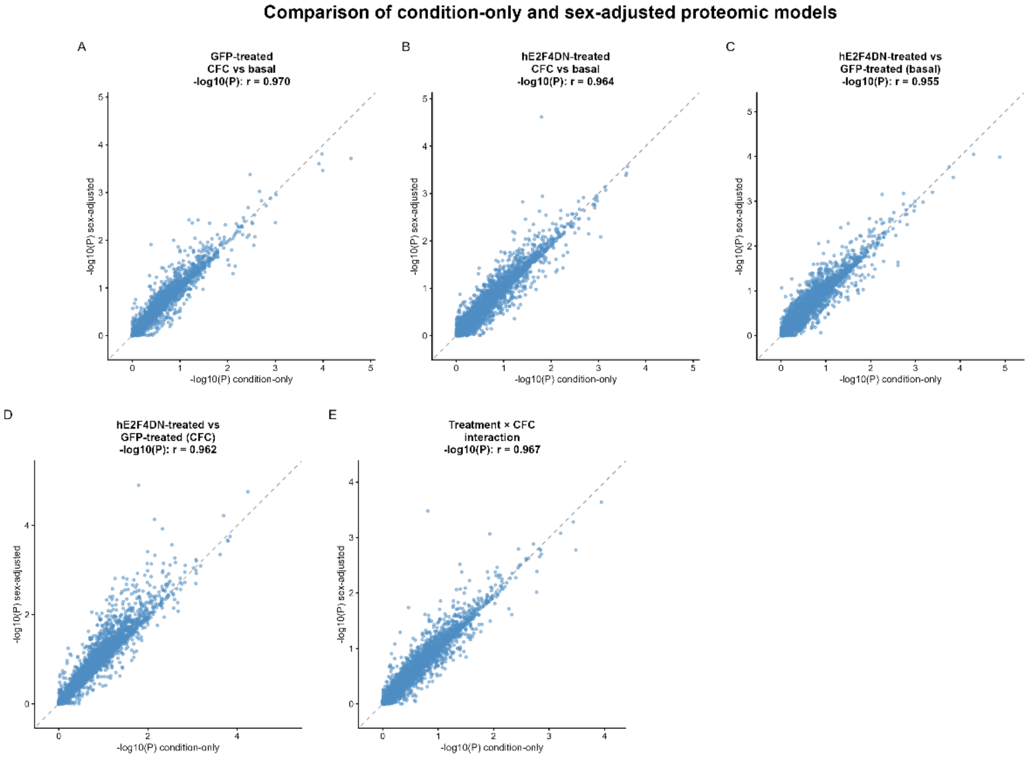


**FIGURE S13. Concordance of proteomic statistical significance between condition-only and sex-adjusted models.** Comparison of nominal protein-level statistical significance from the primary condition-only limma model (x axis) and the corresponding sex-adjusted sensitivity model (y axis), expressed as $-\log_{10}(P)$. Panels show (A) GFP-treated CFC versus basal; (B) hE2F4DN-treated CFC versus basal; (C) hE2F4DN-treated versus GFP-treated under basal conditions; (D) hE2F4DN-treated versus GFP-treated following CFC; and (E) the formal treatment-by-learning interaction. Each point represents one protein evaluated in both models. Dashed diagonal lines indicate identical values, and Pearson correlation coefficients (r) are displayed in each panel. Correlation of nominal P values assesses overall model concordance but does not imply identical significance classifications for individual proteins.

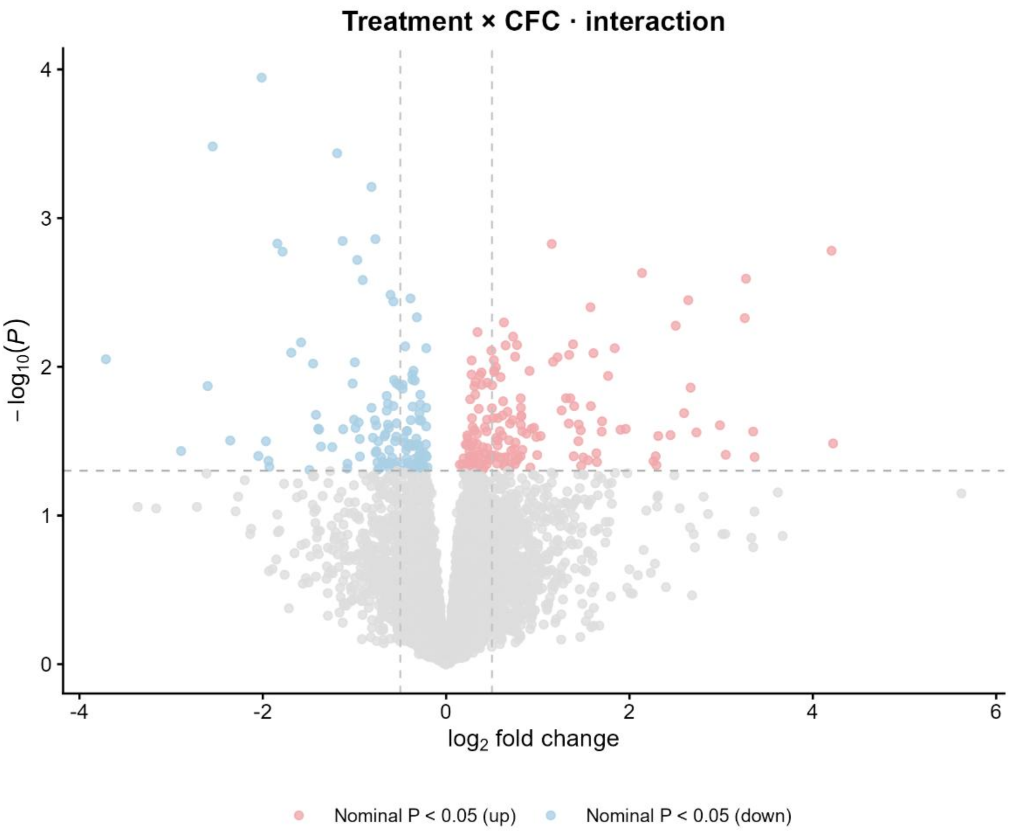


**FIGURE S14. Differential proteomic response identified by the treatment-by-learning interaction.** Volcano plot of the formal treatment-by-learning interaction in hippocampal total-protein abundance from GFP-treated and hE2F4DN-treated h5xFAD mice. The x axis represents the interaction $\log_2$ fold change, calculated as the CFC-associated response in hE2F4DN-treated mice minus the corresponding response in GFP-treated mice, and the y axis represents $-\log_{10}(P)$. Positive values indicate a more positive CFC-associated response in hE2F4DN-treated mice; negative values indicate a more negative response. Among 8,182 tested proteins, 296 showed nominal interaction evidence ($P < 0.05$), comprising 161 positive and 135 negative interactions; none reached FDR < 0.05. Light red and blue points indicate nominally positive and negative interactions, respectively, whereas grey points indicate proteins without nominal interaction evidence. Vertical lines at $|\log_2$ fold change$| = 0.5$ are reference guides only and were not used as significance thresholds. The corresponding comparison of learning-associated effect sizes is shown in Fig. 7B.

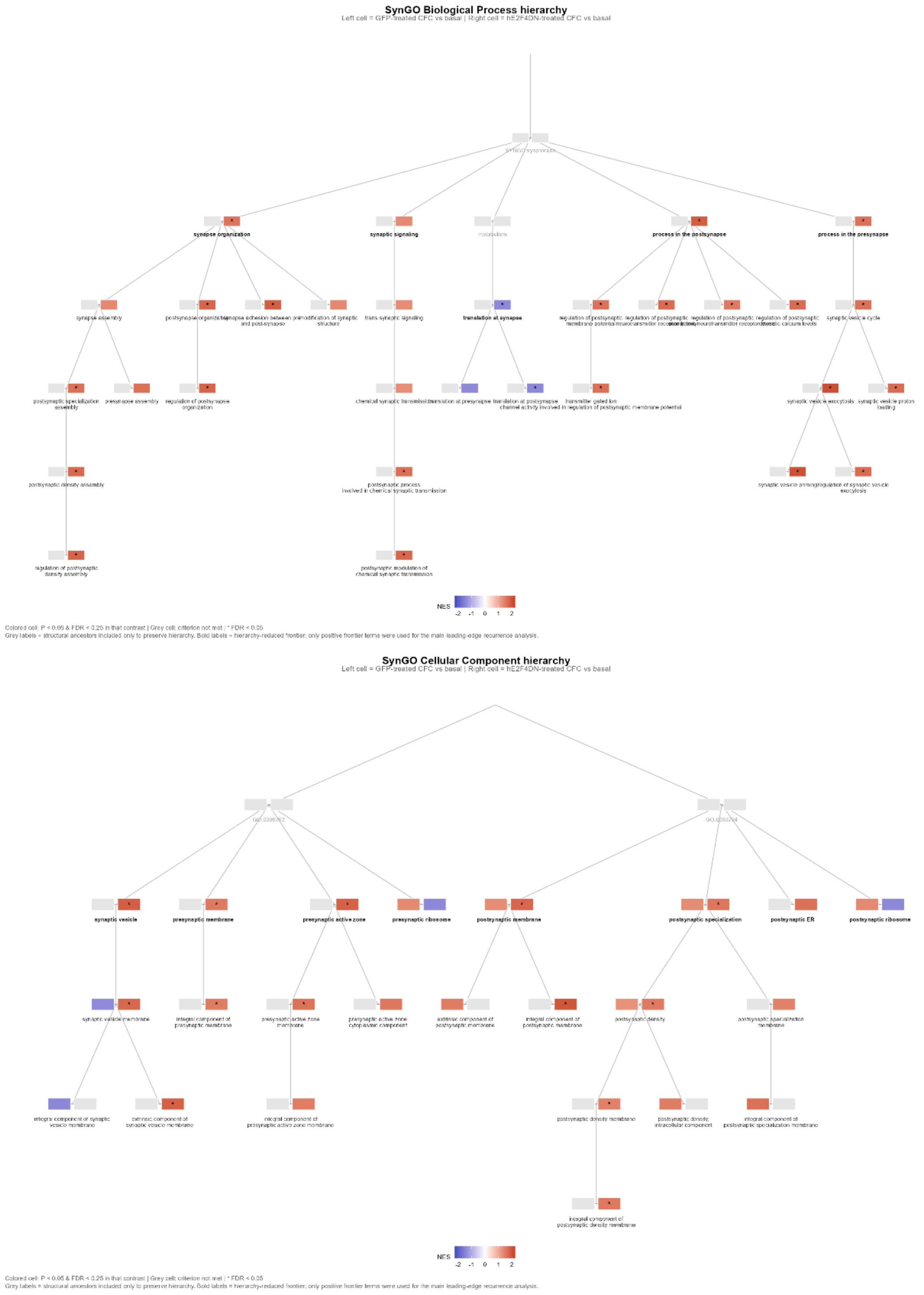


**FIGURE S15. Hierarchical organization of learning-associated SynGO enrichment and selection of representative synaptic pathways.** Hierarchical representation of SynGO terms enriched in the hippocampal proteomic response to contextual fear conditioning (CFC). The upper and lower panels depict the SynGO Biological Process (BP) and Cellular Component (CC) hierarchies, respectively. Terms meeting the exploratory criterion (nominal $P < 0.05$ and $FDR < 0.25$) in at least one learning contrast were included, together with structural ancestors required to preserve their SynGO parent–child relationships. A total of 30 BP and 23 CC terms met this criterion across the learning contrasts. Paired cells beside each term represent GFP-treated CFC versus basal (left) and hE2F4DN-treated CFC versus basal (right). Cell colors indicate normalized enrichment scores (NES; red, positive; blue, negative) only when the corresponding contrast meets the exploratory criterion; grey cells indicate that it was not met, and asterisks indicate $FDR < 0.05$. Grey labels identify structural ancestors retained solely for hierarchy visualization; bold labels identify the hierarchy-reduced frontier. The frontier contained five BP and eight CC terms, including four BP and six CC terms with positive enrichment following CFC in hE2F4DN-treated mice, which were used for the leading-edge recurrence analyses in Fig. 7D, E. Negatively enriched terms remain represented in the full hierarchy but were not included in positive leading-edge recurrence analyses.

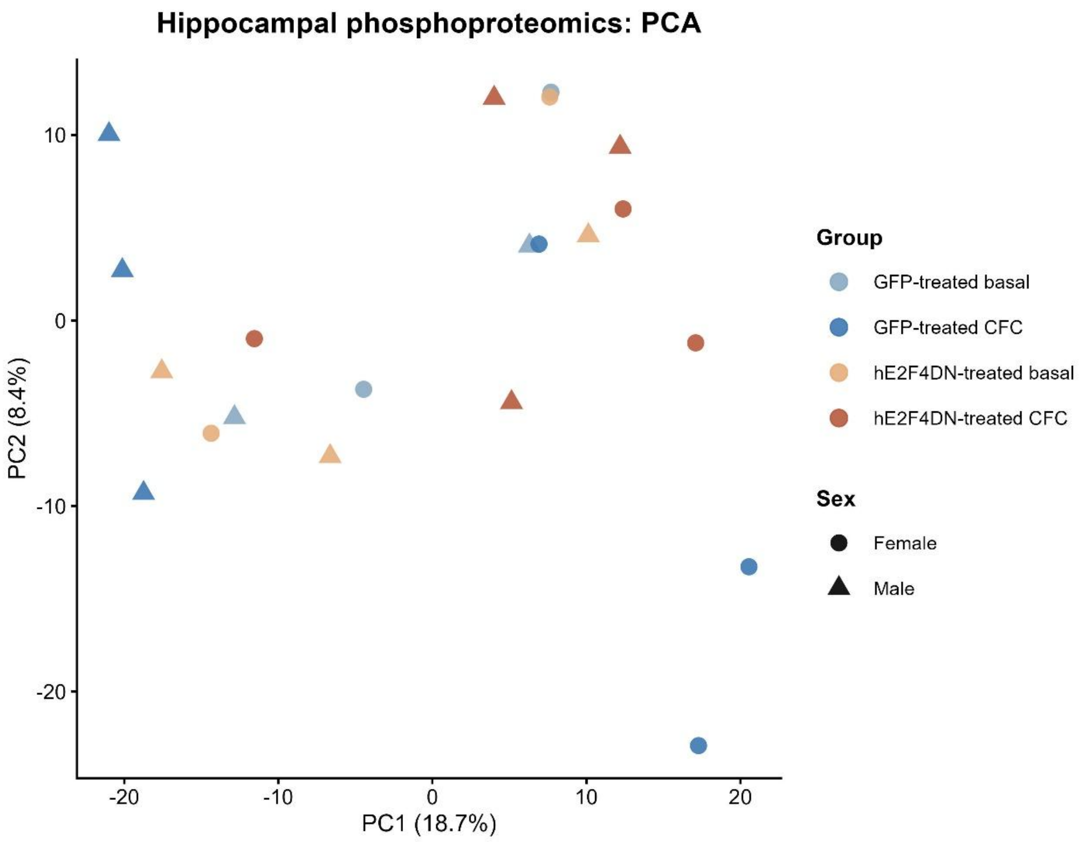


**FIGURE S16. Principal component analysis of hippocampal phosphoproteomic profiles following contextual fear conditioning.** Principal component analysis (PCA) of hippocampal phosphoproteomic profiles from GFP-treated and hE2F4DN-treated h5xFAD mice under basal conditions or following contextual fear conditioning (CFC). Each point represents an individual biological sample. Colors distinguish the four experimental groups (GFP-treated basal, GFP-treated CFC, hE2F4DN-treated basal and hE2F4DN-treated CFC), while point shapes indicate sex (circles, females; triangles, males). The first two principal components explain 18.7% and 8.4% of the total variance, respectively. Samples from the experimental groups show substantial overlap in the first two principal components, indicating that treatment and learning conditions do not produce a clear global separation within this projection.

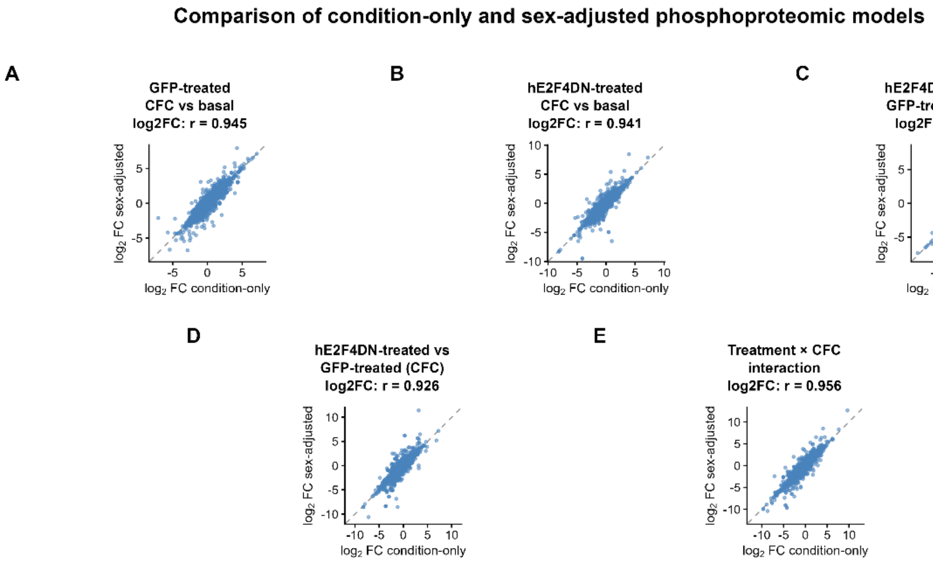


**FIGURE S17. Concordance of phosphoproteomic effect sizes between condition-only and sex-adjusted statistical models.** Comparison of phosphoproteomic $\log_2$ fold-change estimates obtained using the primary condition-only model and a sensitivity model additionally adjusted for sex. Each point represents a phosphofeature evaluated in both models. The x axis shows the $\log_2$FC estimated by the condition-only model, whereas the y axis shows the corresponding estimate from the sex-adjusted model. Five contrasts are presented: (A) GFP-treated CFC versus basal; (B) hE2F4DN-treated CFC versus basal; (C) hE2F4DN-treated versus GFP-treated under basal conditions; (D) hE2F4DN-treated versus GFP-treated following CFC; and (E) the formal treatment-by-learning interaction. Pearson correlation coefficients were 0.945, 0.941, 0.942, 0.926 and 0.956, respectively. Dashed diagonal lines indicate identical effect-size estimates between models. The high concordance across contrasts indicates that adjustment for sex had a limited influence on the overall estimated phosphoproteomic effect sizes.

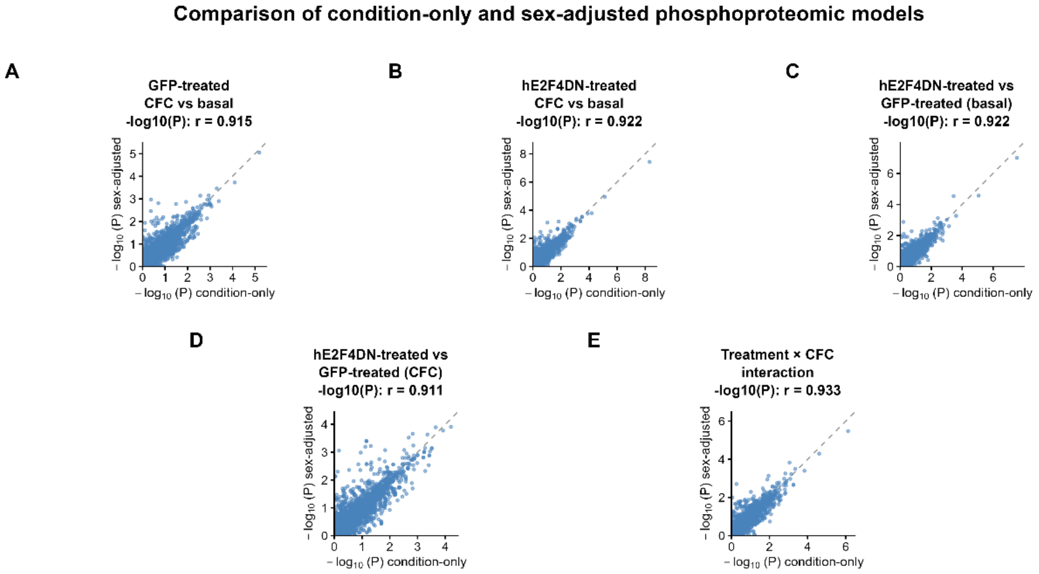


**FIGURE S18. Concordance of phosphoproteomic statistical significance between condition-only and sex-adjusted models.** Comparison of nominal statistical significance obtained using the primary condition-only phosphoproteomic model and the corresponding sex-adjusted sensitivity model. Each point represents a phosphofeature evaluated in both models, with nominal P values transformed as $-\log_{10}(P)$. The x axis shows results from the condition-only model and the y axis shows results from the sex-adjusted model. Panels correspond to (A) GFP-treated CFC versus basal; (B) hE2F4DN-treated CFC versus basal; (C) hE2F4DN-treated versus GFP-treated under basal conditions; (D) hE2F4DN-treated versus GFP-treated following CFC; and (E) the formal treatment-by-learning interaction. Pearson correlation coefficients were 0.915, 0.922, 0.922, 0.911 and 0.933, respectively. Dashed diagonal lines indicate identical nominal P values between models. The strong correlations support overall consistency of statistical evidence after adjustment for sex, although individual phosphofeatures may exhibit differences in significance.

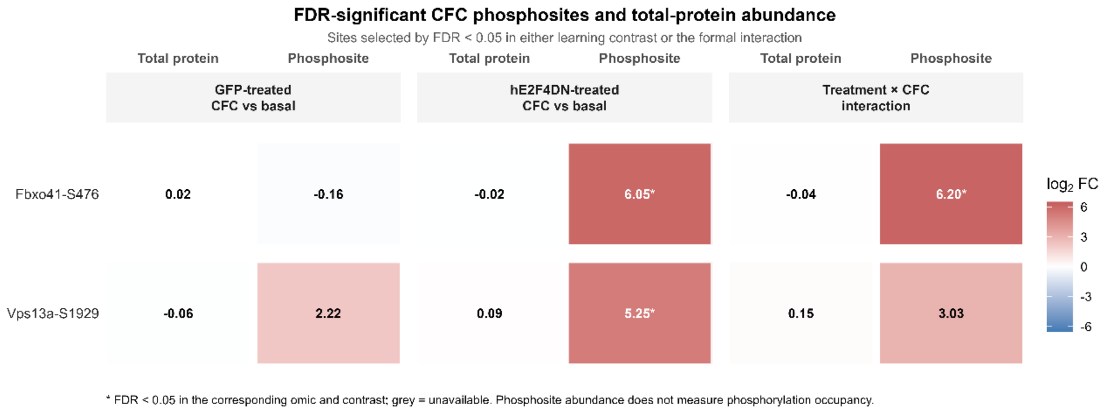


**Figure S19. FDR-significant learning-associated phosphosites exhibit pronounced changes without corresponding changes in total-protein abundance.** Comparison of phosphosite-level and total-protein abundance changes for FBXO41-S476 and VPS13A-S1929, the two phosphosites reaching FDR < 0.05 following CFC in hE2F4DN-treated mice. Treatment-specific learning responses (GFP-treated CFC versus basal and hE2F4DN-treated CFC versus basal) and the formal treatment-by-learning interaction are shown separately for phosphosite and total-protein measurements. FBXO41-S476 exhibited a marked increase following CFC in hE2F4DN-treated mice ($\log_2$FC = 6.05, FDR = 4.07 × $10^{-5}$) and a significant positive treatment-by-learning interaction ($\log_2$FC = 6.20, FDR = 0.0058). VPS13A-S1929 also increased following CFC in hE2F4DN-treated mice ($\log_2$FC = 5.25, FDR < 0.05), although its positive interaction did not survive multiple-testing correction ($\log_2$FC = 3.03). In contrast, total FBXO41 and VPS13A abundance showed only minor changes following CFC in hE2F4DN-treated mice ($\log_2$FC = −0.02 and 0.09, respectively), neither reaching FDR significance. Cell colors represent $\log_2$FC on a shared diverging scale (blue, decreased; red, increased). Asterisks indicate FDR < 0.05, whereas dots indicate nominal P < 0.05 without surviving multiple-testing correction, assessed independently for each omic layer and contrast. Grey cells indicate unavailable measurements. These findings indicate that the observed phosphosite responses are not accompanied by proportional changes in total-protein abundance; phosphosite abundance does not directly measure phosphorylation occupancy.

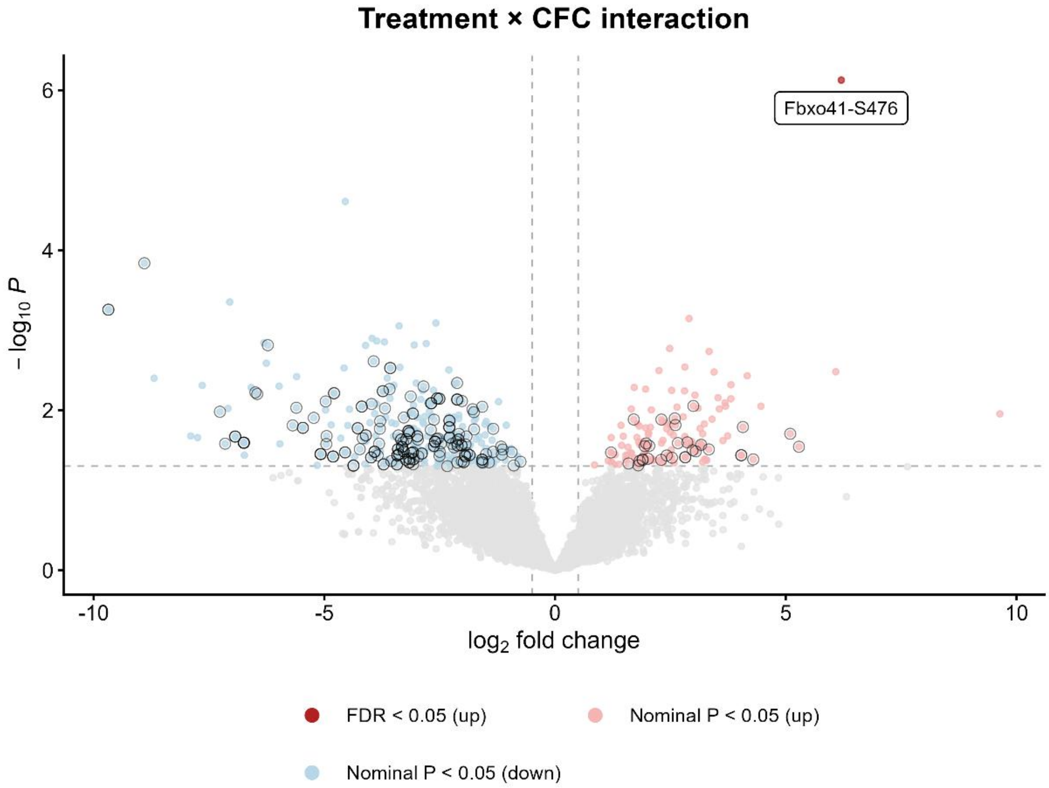


**FIGURE S20. Differential phosphoproteomic response to learning identified by the treatment-by-CFC interaction.** Volcano plot showing the formal treatment-by-learning interaction in hippocampal phosphoproteomic profiles from GFP-treated and hE2F4DN-treated h5xFAD mice. The x axis represents the interaction $\log_2$ fold change, calculated as the CFC-associated response in hE2F4DN-treated mice minus the corresponding response in GFP-treated mice, while the y axis indicates $-\log_{10}(P)$. Positive values indicate a more positive learning-associated response in hE2F4DN-treated mice, whereas negative values indicate a more negative response. Among 7,734 tested phosphofeatures, 473 showed nominal interaction evidence ($P < 0.05$), comprising 127 positive and 346 negative interactions. FBXO41-S476 was the only phosphosite reaching FDR < 0.05 (interaction $\log_2FC = 6.20$, FDR = 0.0058). Red and blue points indicate positive and negative nominal interactions, respectively, with FDR-significant phosphofeatures highlighted separately. Grey points indicate phosphofeatures without nominal interaction evidence.

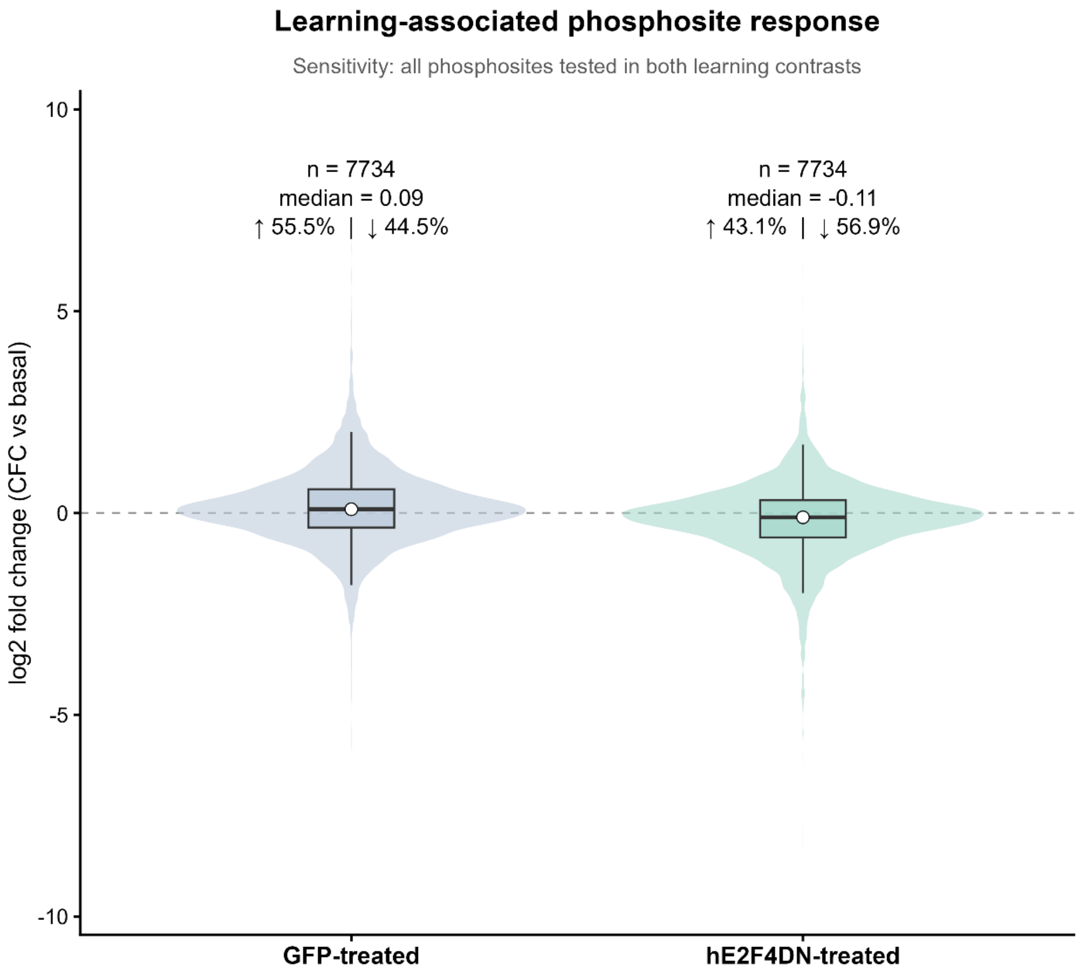


**FIGURE S21. Global directionality of learning-associated phosphoproteomic changes across all tested phosphofeatures.** Distribution of CFC-associated $log_2$ fold changes across all 7,734 phosphofeatures tested in both learning contrasts, independently of statistical significance. Violin plots show the distribution of $log_2FC$ estimates in GFP-treated and hE2F4DN-treated h5xFAD mice, with embedded boxplots summarizing the central distribution. GFP-treated mice exhibited a modest predominance of positive learning-associated changes (median $log_2FC = 0.09$; 55.5% positive and 44.5% negative), whereas hE2F4DN-treated mice showed a shift toward negative changes (median $log_2FC = -0.11$; 43.1% positive and 56.9% negative). The horizontal dashed line indicates no change ($log_2FC = 0$). This analysis extends the directionality assessment shown in Fig. 8C beyond nominally responsive phosphofeatures, demonstrating that the contrasting directional tendency is also observed across the complete set of commonly tested phosphofeatures.

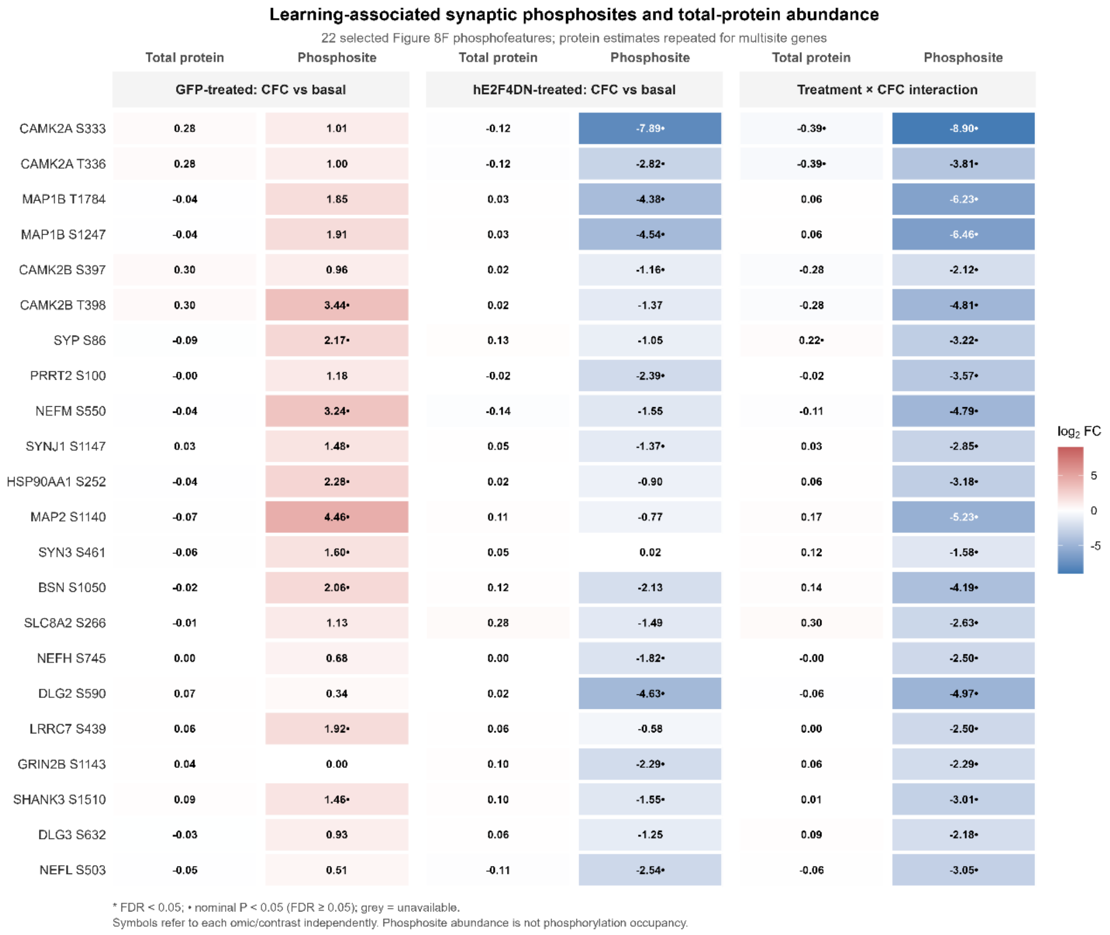


**Figure S22. Learning-associated synaptic phosphosite remodeling is largely uncoupled from changes in total-protein abundance.** Comparison of phosphosite-level changes and corresponding total-protein abundance measurements for the 22 representative phosphosites from 19 proteins included in Fig. 8F. $Log_2$ fold changes are shown for GFP-treated CFC versus basal, hE2F4DN-treated CFC versus basal, and the formal treatment-by-learning interaction, with separate columns for phosphosite and total-protein measurements. All 22 phosphosites were matched to corresponding total-protein measurements. Absolute phosphosite $log_2FC$ values exceeded those of the corresponding proteins for 21 of 22 sites (95.5%) in each learning contrast and for all 22 sites (100%) in the interaction analysis. These differences were generally not accompanied by equivalent changes in total-protein abundance, consistent with a predominantly phosphoregulatory response within the selected synaptic network. Protein-level estimates are repeated for genes represented by multiple phosphosites. Cell colors represent $log_2FC$ on a shared diverging scale (blue, decreased; red, increased). Asterisks indicate FDR < 0.05, whereas dots indicate nominal P < 0.05 without surviving multiple-testing correction. Grey indicates unavailable measurements. The comparisons are descriptive and do not represent direct measurements of phosphorylation occupancy or independent validation of the selected phosphosites.

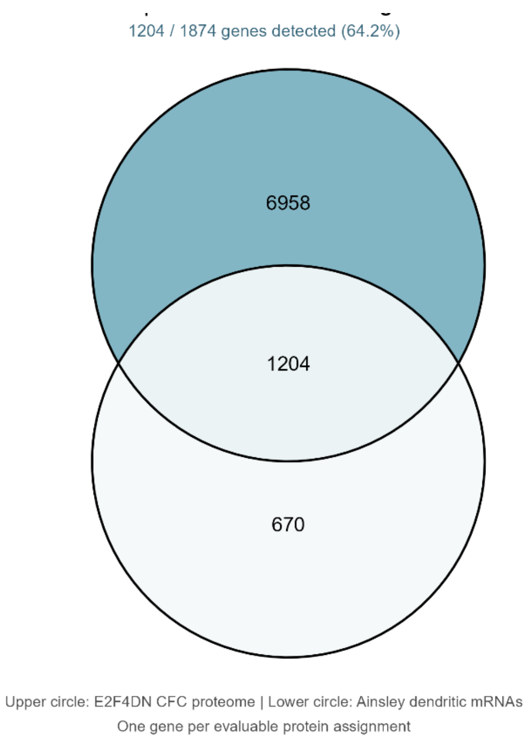


**FIGURE S23. Overlap between dendritic mRNA-associated genes and the hippocampal proteomic response to contextual fear conditioning.** Comparison of the dendritic mRNA-associated gene repertoire reported by Ainsley et al. (2014) with the hippocampal proteomic dataset obtained from E2F4DN-treated h5xFAD mice 60 min after contextual fear conditioning (CFC). Venn diagram showing the overlap between 1,874 genes associated with dendritic ribosome-bound mRNAs in Ainsley et al. (2014) and 8,162 unique genes represented by statistically evaluable proteins in the E2F4DN CFC proteomic contrast. A total of 1,204 dendritic mRNA-associated genes (64.2%) were represented in the proteomic dataset, whereas 670 were not detected among the evaluable gene assignments. Overlaps were calculated using unique gene symbols, with one representative protein assignment per gene. Complete gene-level overlap information and differential abundance statistics are provided in Supplementary Table S12.
